# InsectChange: Comment

**DOI:** 10.1101/2023.06.17.545310

**Authors:** Laurence Gaume, Marion Desquilbet

**Affiliations:** AMAP, University of Montpellier, CNRS, CIRAD, INRAE, IRD, Montpellier, France; Toulouse School of Economics, INRAE, University of Toulouse Capitole, Toulouse, France

**Keywords:** Insects, Terrestrial invertebrates, Freshwater invertebrates, Insect abundance, Insect decline, Time series meta-analysis, Methodological biases, Agriculture, Landcover

## Abstract

The InsectChange database (van Klink et al. 2021) underlying the meta-analysis by van Klink et al. (2020a) compiles worldwide time series of the abundance and biomass of invertebrates reported as insects and arachnids, as well as ecological data likely to have influenced the observed trends. On the basis of a comprehensive review of the original studies, we highlight numerous issues in this database, such as errors in insect counts, sampling biases, inclusion of noninsects driving assemblage trends, omission of drivers investigated in original studies and inaccurate assessment of local cropland cover. We show that in more than half of the original studies, the factors investigated were experimentally manipulated or were strong -often not natural- disturbances. These internal drivers created situations more frequently favouring an increase than a decrease in insects and were unlikely to be representative of habitat conditions worldwide. We demonstrate that when both groups were available in original freshwater studies, selecting all invertebrates rather than only insects led to an overestimation of the “insect” trend. We argue that the disparate and non-standardised units of measurement of insect density among studies may have detrimental consequences for users, as was the case for van Klink et al. (2020a, 2022) who log10(x+1)-transformed these heterogeneous data, compromising the comparison of temporal trends between datasets and the estimation of the overall trend. We show that geographical coordinates assigned by InsectChange to insect sampling areas are inadequate for the analysis of the local influence of agriculture, urbanisation and climate on insect change for 68% of the datasets. In terrestrial data, the local cropland cover is strongly overestimated, which may incorrectly dismiss agriculture as a driving force behind the decline in insects. Therefore, in its current state, this database enables the study of neither the temporal trends of insects worldwide nor their drivers. The supplementary information accompanying our paper presents in detail each problem identified and makes numerous suggestions that can be used as a basis for improvement.

## Introduction

Currently, experts agree that biodiversity is shrinking in the face of global changes of anthropogenic origin (IPBES, 2019). However, with respect to insects, which provide invaluable ecosystem services, the extent to which declines vary among insect groups and regions is still the subject of intensive investigation, with trend assessments hampered by lack of data and analytical weaknesses (Didham et al., 2020; Duchenne et al., 2022). There is also no consensus on the main drivers of insect changes, including land use (urbanisation/agriculture), climate change, pesticides, other pollution types and invasive species, mostly because these drivers are not easily disentangled or may act in synergy (Wagner et al., 2021; Outhwaite et al., 2022).

While many authors have warned of insect extinctions worldwide (Cardoso et al., 2020), van Klink et al. (2020a) added to the debate by estimating a smaller decline in the abundance of terrestrial insects than reported by previous authors, and further proposed that freshwater insects were increasing rather than decreasing. They found that increasing cropland cover was not associated with terrestrial insect decline and proposed that improved water quality was a driver of increasing abundance of insects in freshwaters. Yet their meta-analysis gave rise to comments by various authors regarding (1) their data selection and methodology (Desquilbet et al., 2020), which led to some corrections (van Klink et al., 2020b); (2) the limitations of abundance and biomass as sole indicators of insect trends, masking the possible replacement of sensitive species by stress-tolerant ones (Jähnig et al., 2021); and (3) the heterogeneity in temporal coverage, with a lack of old baselines (Duchenne et al., 2022).

The study of insect trends and their drivers addresses major environmental, societal, political and economic issues. This sensitive subject therefore requires, first and foremost, the utmost rigor in databases intended to serve as references. The InsectChange database (van Klink et al., 2021) underlying the analysis by van Klink et al. (2020a) includes time series of the abundance and biomass of invertebrates reported as insects and arachnids in terrestrial and freshwater realms worldwide, together with ecological data on anthropogenic changes likely to have influenced trends. We conducted a comprehensive and in-depth analysis of the relevance and accuracy of the InsectChange datasets by systematically reviewing the original studies. Our analysis highlights numerous limitations in the constitution of this database, the accumulation of which is likely to bias any assessment of insect change and drivers of change.

## 1 Different issues in the InsectChange database

The invertebrate taxa included in InsectChange are not only insects and arachnids as described in the title and abstract, but also entognaths (i.e., noninsect arthropods comprising springtails, diplurans and proturans), as indicated only in the keywords and appendices. Considering them within the scope of InsectChange and updating the analysis of Desquilbet et al. (2020) after the erratum by van Klink et al (2020b), we found that the sum of the remaining issues affected 161 of the 165 datasets. We found 553 issues, which belong to 17 types of problems pertaining to errors (153), inconsistencies (40), methodological issues (279) and information gaps (81), with 3.4 ± 1.6 problem types per dataset (Table 1, Figure 1a), as well as a methodological issue concerning the entire database. These multiple problems and the consequences they may have for the assessment of insect trends are detailed in Table 1. There were more problem types per dataset in the freshwater realm than in the terrestrial realm (Figure 1b, Appendix S1, *Problems.xlsx*), mainly because freshwater datasets were more affected than terrestrial datasets by problems related to the inclusion of invertebrates other than insects, the inclusion of studies with internal drivers and the assignment of inadequate geographic coordinates for local-scale analysis (Appendix 1, Figure 2b).

**Figure 1.**
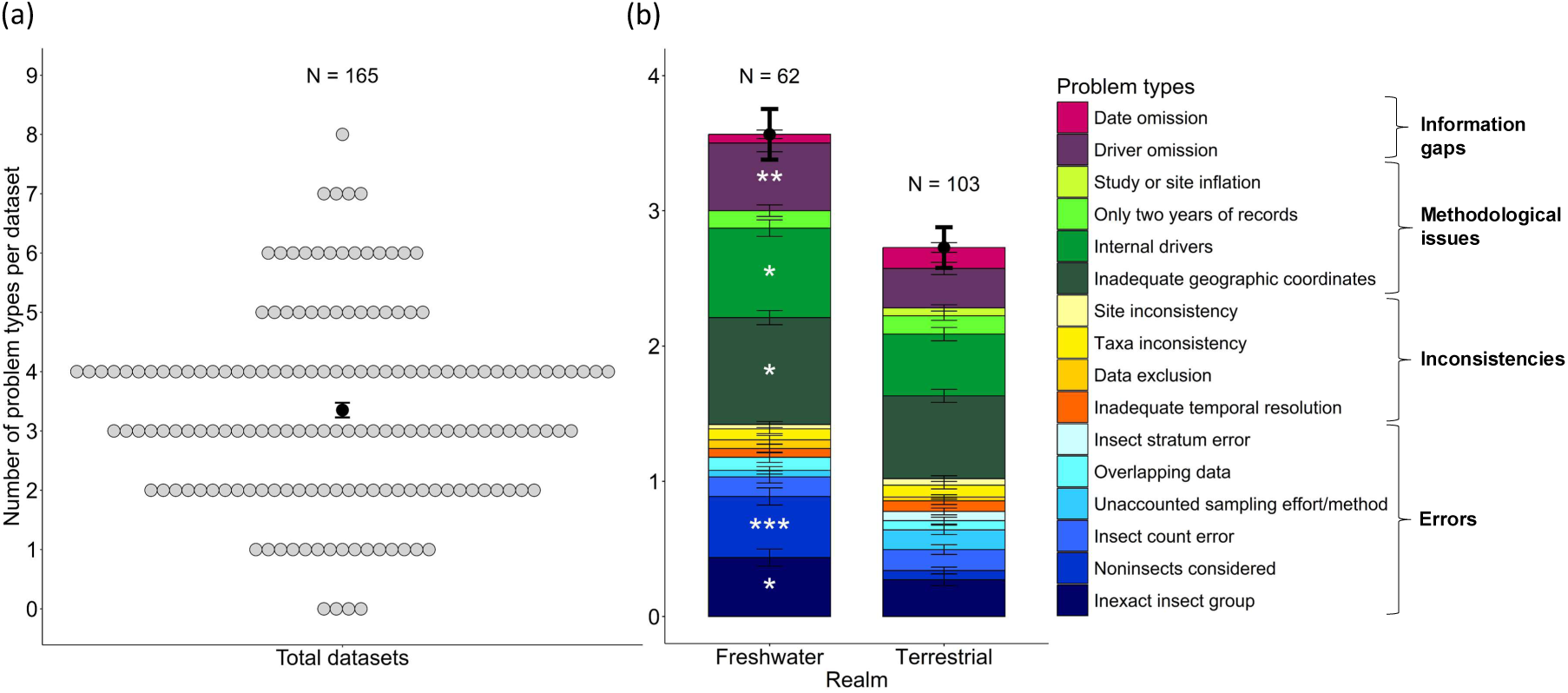
Distribution of the types of problems encountered in the InsectChange database (details in *Problems.xlsx*). (a) Mean (± SE) number of problem types per dataset and distribution of datasets according to the number of problem types. Each dot refers to a dataset; thus, the occurrence of *i* dots on the *y* line indicates that *i* datasets have *y* problem types. (b) Comparison of the mean number and distribution of problem types per dataset between freshwater and terrestrial realms. The problem type related to cropland cover, which was only assessed for the terrestrial realm, was not included in this comparative analysis, as well as the general problem of data heterogeneity. White stars were placed in the ‘Freshwater’ barplot when, on the basis of binary logistic regression, the problem type affected significantly more freshwater datasets than terrestrial datasets (Appendix 1). Terrestrial datasets were never significantly more affected by a given problem type than freshwater datasets were. *:0.01<P<0.05; **:0.001<P<0.01;***:P<0.001.

**Figure 2.**
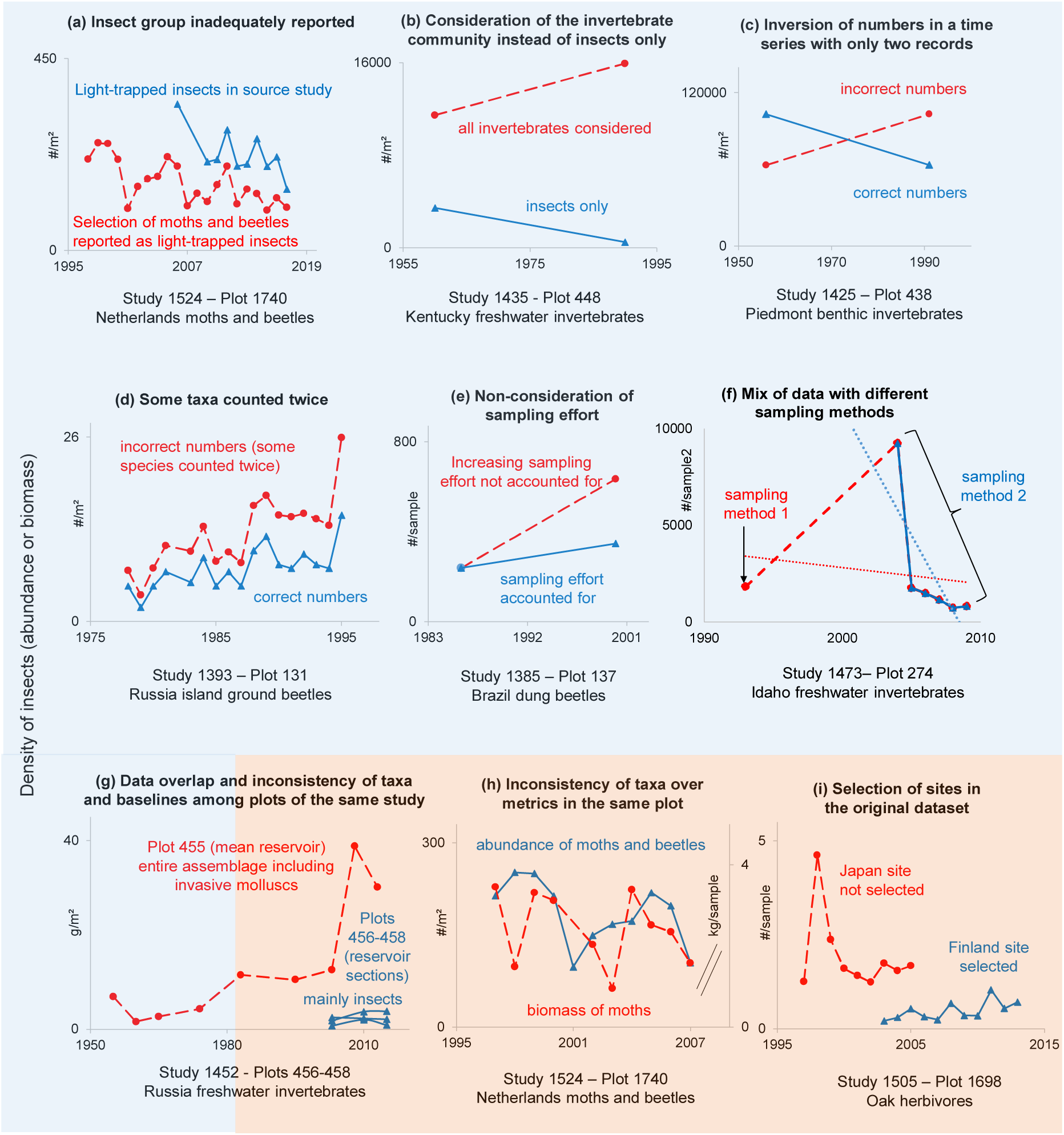
Examples of errors (blue blackground) and inconsistencies (orange background) in the selection of data, which affected the temporal trends in the original datasets (Appendix S1, *Problems.xlsx*, *Fig2and5.xlsx*). (a-g) Different types of errors; (g-i) Inconsistencies regarding taxa across plots (g) or metrics (h) and plot inclusion (i). Problematic insect dynamics are represented by red dashed lines, whereas nonproblematic or corrected insect dynamics are represented by solid blue lines.

**Table 1.**
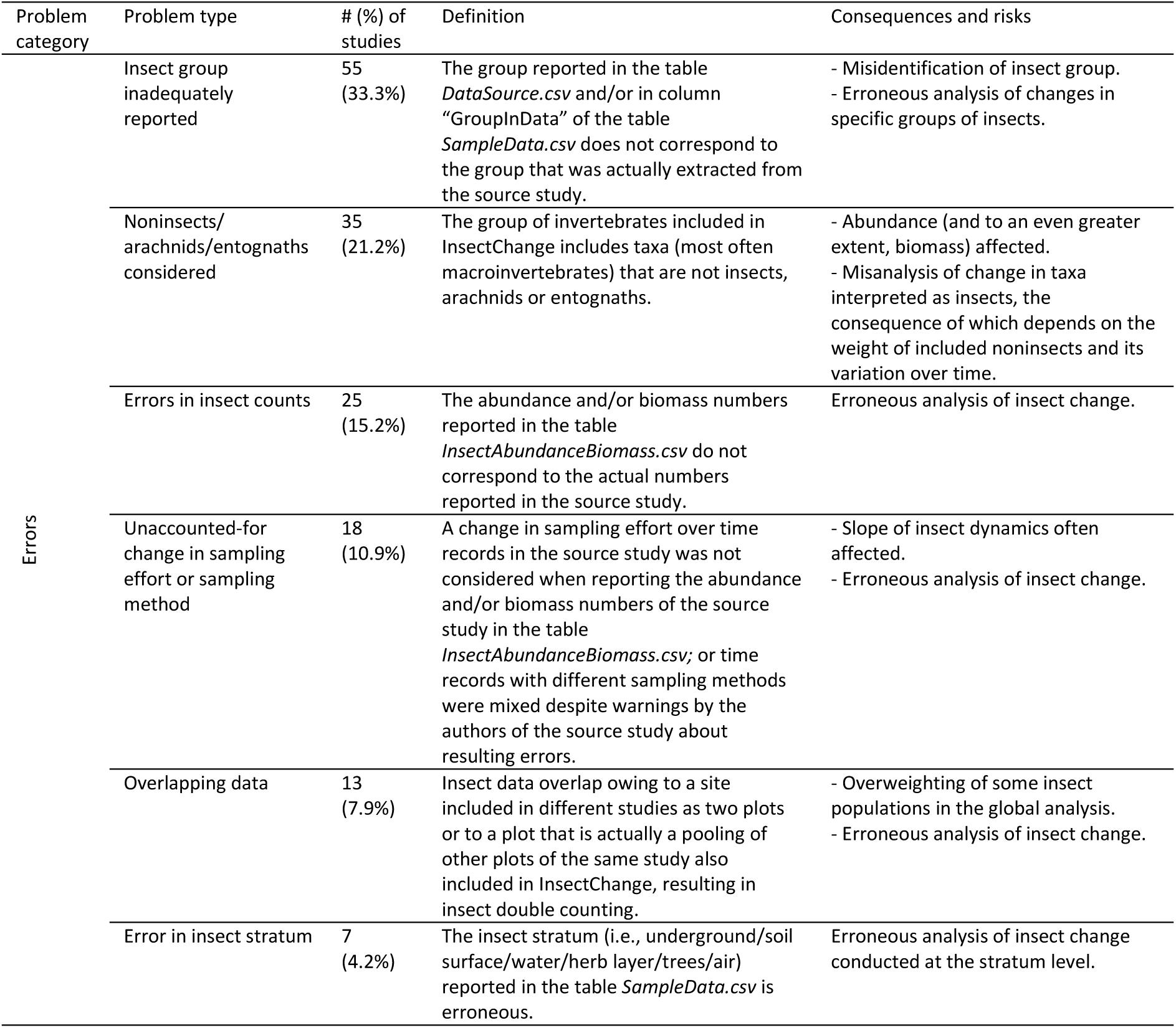
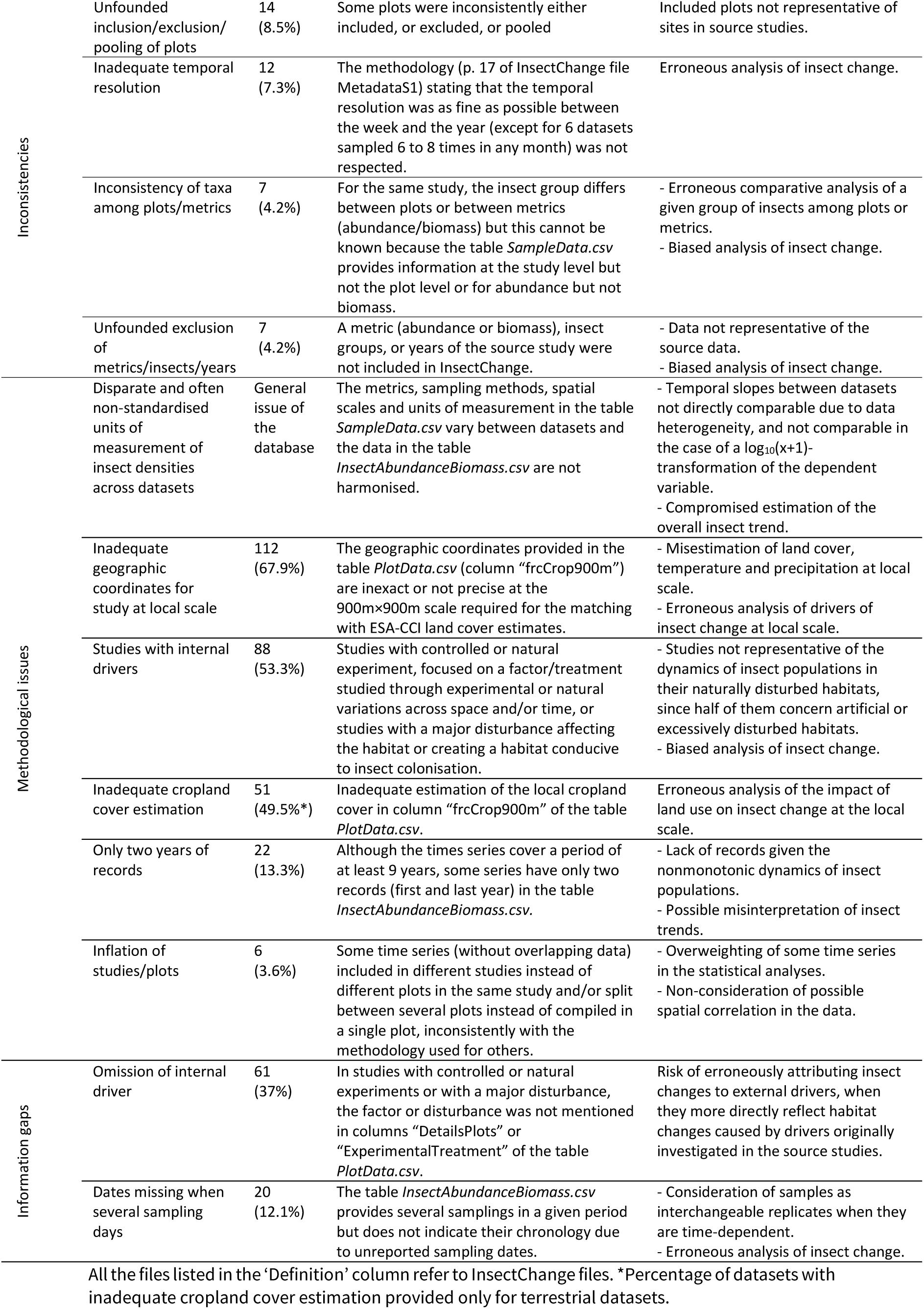
Description of the problem types, their frequencies and their possible impact on insect trend analysis.

### 1.1 Errors

Among the errors, the composition of the invertebrate group selected from the source datasets was misreported in 55 datasets, often because the authors of InsectChange neglected to specify that they had selected only certain taxa from the original community (e.g., Figure 2a). Moreover, 35 datasets considered taxa other than insects, arachnids or entognaths (hereafter collectively referred to as “insects” for brevity) and most often included the entire invertebrate assemblage instead of insects only, sometimes changing the insect trend of the original time series to the point of reversal (e.g., Figure 2b, details in Section 2). Insect counts were misreported from source studies in 25 datasets because of misinterpretation, calculation errors, the inversion of numbers, or species counted twice (e.g., Figures 2c and 2d). The stratum in which insects were sampled was misreported in 7 datasets, for example indicating that insects were sampled in the herb layer instead of trees. This may affect trend estimates by stratum, particularly those, such as trees, which are represented by only a few datasets (8 datasets for the tree stratum).

In addition, variation in sampling effort or method over time is a classic methodological bias in time series (Isaac & Pocock, 2015). It becomes an error when it is not noticed and considered by authors of meta-analyses, as was the case for 18 InsectChange datasets. This type of error may affect the insect trend of the included dataset, for example, when the number of sampling repetitions increased over time but the number of insects was added rather than averaged (e.g., Figure 2e), or when the authors of the source study specified that the sampling method changed between the first and subsequent records and did not themselves create a single time series from these two types of records, unlike the authors of InsectChange (Figure 2f).

Finally, some datasets or plots had overlapping data for all or part of the time periods (13 datasets), resulting in double counting, either because different datasets included the same plots or because a plot in a given dataset was actually a pooling of others from the same dataset. This leads to overweighting some insect populations in the global analysis. In 9 datasets, the exact same insects were counted twice. For example, InsectChange Study 1452, which is illustrated in Figure 2g, examined the change in biomass of the invertebrate assemblage after the creation of the Kama Reservoir in Russia. InsectChange Plots 456, 457 and 458 corresponding to the upper, central and dam sections of the reservoir, respectively, include data from 2003 to 2015 mainly for insects, and Plot 455, corresponding to the average sampling in the three sections of the reservoir, includes data from 1955 to 2013 on the entire zoobenthic assemblage. From 2003 to 2013, insect data from Plot 455 therefore overlap with invertebrate data from Plots 456, 457 and 458, with the same insects counted twice. In two other InsectChange datasets, data overlapped because one study reported the abundance dynamics of ant nests and the other, centred on the same ants, reported the abundance dynamics of the ants themselves. The last two datasets included the dynamics of grasshoppers in the soil stratum of the same three sites, obtained by visual counting for one dataset and collection in pitfall traps for the other. These different cases of overlapping data may affect the analysis of overall insect trends.

### 1.2 Inconsistencies

There were also a number of inconsistencies. In 7 datasets, there were inconsistencies of taxa between plots of the same dataset (e.g., Figure 2g, shows time series considering the entire assemblage of invertebrates including invasive molluscs in a plot and insects and crustaceans in other plots) or between metrics in the same plot (abundance or biomass; e.g, Figure 2h shows time series of the abundance of moths and beetles and the biomass of moths only). Because InsectChange does not indicate the insect group at the plot or metric level, users cannot identify these inconsistencies, which may lead to errors in comparative analyses of insect groups between different plots or metrics or in the estimation of the global trend of a particular insect group. Moreover, unfounded inclusion, exclusion (e.g., Figure 2i) or pooling of original sites affected 14 datasets, with potential consequences for insect trend analysis. In 7 datasets, there were also unfounded exclusions of data regarding a metric, some insect groups, or some time records. Furthermore, 12 datasets had temporal resolutions that did not match the resolutions of the original datasets or those stated in the InsectChange metadata (Table 1, Appendix S1, *Problems.xlsx*). While the temporal resolution should have been “as fine as possible between the week and the year” (“except for 6 datasets sampled 6 to 8 times in any month”), the data were sometimes averaged at the yearly level even though data for months were available and there were sometimes more than 8 records per month. All these inconsistencies in data selection mean that the data are not representative of the source studies.

### 1.3 Methodological issues and information gaps

With respect to methodological issues and information gaps, the inclusion of studies with internal drivers, i.e., experimental conditions or major disturbances, and the frequent omission of information on these drivers are the focus of Section 3; the adequacy of geographic coordinates and the estimation of the local cropland cover are the focus of Section 4.

In addition, a major methodological issue affecting the whole database is that the comparability of temporal trends between datasets is compromised by the heterogeneity of insect measurements, contrary to what is stated in InsectChange (e.g., p. 24 of MetaDataS1 file). Harmonisation of measurements was either not possible, due to variations in metrics (abundance/biomass), sampling methods and spatial scales between datasets, or was possible using standardisation for a given metric and sampling method, but was not achieved. Abundance and biomass units were thus not harmonised in the table *InsectAbundanceBiomass.csv* of InsectChange and were not clearly and/or systematically indicated in the table *SampleData.csv* of InsectChange. For example, abundance could be expressed as the number of individuals per m², per 0.1 m², or per sample, and biomass in g/m², mg/m² or g/sample. In many instances, the source and units for biomass data were not provided, notably when both abundance and biomass were available in the dataset (Appendix S1). This means that users often need to return to the source data to determine the data units. This problem may thus have detrimental consequences for users of the database who wish to estimate insect temporal trends. These detrimental consequences depend on whether the dependent variable in the model is transformed before analysis or not and on the type of transformation. For example, to avoid the log of 0 and reduce the high discrepancies in insect counts, van Klink et al. (2020a) and van Klink et al. (2022) used a log_10_(x+1)-transformation of these nonharmonised data, adding 1 to each abundance or biomass number before log-transformation. However, whereas a log_10_(x)-transformation gives the same regression slope over time whether the dependent variable in a time series is expressed, for example, in mg/m² or g/m², a log_10_(x+1)-transformation gives different regression slopes. This raises a problem in the case of a meta-analysis focused on trend estimation where the dependent variable is expressed in different units of measurement. More precisely, in the case of a log_10_-transformation of x, the slope of x (e.g., biomass in our case) with respect to t (time in our case), i.e., (log(x_2_)-log(x_1_))/(t_2_-t_1_) = log(x_2_/x_1_)/(t_2_-t_1_), expresses the relative variation of x over t (e.g., +10%/year) and not the absolute variation (e.g., + 2.7 g/year). With a log(x+1)-transformation, if x is numerically close to 0, log(x+1) is comparable to x and the slope is almost an absolute variation. If x is numerically high, log(x+1) is comparable to log(x) and the slope is almost a relative variation. Therefore, the interpretation of log(x+1) changes with the magnitude order of x. This issue is especially problematic in InsectChange, where magnitude orders of nonstandardised data vary between datasets from 10^-16^ to 10^6^. For these reasons, this methodological issue compromises the comparison of temporal trends between datasets or groups of datasets and the overall insect trend estimation, and calls into question the results obtained from the InsectChange database.

Besides, 22 datasets had only two years of records (20 of these 22 had two records per plot), whereas the nonmonotonic dynamics of insect populations require more records (Roubik, 2001; Didham et al., 2020). It is well known that time series without sufficient records lack statistical power and are potentially misleading (Roubik, 2001; White, 2018). This problem of only two years of records involves 13.3% of the studies (n = 165) and is not randomly distributed across continents. For example, it affects a quarter of the datasets and plots in Asia (4 of 16 datasets and 22 of 84 plots), suggesting that insect trend assessment for this continent on the basis of InsectChange data is likely biased. This methodological issue of only two years of records is particularly problematic when combined with other types of problems, such as considering the whole assemblage of invertebrates instead of just insects (Figure 2a), reversing insect counts (Figure 2c) or failing to correct for a change in sampling effort (Figure 2e) because, as a result, insect trends may be radically altered compared with those of the original datasets.

Another methodological issue is the inflation of datasets and/or sites (without overlapping data) compared with the original studies (6 datasets). For example, site inflation may result from splitting some sites of the original datasets into several InsectChange plots separated by only a few metres. This leads to overweighting of these datasets in the statistical analyses. This may also lead uninformed users to carry out statistical analyses without accounting for possible spatial correlation.

Finally, in 20 datasets, dates were not indicated when there were successive samplings per month. This information gap may lead users to consider samples as interchangeable replicates when they are time dependent.

## 2 Focus on the problematic inclusion of clams, snails, worms and shrimp in freshwater data

In the freshwater realm, 80% (19 of 24) of the biomass datasets and 40% (21 of 54) of the abundance datasets included invertebrates other than insects. This issue concerned 28 distinct datasets, 24 of which included the entire freshwater invertebrate assemblage (Figure 3, Table S1 in Appendix S2). The great majority of these 24 datasets included data on worms, molluscs and crustaceans, and taxa such as Oligochaeta, Hirudinea, Turbellaria and Amphipoda, which are often indicative of poor water quality (Enns et al., 2023) (Table S2 in Appendix S2). The inclusion of these datasets is not consistent with the purpose of the database because the dynamics of insects cannot be inferred from those of entire invertebrate assemblages. This is illustrated in Figure 3a, which presents examples from datasets in the study in which insect and invertebrate assemblages have contrasted trajectories (top), and in which proliferating invasive molluscs drive the trend of the invertebrate assemblage (bottom, *FreshwaterNonInsects.xlsx*).

In addition, we calculated that on average insects made up 48.7% ± 31.9% of the entire assemblage in the 13 datasets (48 plots) with information on all or part of the time records (Figure 3b, *FreshwaterNonInsects.xlsx*). The insect share in the assemblage also highly varied over time (Figure S1 in Appendix S2), with a coefficient of variation averaging 36.5% ± 21.6% in the nine datasets (42 plots) where information was available for more than one time record. Therefore, considering noninsects in the assemblage can considerably alter the temporal trend of abundance or biomass at the scale of the source study. Out of the 53 plots of 15 datasets with information on invertebrates driving the trend of the entire assemblage, noninsects (invasive molluscs, opportunistic oligochaetes and/or amphipods, etc.) were found to drive the assemblage trend in almost half of the plots (25), affecting two-thirds of the datasets (10)

**Figure 3.**
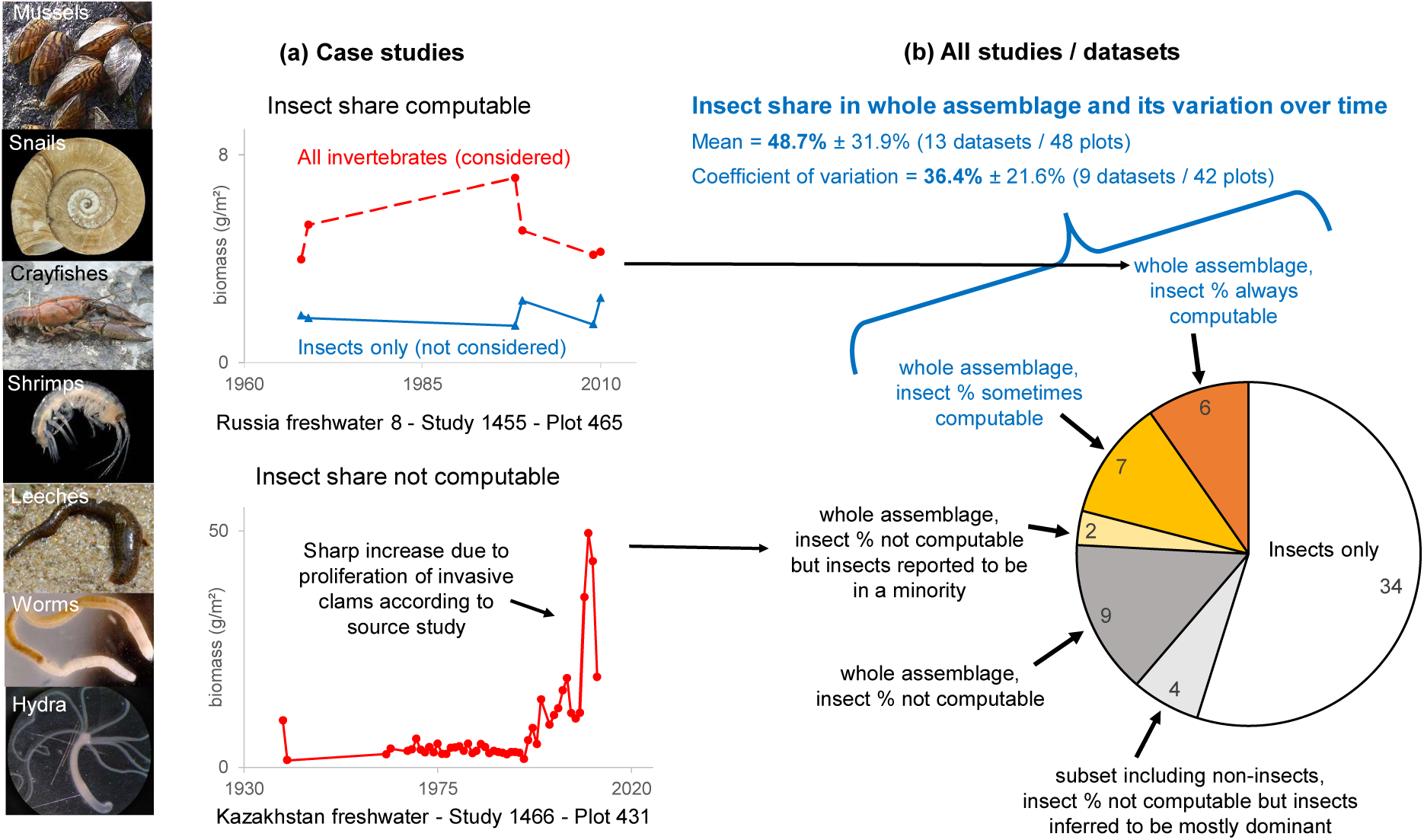
Freshwater time-series including noninsects, while the insect share was often low and variable over time (Appendix S2, *FreshwaterNonInsects.xlsx*). (a) Case studies 1455 and 1466 (Appendix S1) illustrating (top) contrasted trajectories of the entire assemblage and insects only, and (bottom) invasive noninsects driving the trend. (b) The 62 freshwater datasets, 28 including noninsects (24 of which included the entire invertebrate assemblage). The percentage of insects, which could be extracted from 13 of these, averaged 48.7% with a 36.4% coefficient of variation over time. (Appendix S1, *Problems.xlsx*). Insect % not computable: the insect data were not available from the original time series focused on the whole invertebrate assemblage. Insect % ‘sometimes’ or ‘always’ computable: it was possible to extract the percentage of insects for some or all records of the time series, respectively. Insects inferred to be mostly dominant: the percentage of chironomids, which are part of the insects in these InsectChange-selected data subsets of original datasets, could be calculated for each time record and was most frequently well over 50% (Table S1 in Appendix S2). Credits for the photographs are detailed in Table S3 in Appendix S2.

To visualise the differences between trends between total invertebrates and insects at the plot level, we extracted the estimates of regression slopes for each plot and the two assemblage types when possible (21 plots, 7 datasets, three from the USA, three from Russia, one from Italy). To this end, we first converted data units into international units. Unlike van Klink et al. (2020a), we did not log_10_(x+1)-transform the data (Section 1.3) but log_10_(x)-transformed them, which was possible given the absence of zeros in the abundance and biomass counts in this data subset. We ran as many analyses of covariance as there were plots, each on log_10_(x)-transformed insect densities, using the time covariate expressed in years, the assemblage type as a factor and the interaction between them as the explanatory variables (*FreshwaterNonInsects.xlsx*). The temporal trend estimates were very different for insects and total invertebrates for most plots and were even reversed for one-third of them (7 out of 21, Figure 4a).

**Figure 4.**
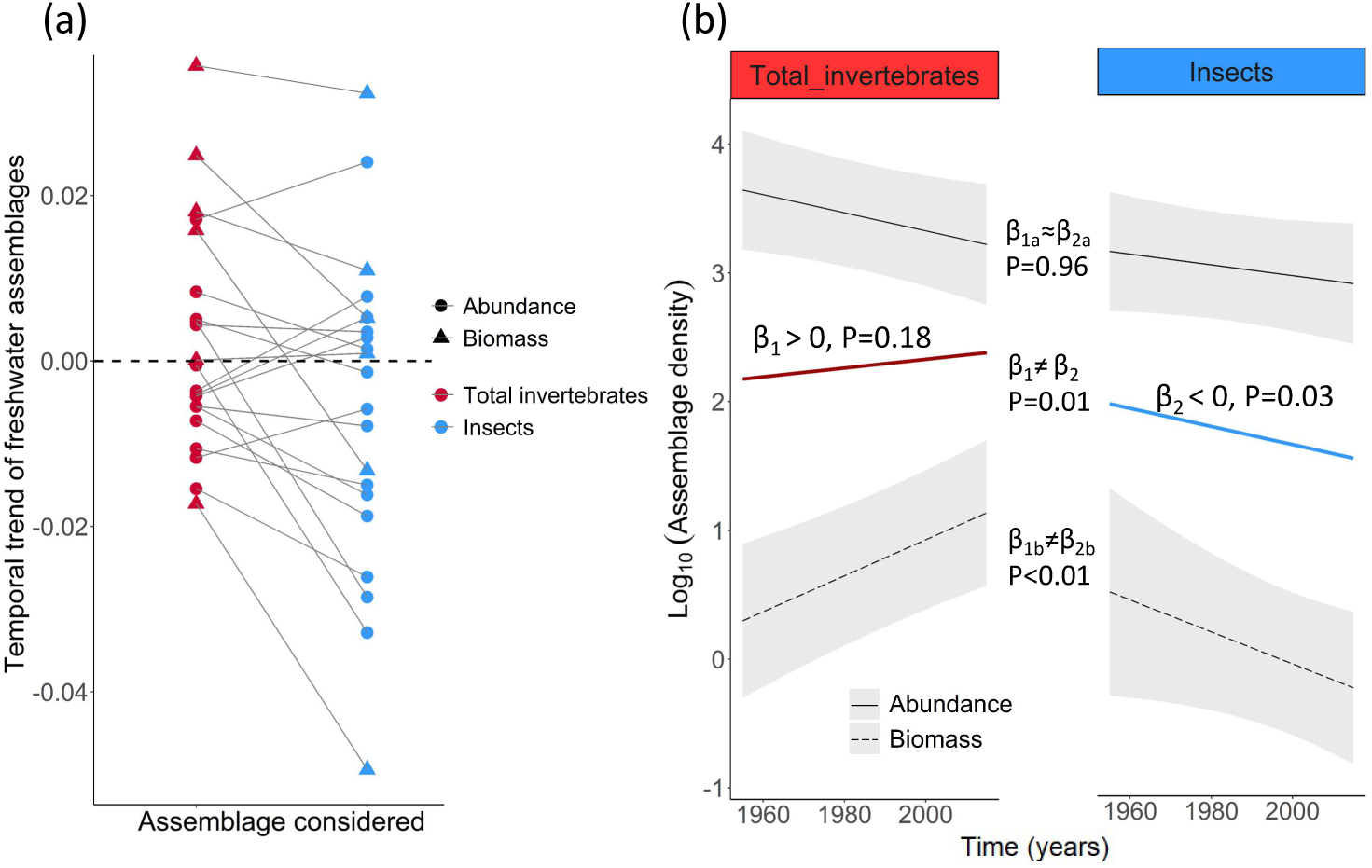
Comparison of temporal trends (in the log_10_ space) of invertebrate abundance or biomass between all invertebrates and insects only, for freshwater time series on whole assemblages and for which the insect share was always computable. (a) Comparison at the plot level. The trends were very different for insects and total invertebrates for most plots, and reversed for 7 of the 21 plots. (b) Comparison at a larger scale (mean estimate for the subset of 7 datasets and 21 plots). The results of the mixed linear model showing significantly different trend estimates between the two assemblage types for biomass data, for which a positive trend (β_1b_) was observed when all invertebrates were considered and a negative trend (β_2b_) when only insects were considered. The overall (abundance and biomass combined) trend estimate was positive (β_1_) but not significantly different from zero for all invertebrates, and significantly negative (β_2_) for insects (Appendix 2).

To test whether this problem affects the trend on a wider scale than that of the plot, we compared the mean trends between insects and all invertebrates in this data subset, which was composed of 21 plots from seven datasets, four with abundance data, and three with biomass data. To this end, we performed a mixed linear model on the log_10_-transformed insect densities. The fixed variables were the same variables as those used previously, and in addition the metric as a factor and the associated second and third-order interactions. We chose datasets and plots nested within datasets as random variables, considering them as independent and identically distributed, such as van Klink et al. (2020a). We found that the temporal trends differed significantly depending on the assemblage considered (significant interaction with time, Appendix 2), especially for biomass data (significant third order interaction). While the temporal trend was negative for abundance and did not differ significantly between assemblage types (P=0.96), for biomass, the temporal trend was positive for total invertebrates (P=0.0003) but was significantly lower (P<0.01) for insects for which it tended to be negative (P=0.066) (Figure 4b, Appendix 2, *FreshwaterNonInsects.xlsx*). This first demonstrates that abundance and biomass trends can be very different, particularly when considering entire assemblages that can include large-size and invasive taxa such as certain molluscs. This further demonstrates that, on a wider scale than that examined in the study, considering entire invertebrate assemblages rather than only insects can lead to significant overestimation of the temporal trend (also see results of the post hoc tests for trend comparisons for abundance and biomass combined with a significant trend difference (P=0.01) between the two assemblage types, Appendix 2).

## 3 Inclusion of datasets specifically designed to study particular, often experimentally manipulated, factors of insect changes

A major limitation of the InsectChange database is that users are inclined to erroneously attribute insect changes to possible anthropogenic drivers, such as changes in cropland cover, urban cover or climate, included in InsectChange after extraction from external databases, when they more directly reflect habitat changes caused by internal drivers, i.e., factors of insect changes specifically investigated in the original studies. Indeed, 88 out of the 165 datasets were extracted from controlled or natural experiments or from strongly disturbed contexts, and the factors that were originally investigated or major disturbances affecting the results were not mentioned in 69% of these 88 datasets (Table 1, Appendix S3, *Problems.xlsx*). Among these 88 datasets, 14 concerned controlled experiments testing the effect of one or several treatments in different plots (Figure 5a) and 53 concerned natural experiments (Diamond, 1983) investigating the effect of a natural disturbance by comparing insect abundance in more or less disturbed plots (Figure 5b) or before and after the disturbance in a plot (Figure 5c). In these experimental datasets, only control plots, only experimental plots or both types of plots were inconsistently included in InsectChange. In 21 observational datasets of the 88, a strong disturbance affected insect trends (Figure 5d).

**Figure 5.**
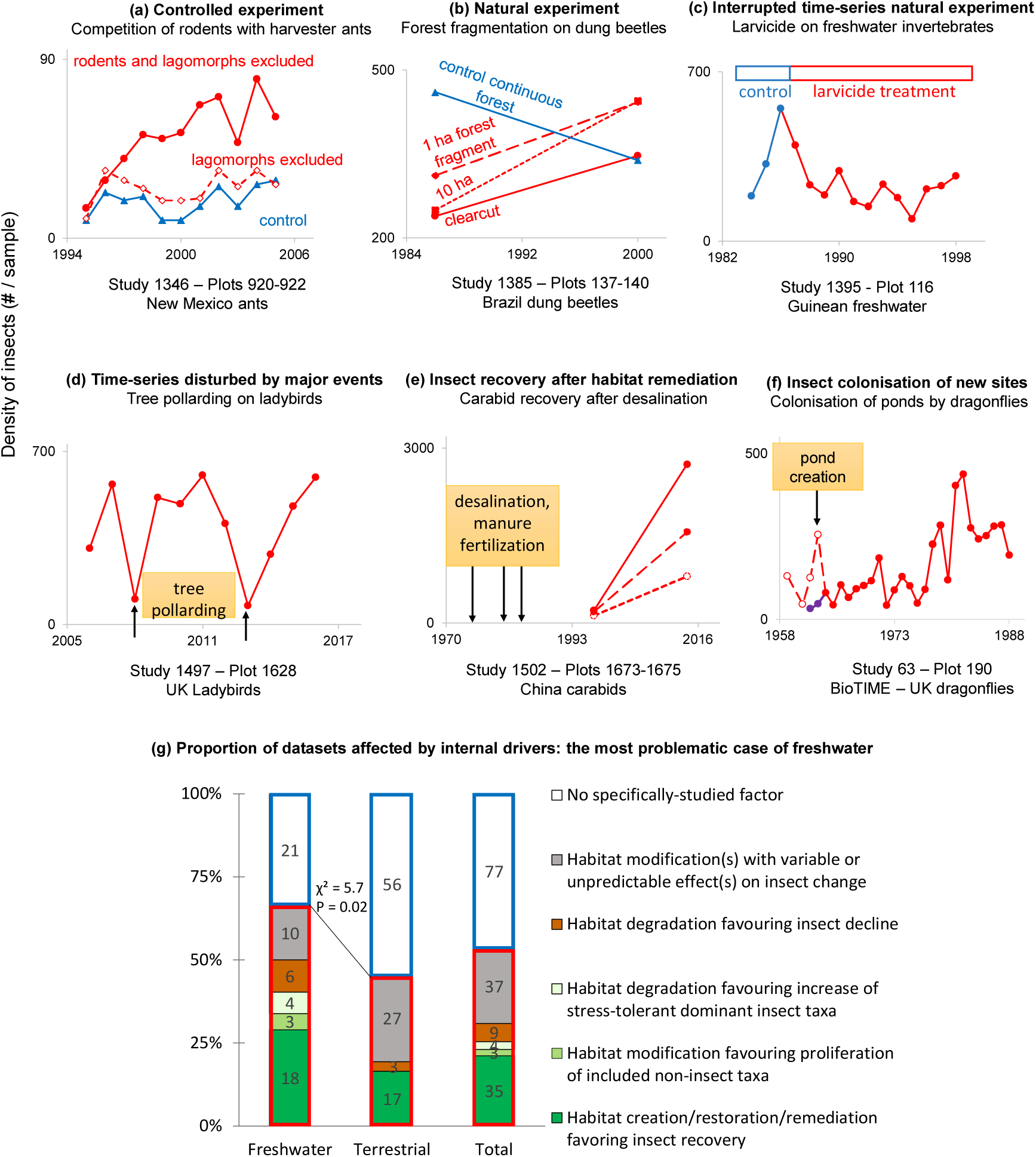
Inclusion of datasets specifically designed to study particular factors of insect change (internal drivers), the combination of which is unlikely to be representative of habitat conditions worldwide (Appendix S3, *Fig2and5.xlsx*). Examples of (a-c) controlled or natural experiments and (d-f) datasets with major disturbances; in (f), dashed and purple curves represent erroneous and actual data, respectively. (g) Comparison of the proportions of datasets affected by internal drivers between freshwater and terrestrial realms, showing the particularly problematic case of freshwater. It is also worth noting the frequency of situations favouring an increase in insects compared with their decrease. Red frame: datasets with internal drivers; blue frame: datasets without internal drivers; green: increases in insects favoured; brown: decreases in insects favoured.

Among these 88 datasets, the internal factors could be expected to have positive effects, for example, the effects of cessation of harmful activities, remediation measures (e.g., Figure 5e), active restoration or creation of new habitats such as nesting sites, reservoirs or ponds (e.g., Figure 5f) that favour insect recovery or colonisation. The studied factors could also be negative, such as severe drought, fire or pesticide application, creating deleterious conditions for insects at the beginning, middle or end of the observation period, followed by recovery, the timing of which strongly influences insect trends (Appendix S3). An increase in invertebrate abundance or biomass after a negative factor of pollution was paradoxically expected in six freshwater studies (Figure 5g, Appendix S3), because only or mostly stress-tolerant chironomids were considered or proliferating noninsects were included such as oligochaetes, opportunists in waters affected by eutrophication (Rosa et al., 2014), or invasive amphipods.

Two-thirds (41 of 62) of the freshwater datasets were affected by internal drivers, a proportion significantly higher than that (one half: 47 of 103) of the terrestrial datasets (χ² = 5.7, P = 0.02, Figure 5g left and middle, Appendix S3). Among these two-thirds, internal drivers were found to create situations that favour an increase in the number of insects in 61% of the cases (25 freshwater datasets in green in Figure 5g, left). Considering all the freshwater and terrestrial datasets impacted by specific drivers (Figure 5g, right), there were five times more situations favouring an increase in insects (42 datasets in green) than those favouring a decline in insects (nine datasets in brown).

This analysis raises the question of whether the data included in InsectChange are representative of habitat conditions and associated insect abundances worldwide, particularly in freshwater. While the selection of data according to specific and consistent criteria is a necessary condition for a meta-analysis to lead to robust conclusions (Englund et al., 1999), it was not met in InsectChange. The inclusion of time series with specific experimental designs to address ecological questions with differing purposes and expectations raises three issues for meta-analyses and other syntheses carried out using this database. (1) Such inclusion does not fit the definition of a meta-analysis as “a set of statistical methods for combining the magnitudes of the outcomes (effect sizes) across different datasets addressing **the same research question**” (Koricheva et al., 2013); (2) it implies that plots within datasets are not independently and identically distributed, which is not indicated in InsectChange; and (3) it introduces the problem of the “false baseline effect” (Didham et al., 2020), i.e., any nonrandom bias towards an above-average or a below-average starting point in a time series comparison, with a subsequent bias in the overall trend estimation. Therefore, because of these often artificial situations, which lead to below-average starting points much more frequently than above-average starting points, the insect trends obtained from InsectChange data (van Klink et al., 2020a) for freshwater and terrestrial realms are most likely overestimated.

How could data selection be improved in InsectChange? First, to reach more robust and meaningful conclusions, the best way to proceed would be to select more homogeneous datasets enabling testing of a single clear hypothesis, or alternatively to control for heterogeneity among studies with statistical analyses that take these differences into account with predictor variables. For controlled experiments, it would be relevant to consider only control sites. For other datasets, care should be taken to ensure the representativeness of situations and drivers in terms of sites with or without disturbance and in terms of timings of disturbance, and disturbance types could be weighted according to their frequency (Cardinale et al., 2018). Maps of human impacts on ecosystems, for example, could guide the choice of data and/or their weighting (Gonzalez et al., 2016).

In any case, users are exposed to the risk of misinterpreting trend drivers if they use InsectChange data, i.e., insect changes and local indicators of anthropogenic changes extracted from external databases, without knowledge of the factors originally investigated in the source studies.

## 4 Methodological issues resulting in a strong overestimation of the local cropland cover

Finally, we found a strong overestimation of local cropland, a possible driver included in InsectChange, by matching study plots to land covers provided in the European Space Agency Climate Change Initiative (ESA CCI) database (ESA, 2017) via the geographic coordinates that were either provided in the source studies or inferred by the authors of InsectChange. According to our analysis of terrestrial plots, this problem arises because (1) the geographic coordinates assigned to InsectChange plots are often inadequate for indicating sampling locations and (2) the interpretation of satellite images to determine land cover at actual sampling locations is often imperfect.

### 4.1 Assignment of inadequate geographic coordinates for local analysis

The local scale around each plot is defined in InsectChange as the area of 900 m × 900 m centred on the 300 m × 300 m ESA-CCI cell encompassing the geographic coordinates assigned to the plot and including the eight surrounding ESA-CCI cells. This area is used to estimate cropland or urban cover at the local scale. The adequacy of these local-scale indicators hinges on the premise that, for each plot, the geographic coordinates assigned to the plot in InsectChange are precise enough to point to the insect sampling area, and that this sampling area is included in a 900 m × 900 m square (hereafter referred to as a “local-scale square”, Figure 6a). However, this was not the case for almost a quarter of the terrestrial plots (233 out of the 985 plots). This methodological issue affected 63 of the 103 terrestrial datasets included in InsectChange.

**Figure 6.**
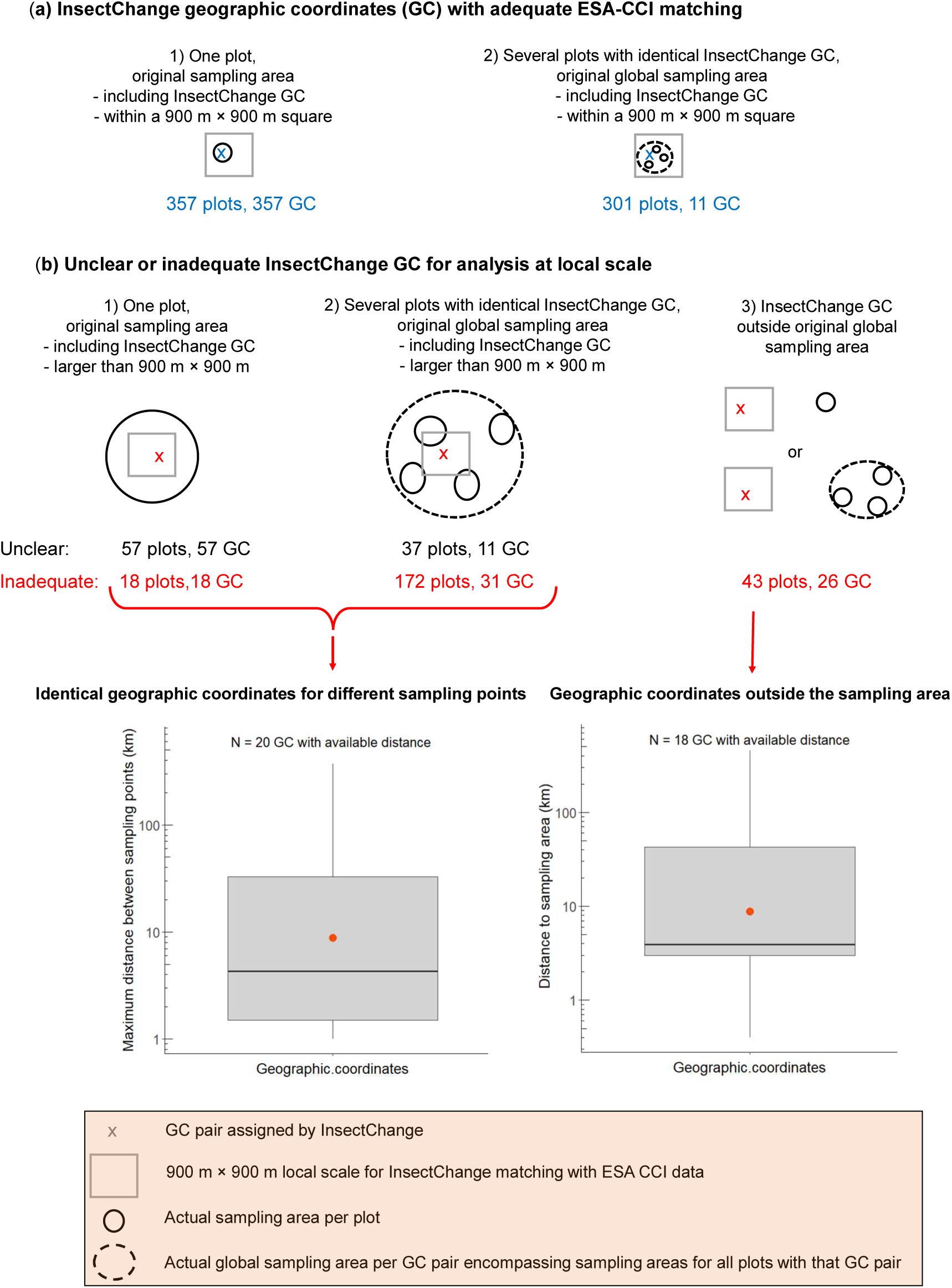
Inadequate assignment of geographic coordinates (GCs) for local analysis: the case of terrestrial plots. (a) adequate InsectChange GCs, (b) unclear or inadequate GCs, and, for inadequate GCs, boxplots (including the mean in red) of the maximum distance among sampling points in case of identical InsectChange GCs for different sampling points (left) and of the distance to the sampling area when the GCs were outside the sampling area (right).

We assessed the matching with ESA-CCI as adequate when the actual sampling area was at the location indicated by InsectChange geographic coordinates and small enough to be encompassed in a local-scale square (Figure 6a). By this criterion, matching was adequate for 658 out of 985 terrestrial plots of InsectChange. Among these, 357 were assigned different geographic coordinates. Each of these geographic coordinates adequately indicated the actual sampling area, which was adequately encompassed in a local-scale square (Figure 6a1). The remaining 301 plots shared geographic coordinates with others, with a total of 11 distinct geographic coordinates assigned by InsectChange. Each of these 11 coordinates adequately pointed to a zone included in the global sampling area of the original study comprising the sampling areas of different plots assigned a unique pair of geographic coordinates. This global sampling area was itself small enough to be encompassed in a local-scale square (Figure 6a2).

By contrast, we assessed the matching with ESA-CCI as unclear for 94 plots and inadequate for 233 terrestrial plots (Figure 6b), as detailed in our supplementary table *CroplandCover.xlxs*. We assessed the matching as unclear either when the sampling area was a butterfly transect and we found no information on the size of this transect, or when several plots shared the identical geographic coordinates and we found no information on their precise sampling areas.

Among the plots with an inadequate matching, the actual sampling area was larger than a local-scale square for 190 terrestrial plots, either because a unique InsectChange plot aggregated data from actual sampling points more than 900 m distant from each other (18 plots; Figure 6b1) or because several InsectChange plots with the same assigned geographic coordinates corresponded to actual sampling areas more than 900 m distant from each other (172 plots; Figure 6b2). Both cases contradicted the statement in InsectChange that data on the cropland cover were extracted “at and surrounding the sampling sites”, which implicitly assumes that for each plot, the sampling area was fully encompassed in a local-scale square. Matching with an external database is thus not appropriate, as it provides land cover information either for only part of the sampling area or for an unsampled area. When information was available, the maximum distance between sampling points in a sampling area varied from 1 to 370 km, as shown on the left boxplot in Figure 6b. For example, the 370 km distance is related to Study 1470, where InsectChange extracted a mean hymenopteran time series from Belarus in a unique plot and assigned it a location in Belarus where no sampling actually occurred. The information from the source study gave the names of the areas where the insects were sampled, allowing calculation of the distances between sampling points, which ranged up to approximately 370 km. Therefore, the local-scale indicators calculated around the geographic coordinates assigned to this unique “plot” are not meaningful for informing on the local conditions around the actual sampling points.

For the remaining 43 plots with inadequate matching, the geographic coordinates were included in a local-scale square that was outside the sampling area (Figure 6b3). When information was available, the distance between the InsectChange geographic coordinates and the actual sampling area varied from 400 m to 450 km, as shown in the right boxplot on Figure 6b. For example, from the columns PlotName, Location and DetailsPlot in the table *PlotData.csv* of InsectChange, Plots 1656 (Study 1266) and 1670 (Study 1006) represent the Cairngorms site of the UK Environmental Change Network, but were inadequately assigned the geographic coordinates of the 450 km-distant Yr Wyddfa/Snowdon site. Other sources of inadequacy are detailed in our supplementary table *CroplandCover.xlxs*. They include the use of different geographic coordinates than those provided in the source study, an error when transforming geographic coordinates to the decimal format, the inexact attribution of geographic coordinates in cases when they were not provided in the original study, and the use of geographic coordinates that were approximate or erroneous in the original publication or database from which they were extracted.

### 4.2 Overestimation of cropland cover at the local scale

To assess the local cropland cover area, we used information available in the original studies, in other publications, on Google Earth around the correct sampling areas, on satellite images from Landsat 8 or Sentinel 2 for more dates, on the internet and in ESA CCI. The information available generally did not allow us to establish precise cropland covers, but often enabled us to determine whether InsectChange estimates of the percentages of land covered by crops were of an adequate order of magnitude, overestimated or underestimated, on the basis of clearly identifiable parts of local land covers. In some cases, we were unable to make a decision, either because the precise sampling location was unknown and could include crops, or because satellite images were difficult to interpret, and we found no other source of information. We found that the assessment of local cropland cover was inadequate for half (486 of 985) of the terrestrial plots, with a very uneven distribution of errors (Figure 7a1, *CroplandCover.xlsx*). Most plots assessed as having no surrounding crops were well assessed (455 of 486 plots), whereas most plots assessed as having surrounding crops suffered from an overestimation of the cropland cover (353 of 499 plots), with 71% of these 353 (252 plots) in fact having no surrounding crops. On the basis of only clearly identifiable parts of the land cover, we found that for 129 geographic coordinates for which assessment was possible, the assessment errors were very wide-ranging: the minimum overestimation of the cropland cover varied between 3% and 100% (mean: 45%, median: 36%, N = 114) and its minimum underestimation varied between 1% and 67% (mean: 15%, median: 12%, N = 15, Figure 7a2). Because of the strong overestimation of cropland cover, we argue that InsectChange cannot provide a reliable analysis of the impact of local cropland cover on insect changes and could lead to incorrect dismissal of the impact of cropland cover on insect decline.

**Figure 7.**
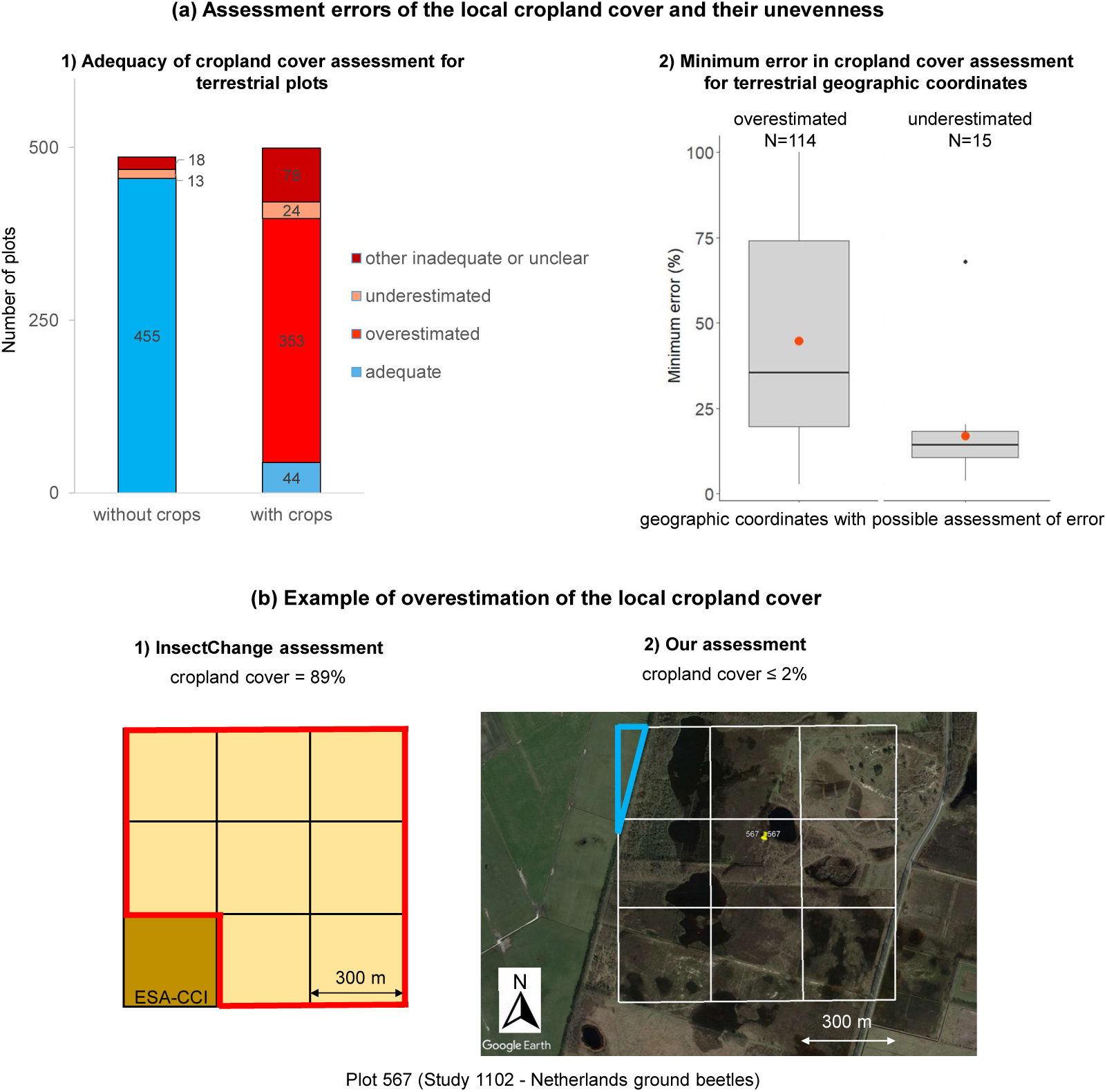
Overestimation of local cropland cover (*CroplandCover.xlsx*). (a) Assessment errors of the cropland cover of plots and their unevenness: 1) InsectChange assessment of local cropland cover of an adequate order of magnitude, overestimated, underestimated or inadequate or unclear (without a possible assessment); 2) Minimum error in cropland cover assessment, based on clearly identifiable parts of the land cover, for geographic coordinates with inadequate cropland cover assessment. (b) Example of overestimation of the local cropland cover in Study 1102 (van Klink et al., 2019), Plot 567, with adequate geographic coordinates (latitude: 52.77986, longitude: 6.57968, last year in study: 2016, cell scale: 300 m × 300 m); 1) InsectChange assessment of cropland cover shown in red = (8 × 100%)/9 = 89% from ESA-CCI 2015 coding, i.e., 8 yellow cells (ESA-CCI code 10, “cropland, rainfed”) coded as cropland in InsectChange and one brown cell (ESA-CCI code 110, “mosaic herbaceous cover > 50%/tree and shrubs < 50%”) coded as uncropped in InsectChange; 2) Our assessment of cropland cover on the basis of information from the source study and Google Earth satellite image from May 2, 2016, showing the local area surrounding the plot in Hullenzand heathland (Netherlands). Most of the area was heath land, whereas the northwestern green area outlined in blue, representing ≈ 2% of the local-scale square, was cropland or grassland. The local cropland cover was therefore either 0% or ≈ 2%. On this basis, the 89% assessment in the database was coded as overestimated in our analysis (Appendix S1, Problems.xlsx).

Inadequate geographic coordinates explained only 18.3% of inadequate cropland cover assessments. Indeed, for more than half (127 out of 233) of the plots that were assigned inadequate geographic coordinates, the actual sampling area and the local surroundings totally lacked crops, in line with the InsectChange assessment for these plots. In most cases, therefore, the inadequacy of geographical coordinates had no impact on the assessment of local cropland cover. The main reason for inadequate cropland cover assessments was the inaccurate interpretation of satellite images by the ESA-CCI database (*CroplandCover.xlsx*), notably because grasslands, heathlands, steppes, barrens, prairies, shrublands, marshlands, natural vegetation areas, parks or golf courses may inaccurately be coded as croplands (Peng et al., 2017; Liu et al., 2018), and the representation of land cover is imprecise when used at a local scale composed of nine 300 m × 300 m squares with rough cropland cover assigned to each of them (63.2% of inadequate cropland cover assessments, *CroplandCover.xlsx*, example in Figure 7b). For some plots (but not systematically for all plots), we checked whether cropland covers were adequately retrieved from ESA-CCI. We found that this was not the case for 8.2% of cropland covers, were the InsectChange assessment did not match with ESA-CCI information. Finally, 10.3% of the cropland cover assessments were inadequate for other reasons (for example, insufficient resolution of satellite images at the beginning of the 1990s, tree cover, parking lots, and shadow on the top of a mountain incorrectly coded as cropland).

In terms of freshwater, 49 of the 62 studies matched with ESA-CCI information included inadequate geographic coordinates that should be used with caution. We did not check local cropland cover estimates, as the water quality at the sampling points may be more dependent on upstream land use than on land use of immediately adjoining plots (Allan, 2004; Desquilbet et al., 2020). For terrestrial and freshwater datasets, a quality check of the accuracy of estimates provided for other possible drivers of insect change, notably local-scale drivers (urban cover and climate change) similarly affected by the inadequacy of geographic coordinates, is strongly recommended.

## Conclusion

The numerous problems affecting the InsectChange database call for corrections and extreme vigilance in its use. They call into question the results obtained thus far from this database, in the first place those of van Klink et al. (2020a), which were widely covered by various media reaching a broad readership (Kimbrough, 2020; McGrath, 2020; Ritchie, 2024). The main consequence is that InsectChange conveys unsubstantiated information to scientists, decision-makers and the general public. We argue that InsectChange, in its current state, does not allow the study of insect trends worldwide or their drivers and is particularly unsuitable for the analysis of the influence of agriculture on insects, or for the study of changes in freshwater insect assemblages. We have outlined ways of improving data selection to make the data more representative of habitat conditions and insect numbers at a global scale. Our detailed appendices are designed to facilitate data consolidation. More generally, this comment underlines the need for relevant matching with external databases. Our careful review also illustrates the value of contacting dataset owners to ensure their appropriate use and calls for vigilance to avoid transferring errors across databases, as occurred for 11 datasets incorporated from the Global Population Dynamics Database (Prendergast et al., 2010) and/or Biotime (Dornelas et al., 2018) into InsectChange (Appendix S1). Finally, our in-depth analysis highlights the attention that should be given to the data and their meaning to ensure that large databases built from individual datasets participate in a cumulative knowledge process.

## Supporting information

Problems

FreshwaterNonInsects

CroplandCover

## Appendices

**Appendix 1.**
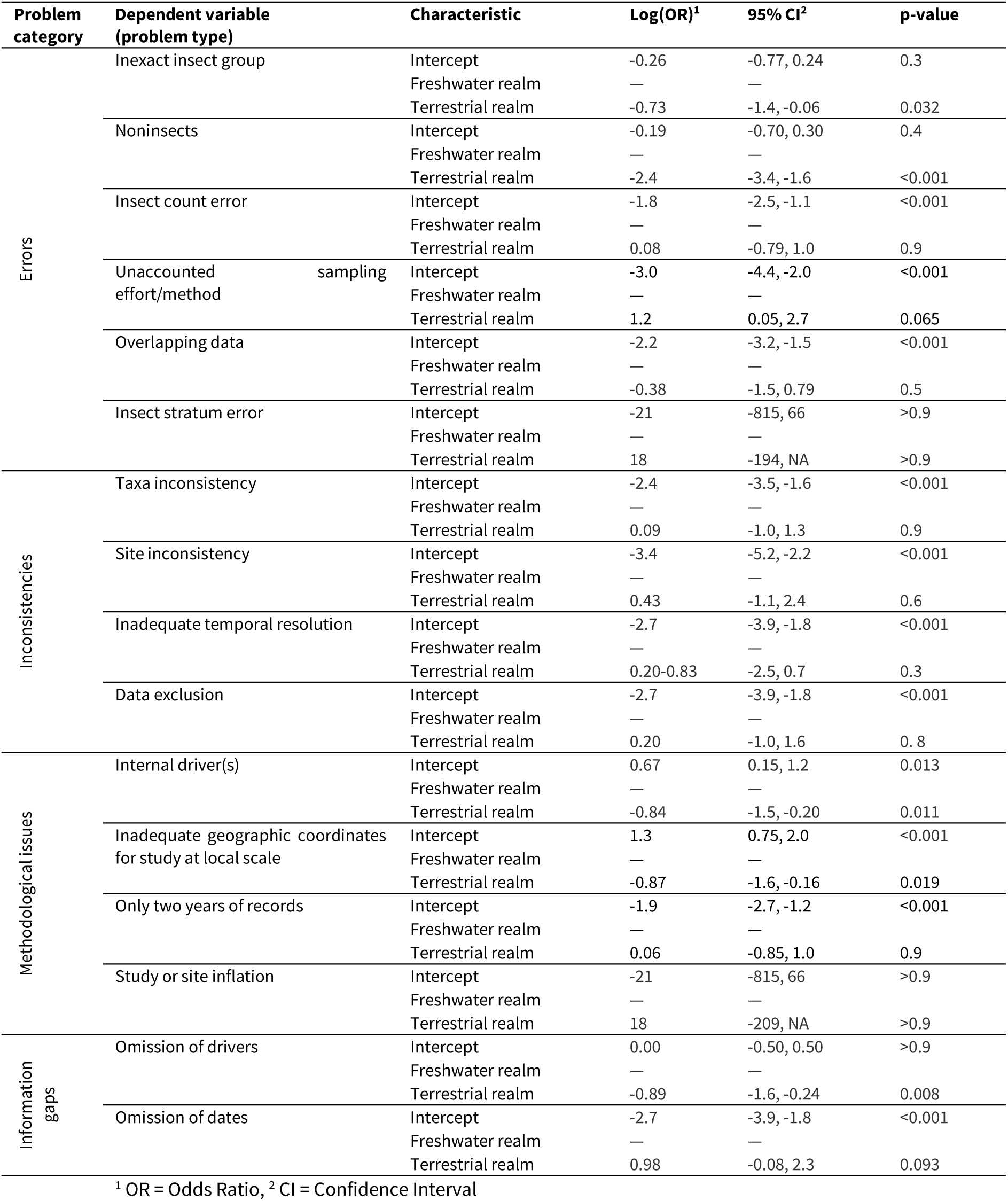
Results of the binary logistic regressions testing for an effect of realm on each problem type.

**Appendix 2.** Effects of considering total freshwater invertebrates instead of freshwater insects only on trend estimation. (a) Results of mixed model testing for effects on insect density (log10-transformed) of fixed variables with respect to time, type of assemblage considered and metric and random variables with respect to datasets and plots within datasets. (b) Trend estimation (in the log10 space) between the different groups and tests for their differences.

**(a) Mixed linear model on log<sub>10</sub>(insect density) dependent variable**
| **- Type III ANOVA-like table with Satterthwaite's method for the fixed effects** |  |  |  |  |  |  |
| --- | --- | --- | --- | --- | --- | --- |
| Independent variable | Sum of squares | Mean square | Num.df | Den.df | F value | P value |
| Assemblage type | 0.799 | 0.799 | 1 | 124.85 | 6.351 | 0.013 * |
| Metric | 0.356 | 0.356 | 1 | 125.80 | 2.861 | 0.093 |
| Time | 0.137 | 0.137 | 1 | 125.53 | 1.104 | 0.295 |
| Assemblage type * Metric | 1.271 | 1.271 | 1 | 124.85 | 10.220 | 0.002 ** |
| Time * Assemblage type | 0.832 | 0.832 | 1 | 124.84 | 6.694 | 0.011 * |
| Time * Metric | 0.240 | 0.240 | 1 | 125.53 | 1.938 | 0.166 |
| Assemblage type*Time*Metric | 1.290 | 1.290 | 1 | 124.84 | 10.371 | 0.002 ** |
| **- ANOVA-like table for the random effects** |  |  |  |  |  |  |
| npar |  | logLik | AIC | LRT | Df | P value |
| none | 11 | -77.573 | 177.15 |  |  |  |
| 1 \| Dataset_ID | 10 | -78.422 | 176.84 | 1.697 | 1 | 0.193 |
| 1 \| Dataset_ID : Plot_ID | 10 | -90.546 | 201.09 | 25.946 | 1 | 3.5e-07*** |

**(b) Time trend estimates for each group**
| Assemblage type | Metric | Time.trend | SE | df | t.ratio | P value |
| --- | --- | --- | --- | --- | --- | --- |
| Total invertebrates | Abundance | -0.00705 **a** | 0.00353 | 126 | -1.995 | 0.048 * |
| Insects | Abundance | -0.00417 **a** | 0.00353 | 126 | -1.181 | 0.240 |
| Total invertebrates | Biomass | 0.01399 **b** | 0.00371 | 124 | 3.768 | 0.0003 *** |
| Insects | Biomass | -0.01242 **a** | 0.00669 | 125 | -1.855 | 0.066 . |
| Total invertebrates | Abundance + Biomass | 0.00347 **a'** | 0.00256 | 125 | 1.353 | 0.1786 |
| Insects | Abundance + Biomass | -0.00830 **b'** | 0.00378 | 125 | -2.192 | 0.0302 * |

\*: P <0.05, \*\*: P <0.01, and \*\*\*: P <0.001
Different letters were associated with the trend estimates when they were significantly different from each other according to post hoc pairwise tests with Satterthwaite's method and Holm's correction for multiple comparisons. The model run without the non-significant factor Dataset\_ID gave comparable results and was not better according to the Akaike information criterion (AIC) with a AIC difference of only 0.3.

## Acknowledgments

We thank Sonja Jähnig for helpful discussions, Gaëlle Viennois for verifying the land cover analysis, Dirk Maes and Hans van Dyck for their help in analysing biased data on Belgian lepidopterans (Study 70), other coauthors of Desquilbet et al. (2020), Sylvain Chabé-Ferret for insights into the analysis of controlled and natural experiments, and Dominique Lamonica and Gilles LeMoguédec for checking the statistics and the argumentation related to trend estimation from heterogeneous data. We also thank Marianne Elias, Marilyne Laurans, Doyle McKey, Denis Bourguet, as well as Bradley Cardinale and another reviewer for their helpful comments on the manuscript.

## Conflict of interest disclosure

We declare that we have complied with the PCI rule of having no financial conflicts of interest in relation to the content of the article.

## Funding

This work was publicly funded by the French National Research Agency (ANR-17-EURE-0010 to MD, ANR-16-IDEX-0006 to LG, *Investissements d’avenir* program).

## Author contributions

Marion Desquilbet and Laurence Gaume contributed equally to this work.

## Data, scripts and codes availability

Data, scripts, outputs and images for the analyses of problem types and invertebrate trends are available from the Figshare repository as a RStudio project in the compressed file *Stat_Invertebrates.zip* (https://doi.org/10.6084/m9.figshare.23458877).

## Supplementary information availability

Supplementary information is permanently archived in the Figshare repository (https://doi.org/10.6084/m9.figshare.23458877). In addition to *Stat_Invertebrates.zip*, it comprises three appendices (Appendix S1, Appendix S2 and Appendix S3) and four datasheets (*Problems.xlsx, FreshwaterNonInsects.xlsx, CroplandCover.xlsx* and *Fig2and5.xlsx*).

- Appendix S1 and *Problems.xlsx* detail and record the problems encountered in InsectChange.
- Appendix S2 summarises the inclusions of freshwater noninsects in the InsectChange assemblages, whereas *FreshwaterNonInsects*.xlsx details the calculation of their parts in the assemblages and the problematic consequences for the inferred ‘insect’ trends.
- Appendix S3 focuses on studies analysing the effect of internal drivers and highlights the driver-induced situations that favour an increase or decrease in insects.

- *Fig2and5.xlsx* provides information supporting Figures 2 and 5 of this comment.

## Notes

### Competing Interest Statement

The authors have declared no competing interest.

### Summary of Updates

The article has been recommended by PCI Ecology: https://doi.org/10.24072/pci.ecology.100641. The PCI Ecology badge has been added on the 1st page.

