## Appendix S1 for "InsectChange: Comment"

**Supporting Information** for "*InsectChange: Comment*". Laurence Gaume, Marion Desquilbet.

### Appendix S1.

### Detail of the problems encountered in the datasets of InsectChange

##### Presentation of the different appendices

This appendix describes the different problem types encountered in studies of the InsectChange database (van Klink et al. 2021). References to original data sources are available in van Klink et al. (2021) and *Problems.xlsx*. *Problems.xlsx* provides a table summarising the different problem types for each study and also provides the specific factor in each study with an experimental factor and/or a major disturbance as well as the associated habitat change and expected change in insect dynamics. *Fig2and4.xlsx* presents information supporting Figures 2 and 4 illustrating the main problems highlighted in the comment as well as the biases they induce. Appendix S2 and *FreshwaterNonInsects.xlsx* provide detailed information on non-insects composing the freshwater assemblages as well as - when possible - the calculation of their share in the assemblage considered and the comparison of their trends with those of the assemblages considered. Appendix S3 details the expected trends in the studies affected by internal drivers linked to experimental conditions or major disturbances. Details on our assessment of the adequacy for a study at a local scale of the geographic coordinates provided by InsectChange and/or the adequacy of the local cropland cover provided for terrestrial plots are presented in *CroplandCover.xlsx*. When not specified, satellite images refer to a visualisation on Google Earth.

#### Errors and problems transferred from other databases to InsectChange

Errors and problems were transferred from other databases to InsectChange (van Klink et al. 2021) in 11 studies, which are marked with an asterisk: studies 63, 70 and 380 incorporate errors from the Global Population Dynamics Database (GPDD) (Prendergast et al. 2010) that were transferred to BioTIME (Dornelas et al. 2018), then to InsectChange; studies 79, 465 and 502 incorporate errors from the GPDD; studies 249, 294, 301, 313 and 375 incorporate errors from BioTIME, as detailed below. These errors and problems impacted multiple publications: for example, the datasets of Blowes et al. (2019), Antão et al. (2020) and Daskalova et al. (2020) include BioTIME studies 63, 70, 249, 294, 301, 313, 375 and 380 and the dataset of Dornelas et al. (2019) includes BioTIME studies 63, 70, 249, 301, 313 and 375.

#### Other problems not included in our analysis

We draw attention to examples of other information gaps or errors in InsectChange that are not included in our analysis due to their lesser importance. First, in the table *SampleData.csv* of InsectChange providing information on sampling methods and data extraction, information on biomass is missing for 23 studies (375, 1382, 1411, 1415, 1416, 1425, 1427, 1428, 1448, 1449, 1453, 1455, 1457, 1473, 1488, 1493, 1496, 1506, 1507, 1508, 1509, 1510, 1524). Second, the table *PlotData.csv* of InsectChange providing details at the plot level indicates as the source of geographical coordinates "NOT PRECISE!" for the 8 plots of study 1261. Third, the table *DataSources.csv* providing descriptive data at the level of the study erroneously indicates in column J referring to number of plots that study 1473 contains 11 plots: for this same study, the table *PlotData.csv* provides information on 19 plots while the table *InsectAbundanceBiomass.csv* containing the insect abundance or biomass numbers inconsistently provides information on 20 plots. This invites users to pay attention to possible problems that may not be included in our analysis.

**New errors in the erratum by van Klink et al. (2020b)**

The InsectChange database presents the data underlying the analysis by van Klink et al. (2020a). The analysis below shows that InsectChange still contains an important number of problems in the data after the erratum by van Klink et al. (2020b). It also shows that this erratum actually introduced some new problems that were absent from the pre-erratum analysis by van Klink et al. (2020a), as was the case for the addition of Plots 455 and 459 to the database (study 1452, written in Russian). These plots deal with entire freshwater invertebrate assemblages shown in detailed tables of source study to increase exponentially over the study period because of the reported proliferation of invasive molluscs.

**Details of the problems encountered in each study**

**Study 63* (Dornelas et al. 2018) – BioTIME UK dragonflies**

The original source of this dataset is Moore (1991). It was successively included in the Global Population Dynamics Database (GPDD) (Prendergast et al. 2010), then BioTIME (Dornelas et al. 2018), then in InsectChange. This study examines the colonisation of 20 newly built ponds by freshwater dragonfly communities. This disturbed context is not specified in InsectChange. The measure of abundance is the total number of mature males on the days of their maximal abundance for the year indicated, while it is not clear how this total number of mature males compares to other insect numbers in InsectChange. Also, an error from the GPPD in the number of insects reported from Table IV of the source publication was passed on to InsectChange. The data series begins in 1962, not 1959. The total number of insects is 32 (and not 126) in 1962 and 46 (not 255) in 1963.

The reason is that the number of *Libellula quadrimaculata* is reported in the GPPD as 130 in 1959 and 45 in 1961 (while no data exist for these years), 95 instead of 1 in 1962, and 210 instead of 1 in 1963. The geographic coordinates assigned in InsectChange from Google maps are inappropriate as they are located 600 m northwest of the ponds, which are visible on Google Earth.

**Study 70* (Dornelas et al. 2018) – BioTIME Belgian Migrating Lepidoptera**

The original source of this dataset is a series of articles (Vermandel, annual from 1985 to 1995; Vanholder, 1996 and 1997). It was included in the GPDD (Prendergast et al. 2010) with a reference to "Vanholder, B. (1997) Belgian Migrating Lepidoptera. Hard Copy – Tabular" and a now broken internet link. It was then successively included from the GPDD to BioTIME (Dornelas et al. 2018), then from BioTIME to InsectChange. This time series deals with the abundance of migratory Lepidoptera, including butterflies and moths. They are inadequately referenced as "Butterflies" in the GroupInData cell of the table *SampleData.csv* of Insect Change. The time series shows a 300-fold increase in the abundance of these migratory lepidopterans in Belgium from 1983 to 1996. However, the sampling effort and the number of sampling sites increased over time. In particular, the number of observers was multiplied by 9, from 18 to 156, between 1983 and 1996. Therefore, this dataset leads to overstate the increasing insect trend or to consider an increasing instead of a decreasing trend compared with the original study. The error comes from the GPPD, which inadequately indicates in the notes of its table *DataSource.csv*: "Survey comprises 180 observers that are regular recorders of Lepidoptera". It should also be noted that the series extends beyond 1996, up to 2018. Besides, this series only includes migratory species. Their numbers can strongly fluctuate and depend on successful breeding in the Mediterranean region and migration to Northern Europe in spring (Stefanescu et al. 2007; Stefanescu et al. 2011). Therefore, it is not appropriate to study their abundance in relation to local conditions in Belgium only. Finally, the location of the unique plot considered in this study is not relevant for either a local or landscape-scale study, as butterflies were sampled in various locations in Belgium. For this reason, the estimation of the local cropland cover in the data paper is incorrect.

**Study 79* (NERC Centre for Population Biology Imperial College 2010) – GPDD UK butterflies**

These observational data on butterfly abundance in the UK between 1976 and 1985 were extracted from the GPDD (Prendergast et al., 2010), where they were included after extraction from Pollard et al. (1986). First, there were errors in the inclusion or exclusion of sites in InsectChange. In Appendix S2 of InsectChange, van Klink and coauthors specify: "*[s]ome species were not consistently reported in the source data, so we removed all years with missing species. We also excluded years in which 0 butterflies were recorded in the entire year. After this exclusion process, we excluded 15 plots of too short duration*". However, three plots were inconsistently excluded while they should have been included as they covered a period of nine years after removing all years with missing species or no butterfly records: the Gibraltar Point National Nature Reserve, the Lindisfarne National Nature Reserve and the Potton Wood (respectively Plots 388, 395 and 401 in the GPDD). On the contrary, two plots were inconsistently included as the data only covered a period of eight years after removing years incorrectly included despite missing species: Foxholes (Plot 866 in the data paper, 387 in the GPDD), for which year 1976 should have been removed; the Rostherne Mere National Nature Reserve (Plot 878 in the data paper, 403 in the GPDD), for which year 1984 should have been removed. Also, other plots were correctly included, but some years were included despite missing species and thus the dataset did not account for inter-year variations in sampling effort for these plots: year 1976 for the Brockwells Farm (Plot 860 in the data paper, 380 in the GPDD), year 1976 for the Holme Fen National Nature Reserve (Plot 868 in the data paper, 391 in the GPDD), year 1985 for the Leighton Moss RSPB (Plot 871 in the data paper, 394 in the GPDD), year 1985 for the Oxwich National Nature Reserve (Plot 875 in the data paper, 400 in the GPDD), year 1985 for the Leighton Moss RSPB (Plot 871 in the data paper, 394 in the GPDD). In addition, some data from Pollard et al. (1986) were incorporated with errors in the GPDD and these errors were transmitted to InsectChange. For example, for Bevills Wood (Plot 859 in the data paper, 379 in the GPDD), the numbers were incorrectly reported for some butterfly species in some years (peacock, small copper and small white) and two species (purple hairstreak and holly blue) were not reported in the GPDD. In particular, for this plot, the number of small white butterflies was incorrectly reported as 20 instead of 209 in 1978 in the GPDD, leading to an incorrect total number reported in InsectChange of 805 instead of 994 in this year. Analogous errors are encountered in other plots. Finally, data of the UK butterfly monitoring scheme are included in two different studies (study 79, spanning from 1976 to 1985 and study 380, spanning from 1978 to 1987), without overlapping data: study 380 includes one plot, the Castle Hill National Nature Reserve, which is included in Pollard et al. (1986) but has too short duration for inclusion in study 79 due to missing data. In both studies, the primary source of data is identified as the UK butterfly monitoring by Ernest Pollard and colleagues. It would have been consistent with the rest of the dataset to consider both studies as different sites of a same study.

**Study 249* (Thomsen et al. 2016) – BioTIME DK light-trapped insects**

This study was extracted from the BioTIME Database (Dornelas et al. 2018), where it was included after extraction from Thomsen et al. (2016). It presents the dataset of an interrupted time series natural experiment examining insect responses to local temperature using a sampling site with more than a quarter-million records from two decades (1992–2009) of full-season, quantitative light trapping of 1543 species of moths and beetles. As the temperature data were not included in the dataset that included insect abundance data, we did not classify it as a natural experiment. Appendix S2 of InsectChange states: "We retained only the months June, July, August, because the other months were not sampled consistently. Years in which no summer samples were taken, including the first year of sampling (1992), were removed. The summer months were summed to provide one value per year". However, insect counts retained in the data paper do not correspond to this description. In years 1993, 1995, 1996 and 2006, where sampling periods add up exactly from June 1st to August 31, insect counts are respectively 5,161, 17,196, 10,133 and 28,963 in the original dataset, while they are systematically higher, in variable proportions, in InsectChange, with respective values of 5,860, 17,309, 10,146 and 29,124. The sampling periods did not add up exactly to June 1st-August 31 in other years. In these years, even taking longer sampling periods to include the whole months of June, July and August, insect counts are systematically higher in InsectChange, in variable proportions, except in 2002 and 2003, when they are lower. In BioTIME, the column "comments" in the table *BioTIMEMetadata_24_06_2021.csv* inaccurately indicates: "*Counts are aggregated over a 10 day period throughout the year. Dates in dataframe are median date between when light trap is set up and light trap is examined*", and therefore this dataset is prone to be used with unaccounted change in sampling effort. Actually, sampling periods in Thomsen et al. (2016) range from 1 to 39 days. For example, for insects trapped during the 39-day period from 11/1/2004 to 12/9/2004, BioTIME inadequately reports month 12 and day 5, and indicates in the SAMPLE_DESCR column "5_11.5_2004", from which the user cannot infer the actual length of the sampling period. Finally, in the table *DataSource.csv*, moths and beetles are referenced as "Coleoptera and Lepidoptera". As the Lepidoptera taxon also involves butterflies, it is more appropriate to reference the invertebrate group as "Coleoptera and Lepidoptera (moths)".

**Study 294* (Dornelas et al. 2018) - BioTIME Tam Dao butterflies**

The original source of this dataset is Bonebrake et al. (2016). It was included in the BioTIME database (Dornelas et al., 2018), then from BioTIME to InsectChange. This natural experiment investigates butterfly community changes in six different habitat types (shrubs and agricultural area, disturbed forest, closed forest, old road, secondary forest, bamboo forest) from 2003 to 2013, thereby examining the impacts of habitat degradation following intensive infrastructure development in 2005. These natural experimental conditions are not reported in BioTIME or InsectChange. The six habitat types are considered in BioTIME and InsectChange as six plots. In both databases, they inadequately share a unique pair of geographic coordinates, while from Vu et al. (2009) they were up to three kilometres apart. For this reason, the estimation of the local cropland cover in InsectChange is incorrect. In addition, plot 237 includes seven abundance counts in one month, and five or less in all other months. This is not consistent with the methodology detailed in InsectChange (p. 17 of the detailed presentation), describing that the temporal resolution was between the week and the year, except in six datasets where plots were sampled between six and eight times in any month. Finally, there are several samplings in most months but InsectChange does not allow to know their chronology because it does not report the sampling date.

**Study 300 (Landis 2018) - LTER Kellog insects**

This study is a long-term crop experiment with 51 plots harbouring different crops and agronomic managements (42 crop plots studied from 1989 to 2013 and nine succession and forest plots studied from 1993 to 2013). It examines the effect of different controlled experimental treatments carried out in each plot on plant dwelling insects pertaining to the "herbivore predators" category. These controlled treatments introduced heterogeneity, but they are not reported in InsectChange. The *DataSource.csv* table refers to "insects" while it would have been more adequate to specify "plant dwelling insects (herbivore predators)" since the source dataset focuses on this category of insects and includes 14 species of Coccinellidae, one species of Chrysopidae, and one species of Lampyridae. InsectChange includes only the Coccinellidae, as adequately specified in the table *SampleData.csv*. The local cropland cover provided in InsectChange is not consistent with ESA-CCI information and, depending on plots, is an overestimation or an underestimation compared with ESA-CCI information (ESA 2017).

**Study 301* (Joern 2016) – BioTIME Konza grasshoppers**

This experimental study is carried out to evaluate the effects of fire and grazing in Konza Prairie on different ecological variables including insect abundance. This dataset was extracted from BioTIME (Dornelas et al. 2018). The 15 plots included in InsectChange received an array of experimental burning and grazing (especially bison) treatments, which are not reported in InsectChange and were mentioned but not detailed per plot in BioTIME. One unique pair of geographical coordinates, corresponding to the location of the Konza Prairie Biological Station, was inadequately assigned to the 15 plots in BioTIME and InsectChange. Actually, each plot corresponded to the starting point for a sweep sample of grasshoppers across Konza Prairie, and the plots were up to approximately four kilometres apart, and at least one kilometre away from the Konza grassland biological station. For this reason, the estimation of the local cropland cover in InsectChange is incorrect. These problems pertaining to the BioTIME database were corrected in the latest BioTIME revision of June 24, 2021 (10.5281/zenodo.5026943), by which the BioTIME dataset 301 was changed. It now includes corrected data for watershed 002d of the Konza Prairie grasshopper dataset and 13 additional datasets, numbered from 528 to 540 for the other watersheds. In addition, three plots (195, 196 and 198) include six or seven abundance numbers, each time only in one month, and five or less in all other months. For example, for Plot 195, there were seven separate records in July 1987, with no further indication, which actually correspond to daily records on July 1, 6, 13, 20, 28, 29 and 31. This is not consistent with the methodology detailed in InsectChange (p. 17 of the detailed presentation), describing that the temporal resolution was between the week and the year, except in six datasets where plots were sampled between six and eight times in any month. Moreover, there are several samplings in most months but InsectChange does not allow to know their chronology because it does not report the sampling date, which was however available in the original dataset.

**Study 313* (Knops and Tilman 2006) - BioTIME LTER Cedar Creek grasshoppers**

This natural experiment examines grasshopper dynamics in different fields of the Cedar Creek LTER previously cultivated, but abandoned from agriculture at various times in the past. This dataset was extracted from BioTIME (Dornelas et al. 2018). The BioTIME file *BioTIMEMetadata_24_06_2021.csv* only mentions that the agricultural fields were "abandoned at various times", while the original dataset specifies for each field the time after agricultural abandonment, varying from 6 to 50 years. The DetailedPlot cell of the table *SampleData.csv* of InsectChange inadequately mentions for each of the plots *"started some 20 years after abandonment. Prescribed burning"*, while first, the time after agriculture abandonment varied from 6 to 50 years depending on the plots included in InsectChange, and second, the prescribed fires reported here began in 2006, after the end of the observation period included in InsectChange, and in all plots except one. The same geographic coordinates were inadequately assigned to 20 plots corresponding to 20 old fields of the Cedar Creek LTER, up to approximately 4.6 km away from each other, in BioTIME and in InsectChange. It was specified in the original dataset that *"[o]nly the family Acrididae was recorded throughout the entire survey period (1989-2006). Other families were sorted and identified in all years except 1992 and 1993."* This smaller sampling effort in 1992 and 1993 was not accounted for in InsectChange, where abundances of recorded grasshoppers were summed by month and year regardless of the change in considered grasshopper families. This results in an artificial dip in the abundance curve in 1992-1993, early in the observation period. This change in sampling effort could have been addressed by removing the 1992 and 1993 observation years or by focusing only on the family Acrididae, whose sampling remained constant over the years.

**Study 375* (Dornelas et al. 2018) - BioTIME Japan ground beetles**

This observational study examines the dynamics of ground beetles in different forest sites in Japan. The dataset, covering the period 2004-2014, was extracted from BioTIME (Dornelas et al. 2018). The primary source of data indicated in BioTIME is the website of the Japanese ministry of environment (Monitoring Site 1000 Project, Biodiversity Center, Ministry of Environment of Japan (2014), where the data is available in Japanese. The data paper in English by Niwa et al. (2016) allows an easier access to the primary data for the shorter period 2004-2012. By comparing the dataset of Niwa et al. (2016) with the dataset included in BioTIME, we identified several shortcomings in BioTIME, which were carried over into InsectChange. In this study, in each plot, ground beetles were sampled at various dates usually, but not always, in 5 subplots, each containing four pitfall traps, after opening the traps for three days. For each subplot of each plot, the dataset by Niwa et al. (2016) distinguishes situations where trapping was not carried out from situations where no beetle was captured and informs on situations where trapping was carried out for more or less than three days and/or with less than four traps. This information on the sampling effort is absent in BioTIME. BioTIME only includes subplots with a positive record of insects at a given date, without information on the duration of trapping and the number of traps. Zero records, corresponding to situations where traps were in operation but no beetle was captured in any subplots of a given plot, were not included in BioTIME, resulting in errors in insect counts. When beetles were captured in some but not all subplots, it is possible to identify missing subplots in BioTIME, but it is not possible to know whether no trapping was carried out or whether no beetle was captured in these subplots. Niwa et al. (2016) also provide information on disturbances in each plot, including strong typhons in 2004 in Plots TM-DB1 and TM-DB2 (257 and 258 in InsectChange) and typhon disturbance in 2004 and 2005 in Plot AY-EB1 (244 in InsectChange), which are not reported in BioTIME or InsectChange. InsectChange includes the sum of all subplots per plot at sampling dates available in BioTIME (that is, excluding dates where sampling was carried out and no beetle was captured). It carries over the unaccounted changes in sampling effort in BioTIME, contrary to the description in Appendix S2 of InsectChange that all plots that were not sampled consistently were removed. It also amplifies the problems in BioTIME in two ways. First, it does not inform on the number of subplots with positive beetle records at each date. Second, in months with several samplings, it does not report the sampling date, which was however available in BioTIME, and therefore does not allow to know their chronology. In addition, contrary to its methodology, it includes daily records, some of them occurring in a same week, without maintaining the week as the finest temporal grain. We note further inadequacies in the description of this study in Appendix S2 of InsectChange: ten plots, not seven, were excluded for too short duration, leaving 22 plots, and not 26, in InsectChange. Finally, the estimation of the local cropland cover provided in InsectChange is not consistent with satellite images for Plots 243 and 259 and unclear for Plot 256.

**Study 380* (Dornelas et al. 2018) - BioTIME UK chalk grassland butterflies**

The original source for this dataset is the book chapter by Pollard, 1991. It was successively included in the GPDD (Prendergast et al. 2010), then in BioTIME (Dornelas et al. 2018), then in

InsectChange. The geographical coordinates indicated in these three databases are inadequate: according to the map on p. 99 of this book, the route used for monitoring butterflies in Castle Hill National Nature Reserve is between Newmarket Hill and Castle Hill, Sussex, which according to Google Earth is 8 km away from the location used in these three databases. Moreover, there is an unaccounted change in sampling effort: a missing value for the first flight period of peacock (*Inachis io*) in 1978 is inadequately considered as a zero count in BioTIME, where the abundance numbers for this butterfly species correspond to the first flight period in 1978 and the sum of the first and second flight periods in subsequent years, and in InsectChange, where abundance numbers of all butterfly species are summed up without taking into account this missing value. There are also errors in insect counts: the GPDD did not report from the original source the counts of several butterflies: the small/Essex skipper (*Thymelicus sylvestris*), the clouded yellow (*Colias crocea*), the brimstone (*Gonepteryx rhamni*), and the first flight period of the green-veined white (*Pieris napi*), which are therefore not considered in BioTIME and InsectChange. As detailed in the comments on study 79, it would have been adequate to consider studies 79 and 380 as two sites of a same study for consistency with the rest of the database.

**Study 465* (NERC Centre for Population Biology Imperial College 2010) - GPDD Prague moths**

These observational data on moth abundances in the Czeck Republic between 1967 and 1992 were extracted from the GPDD (Prendergast et al. 2010). From Kadlec et al. (2009), the geographic coordinates indicated in the GPDD and reported in InsectChange are not accurate and are located 30 km from the light trap where moths were sampled (Kadlec et al. 2009). This leads to an overestimation of the local cropland cover in InsectChange.

**Study 478 (Wagner et al. 2011) - Germany EPT Breitenbach**

This observational study from the long-term monitoring program of the Breitenbach stream (Germany) reports the dynamics of freshwater insects at different sampling sites along the stream. The dataset includes data for Ephemeroptera, Plecoptera and Trichoptera only at 5 plots spaced along the stream for different periods within 1969-2004. The distances to the mouth and elevations of the plots are not reported in InsectChange. The five plots included in InsectChange were inadequately assigned the same geographic coordinates, while they were up to approximately 2.6 km apart according to the source study by Wagner et al. (2011). In 1986, during the dataset period, the accidental introduction of an insecticide into the stream system reset almost all arthropod populations in the study stretch to zero (Wagner et al. 2011, p. 201). This highly disturbing event is not reported in InsectChange. Furthermore, while the source dataset systematically provides the abundance data on a daily basis, InsectChange summed up the abundance data to get the total sampling by year, instead of producing a value per week. First, this is in contradiction with the methodology of InsectChange stating that "in data sets where sampling took place at a higher frequency [than month or year] (i.e. daily), these sampling events were summed or averaged, to produce one value per week or month (averaging was only done in cases of variable sampling effort, e.g. due to randomly missing samples)". Second, this sum of abundance counts per year is inadequate because it does not account for two types of changes in sampling efforts: (i) each leap year has one sampling day more than the others; (ii) in plot 575 (Site T4 of the source dataset), sampling began on March 1977 and ended in June 1993, with only 289 sampling days in 1977 and 181 sampling days in 1993, against 365 or 366 sampling days in other years depending on leap years.

**Study 502* (NERC Centre for Population Biology Imperial College 2010) - GPDD UK aphids**

The original source of this dataset on aphids is the Rothamsted Insect Survey Annual Reports covering the period 1969-1990. It was successively included in the GPDD (Prendergast et al. 2010), who quoted Taylor et al. (1990) as the source study, then from the GPDD to InsectChange. By examination of the site names in the GPDD, and of the locations of corresponding suction traps of the Rothamsted Insect Survey, we find that the geographic coordinates of the sites in the GPDD, reported in the table *PlotData.csv* of InsectChange, are inadequate, as can be seen from comparison with those in Morales-Hojas et al. (2020). Notably, sites 895, 896, 900 and 902 (respectively, Hereford, Rothamsted Experimental Station, Wye College and Starcross), are actually the same as sites 1617, 1618, 1620 and 1619, respectively, in study 1495 on flying insects (including aphids) in the period 1973-2001. Therefore, some insect data from this study overlaps with that of study 1495. It would have been appropriate to select only one of these datasets. Furthermore, 4-week periods in the GPDD (the period covering weeks 17-20 to the period covering weeks 41-44, per steps of four weeks) were inadequately coded as months 5 to 11 in InsectChange, while for example in the first year of the time series, 1971, weeks 17-20 covered the period from April 24 to May 21, and thus not only May (5); weeks 41-44 covered the period from October 9 to November 5, and thus not only November (11).

**Study 1006 (Rennie et al. 2018a) - ECN moths**

This observational study examines the dynamics of butterfly populations in different sites in the UK. The precise sampling areas in the different sites of the UK Environmental Change Network (ECN) data are not known from the original study. In addition, Plot 1670 representing the Cairngorms site has been assigned erroneous geographic coordinates, approximately 450 km apart from the correct location of the site. The estimation of the local cropland cover provided in

InsectChange is overestimated for some plots compared to satellite images and not possible to assess for others owing to a variability of the cropland cover within the ECN sites.

**Study 1102 (Bowler et al. 2017, van Klink et al. 2019) - Netherlands ground beetles**

This natural experiment examines the drivers of community stability and synchrony, and their relationship with disturbance, species richness, and functional diversity using a dataset of ground beetles trapped from 1959 to 2016 in 19 plots of two European lowland heathlands and grasslands. In 10 of these plots, severe disturbances (fire, pest outbreaks, or restoration measures) occurred at the start of or during the monitoring program, but these severe disturbances, which could impact beetle abundance, were not mentioned in InsectChange. In addition, the estimation for the local cropland cover provided in InsectChange is inconsistent with information from the source study and satellite images, with an overestimation or an underestimation depending on plots.

**Study 1261 (Ellison 2017) - LTER Harvard forest ants**

This controlled experiment from the Harvard Forest LTER examines the ant dynamics in forests submitted to different experimental treatments for hemlock, a foundation tree species, from 2003 to 2015. For this study, InsectChange does not consider ant individuals, but the number of ant species per trap as a proxy for ant nests. The "GroupInData" cell of the table *SampleData.csv* inadequately refers to "Ants", as is done in other studies of InsectChange dealing with ant individuals, while it would be more appropriate to refer to "Ants (proxy for ant nests)". Two experimental treatments were performed by girdling all hemlocks to simulate death by adelgid and logging all hemlocks >20 cm diameter and other merchantable trees to simulate pre-emptive salvage operations. These treatments were paired with two controls: hemlock controls that were beginning to be infested in 2010 by the adelgid and hardwood controls that represented future conditions of most hemlock stands in eastern North America (see e.g. Ellison et al. 2010). Additionally, by summing the number of nests over all traps per plot, InsectChange failed to account for a major change in sampling effort, as pitfall-trap sampling effort doubled from 2012-2015 relative to 2003-2011 (see Welti et al. 2021). Moreover, there are two samplings for Plot 648 in June 2005 but InsectChange does not allow to know their chronology because it does not report the sampling date, which was available in the original dataset. Finally, the same geographic coordinates were inadequately assigned to the eight plots of the study, while they were up to approximately 1.5 km away from each other.

**Study 1263 (Rennie et al. 2018b) – ECN butterflies**

This observational study examines the dynamics of butterfly populations in different sites in the UK. InsectChange only includes data up to 2012 for plots 904 to 909 and 912 to 915 while data exists up to 2015 in the corresponding sites (T021, T022, T03, T041, T042, T05, T08, T09, T10 and T11) of the original dataset. Also, the data includes plots T021 and T022 both located on the same site (Glensgaugh) as well as plots T041 and T042 both located on the same site (Moor House - Upper Teesdale): these pairs of plots are therefore not interchangeable with other plots. In addition, 5 plots (904, 905, 906, 909, 910) include six to eight abundance data in some months (respectively in 3, 1, 2, 6 and 1 months), and five or less in all other months. This is not consistent with the methodology detailed in InsectChange (p. 17 of the detailed presentation), describing that the temporal resolution was between the week and the year, except in six datasets where plots were sampled between six and eight times in any month. For example, for plot 501 in 1996, the original dataset contains abundance counts on July 5, 8, 10, 15, 18, 22, 25 and 29, all of which are kept in InsectChange, without averaging per week and without indicating the dates of sampling. Also, there are several samplings in most months but InsectChange does not allow to know their chronology because it does not report the sampling date, which was available in the original dataset. The precise sampling areas in the different sites of the UK Environmental Change Network data are not known from the original study. Finally, the estimation of the local cropland cover provided in InsectChange is inconsistent with satellite images, with an overestimation or an underestimation for some plots.

**Study 1266 (Rennie et al. 2018c) - ECN spittlebugs**

This observational study examines the dynamics of spittlebug populations in different sites in the UK. The precise sampling areas in the different sites of the UK Environmental Change Network data are not known from the original study. In addition, Plots 1654 and 1655 representing the Yr Wyddfa/Snowdon site and Plot 1656 representing the Cairngorms site have been assigned erroneous geographic coordinates, approximately 270 km and 450 km apart from the correct location of the sites, respectively. The estimation of the local cropland cover provided in InsectChange is inconsistent with satellite images, with an overestimation or an underestimation for some plots.

**Study 1267 (Rennie et al. 2018d) - ECN ground beetles**

This observational study monitors the dynamics of ground beetles in the UK, as part of the UK Environmental Change Network (ECN) programme. InsectChange includes 36 plots from this study. Each plot had ten pitfall traps operating through most of the years, from which samples were collected at variable intervals. The sampling effort varied among sites for two reasons: first, the number of trapping days per year varied among plots and years; second, for many trap collection dates, records from less than ten traps were available at each plot. Appendix S2 of InsectChange describes the methodology to correct for these changes in sampling effort. The number of sampling days was homogenised by several operations: keeping records from May to October; removing years with short sampling periods; removing the last two weeks of October in years with long sampling periods. Numbers corresponding to quality codes and not abundance were removed, and beetle abundance was averaged over the functioning traps for each plot per date to account for randomly missing traps within plots. InsectChange includes the sum of abundance numbers over the year for each plot and year. Although this methodology is broadly appropriate, there was an error in the assessment of the number of trapping days, with Appendix S2 of InsectChange incorrectly describing that the sampling effort varied between 151 and 160 sampling days per year, with 87.79% of plot-years having a sampling effort of 154 days. This is because this trapping period was calculated from the day of collection of the first sample of each year and plot, instead of the day the trap was set. Instead of 154 trapping days, most combinations of plot and year actually corresponded to 168 trapping days. We found inconsistencies in the inclusion or exclusion of data and errors in insect counts. For example, for plot 1595 (T06-1 in the original dataset), the year 2004 was included although it covered only 164 trapping days (from May 14 to October 25), while the year 1995 covering 167 trapping days (from May 15 to October 30) was not included. For Plot 1590 (T04_2), the year 1994 covering 167 days (May 05 to November 02, without sampling from July 27 to August 10) was included with a correct insect count; for Plot 1584 (T02_2), the year 2003 covering 167 days (from May 08 to October 22) was included with an error in insect count, reported as 7.2 instead of 7.8. Finally, plots T04-4, T12-4, T12-5 and T12-6 from the original study of Rennie et al. (2018d) were not included in InsectChange although they respectively covered periods of 14, 10, 9 and 9 years, with adequate numbers of trapping days. These exclusions are arguably appropriate for plots T04-4 and T12-6 which included only few positive abundance records, but plots T12-4 and T12-5 included sufficient data to be considered. The precise sampling areas in the different sites of the UK Environmental Change Network data are not known from the original study. Finally, the estimation of the local cropland cover provided in InsectChange is inconsistent with satellite images, with an overestimation or an underestimation for some plots.

**Study 1310 (Wolda 1992) - Panama homopterans**

No identified problem.

**Study 1312 (Wolda et al. 1994) - Czech moths**

This natural experiment compares the dynamics of moth populations at four sites with different land uses in Czech Republic, three of which are included in InsectChange, one urban, one in a ruderal/deteriorated agricultural setting and one on a wet forest environment, but these characteristics of sites are not reported in InsectChange. The geographic coordinates of Plot 28 (in Brno) are inadequate for a study at the local scale, as they are 1.5 km apart from the Agricultural University where one of the authors maintained a light trap (the location of which is provided in the article).

**Study 1319 (Lightfoot 2010a) - LTER Seviletta grasshoppers**

This observational study monitored grasshopper species composition and abundance over a long period of time (from 1992 to 2013 for the site with the longest period of record) in four different sites of the Sevilleta National Wildlife Refuge with different habitat types (creosotebush shrubland, black and blue grama grassland, pinyon-juniper woodland), varying for their vegetal cover. The original dataset specified that portions of the Five Points Black Grama site (Plot 916 in InsectChange) were impacted by a natural fire in 2009. This fire disturbance, although it impacted the grasshopper abundance, was not indicated in InsectChange. The strata where grasshoppers were collected were soil surface or herb layer and the original dataset specifies the substrate - soil or a plant species - on which each grasshopper was observed. In InsectChange, all grasshoppers were inadequately assigned to the herb layer stratum. Studies 1319 and 1345 have overlapping data, as both include data on grasshoppers from the soil stratum in three identical plots of the Sevilleta LTER (Five Points Black Grama, Five Points Creosote and Cerro Montosa Pinyon-Juniper sites), obtained by visual counts in study 1319 and by collection in pitfall traps in study 1345. Moreover, there are two samplings for Plot 917 in the period "E" in 1992 but InsectChange does not allow to know their chronology because it does not report the sampling date, which was available in the original dataset.

**Study 1324 (Meijer and Barendregt 2018) - Netherlands beetles & spiders**

This observational study examines colonisation by ground arthropods in a tidal area that was reclaimed by a polder at the beginning of the study. According to the table *SampleData.csv*, the arthropods under consideration were "spiders and beetles and harvestmen + *Paederus* and 2 woodlice", however, while the first three are insects or arachnids, the last two are millipedes (myriapods) and woodlice (crustaceans) respectively, and thus neither insects nor arachnids. Third, the number of individuals of these arthropods was bound to increase in the first years in this newly created terrestrial area. The overall trend is declining in all sites of the study, but the overall declining trend is highly diminished by the high increase of arthropods in the first years of the study.

**Study 1328 (Hassall et al. 2017) - UK hoverflies**

This observational study examines the dynamics of hoverflies abundance in a single site in Leicester, UK, from 1972 to 2001. The "GroupInData" cell of the table *SampleData.csv* refers to "Hoverflies and bees", which is not consistent with the "InvertebrateGroup" cell of the table

*DataSource.csv*, the original publication by Hassal et al. (2017) and the description of the dataset obtained from raw data provided by the owner in Appendix S1 of InsectChange, all three referring to "Hoverflies". Finally, there are four samplings per month but InsectChange does not allow to know their chronology because it does not report the sampling date.

**Study 1335 (Honek et al. 2014) - Czech ladybeetles**

This natural experiment studies changes in coccinellid communities for areas with different land uses between two periods, a first period, 1976-1983, characterised by maximum agricultural intensification and a higher diversity of crops, and a second period, 2002-2010, characterised by a lower proportion of arable land, a lower use of chemical inputs and a lower crop diversity, as well as the spread of the non-native coccinellid species, *Harmonia axyridis*. The observation periods were inappropriately designated as 1983 and 2010 instead of taking the average dates between the two extreme years as in other studies. Insect abundance numbers were not accurately reported from Table 1 of Honek et al. (2014): in site 25 (cereals), abundance numbers decrease from 3678 to 1244 in the article, but only from 3677 to 1246 in figures reported in InsectChange; in site 26 (herbaceous plants), abundance numbers decrease from 1634 to 1415 in the article, but only from 1633 to 1415 in InsectChange. The publication specifies that in the site named "trees" (Plot 27), coccinellids were sampled in trees (and not in the herb layer as assigned in the stratum cell of the table *InsectAbundanceBiomassData.csv*). Finally, a unique pair of geographic coordinates was inadequately assigned to three different plots referred as "cereals", "herbaceous plants" and "trees", while the source publication specifies that habitats were sampled at various locations (31 locations in the first period, 22 in the second period) in lowland and submontane landscapes in the west of the Czech Republic. For this reason, the estimation of the local cropland cover in InsectChange is incorrect.

**Study 1339 (Belovsky 2018) - Montana grasshoppers**

This observational study examines the dynamics of grasshoppers in a bunchgrass prairie ecosystem in the National Bison Range in Montana (USA). The geographic coordinates of Plot 427 are given in the source study (Belovsky, 2018) and are 400 m distant from those used in InsectChange. In addition, the local cropland cover in InsectChange is overestimated compared to satellite images.

**Study 1340 (Valtonen et al. 2017) - Hungary moths**

This natural experiment examines the dynamics of moth abundance in seven sites differing in land use (different combinations of forest, agricultural and urbanised areas) in Hungary from 1962 to 2009. Moreover, the geographic coordinates assigned to Plot 152 in InsectChange were retrieved from Google maps and are inadequate, as Google Earth images around these geographic coordinates do not match the map of land use types around the trap in Figure S4 of the source study by Valtonen et al. (2017). For this reason, the estimation of the local cropland cover in InsectChange is incorrect for this plot. For two other plots, it is underestimated compared to information from the source study and satellite images.

**Study 1345 (Lightfoot 2010b) - LTER Sevilleta arthropods**

This observational study examines the dynamics of ground-dwelling arthropods at four sites of the Sevilleta National Wildlife Refuge with different habitat types (creosotebush shrubland, black and blue grama grassland, pinyon-juniper woodland) from 1992 to 2004. InsectChange includes three plots with sufficient duration of records. The methodology described in Appendix S2 of

InsectChange only corrected partially for changes in sampling effort, by taking into account, at each site, the five replicate lines A to E, and in each line, only traps 1, 3 and 5 that were sampled consistently from 1992 to 2004. Abundance values were then summed up across all sampling months and traps per plot per year. Two other sources of change in sampling effort were not accounted for in InsectChange. First, there are missing values in the original dataset, because some traps were often omitted from data tabulation due to individual traps being disturbed by precipitation runoff or vertebrate animals (see Welti et al. 2021, supplementary information). By summing up data across traps, InsectChange incorrectly considered these missing values as zeroes. Second, the traps remained opened all year and were collected approximately every two months, but the date of the last trap collection was variable depending on years and sites, and the last available data in a year varied from October 17 to December 21. This change in sampling effort due to the variable duration of sampling per year was not accounted for in InsectChange. Furthermore, studies 1319 and 1345 have overlapping data, as both include data on grasshoppers from the soil stratum in three identical plots of the Sevilleta LTER (Five Points Black Grama, Five Points Creosote and Cerro Montosa Pinyon-Juniper sites), obtained by visual counts in study 1319 and by collection in pitfall traps in study 1345.

**Study 1346 (Lightfoot 2010c) - LTER Seviletta ants**

This Small Mammal Exclusion Study (SMES) is a controlled experiment that compares the dynamics from 1995 to 2005 of the number of seed-harvester-ant nests (which are not insects *sensu stricto*) at two sites each with three plots that received different treatments. The three plots correspond to the lagomorph exclusion plot (L), the lagomorph&rodent exclusion plot (R) and the control plot (C), respectively. The treatments are incorrectly mentioned since the R treatment was reported in InsectChange as a rodent exclusion treatment instead of a lagomorph&rodent exclusion treatment. The R treatment had a strong positive effect on the trend of the harvester ants, and resulted in an increase on ant nests (a possible explanation is the removal of rodent competition pressure on harvester ants for seeds; however, this is not discussed in the dataset by Lightfoot et al. (2010c)). This was especially the case in the Rio Salado creosote site, where ant nests were the most numerous. Finally, this study does not consider ant individuals, but ant nests, in contrast with some other studies in InsectChange. The table *SampleData.csv* of InsectChange refers to "Ants", while it would be more appropriate to refer to "Ants (nests)". It would also be appropriate to specify the ant species that were selected by the source study (i.e. all *Pogonomyrmex* species, *Aphaenogaster cockerelli*, and *Myrmecocystus* species; *Pheidole* species were not selected) in the table *SampleData.csv*.

**1347 (Magnuson et al. 2010) - LTER North Temperate Lakes benthic arthropods**

This observational dataset from a long-term monitoring program of the LTER North Temperate Lakes examines the dynamics of benthic macroinvertebrates in different water locations in primary lakes of North America. InsectChange includes insects, arachnids and a small proportion of springtails (Collembola, Entognatha), which are neither insects nor arachnids (and not "freshwater fauna" as inadequately mentioned in the table *SampleData.csv* of InsectChange). Arthropod counts in 2016 and 2017 were available in the original dataset (link https://portal.edirepository.org/nis/mapbrowse?packageid=knb-lter-ntl.11.28 provided in the table *DataSources.csv* of InsectChange) but were not included in InsectChange, which stops in 2015. There is an error in arthropod counts as van Klink and coauthors considered a zero arthropod count for some taxa that were present but not counted (data flag "N") in Crystal Lake site 6 (plot 844) in 1982, 1985 and 1987, site 9 (plot 845) in 1985, site 43 (plot 843) in 1987, and site GN (plot 846) in 1987. This could have been accounted for by excluding the corresponding years for these sites, and therefore excluding site 844 for which the data then only cover six years. Finally, InsectChange included 14 sites on three lakes and inadequately assigned one pair of geographic coordinates per lake, while different plots on a same lake were up to 3.5 km apart.

**Study 1349 (Grimm and Childers 2018) - LTER Arizona pitfall-trapped arthropods**

This observational study examines the abundance dynamics of ground-dwelling arthropods at different sites in the Phoenix metropolitan area and the Sonoran desert region. The original dataset includes a “trap_count” cell recording the total number of traps collected and processed for a given site and a given sampling date that summarises changes in sampling effort. Appendix S2 of InsectChange states that dates with fewer than the full complement of traps were removed and that the sum of all traps per date was taken. Contrary to this statement, InsectChange does not account for changes in sampling effort. For example, for Plot 601 in InsectChange (site AA-17 in the original dataset), InsectChange includes abundance counts on 54 different dates, of which the sampling effort was 10 traps on 44 dates, nine traps on four dates, eight traps on three dates, seven traps on two dates. For this site, five sampling dates with 10 traps sampled were not included in InsectChange. The table *SampleData.csv* inadequately refers to “ground dwelling arthropods” while only arachnids, insects and springtails (Collembola, Entognatha), which are neither insects nor arachnids, were included. In addition, there are several samplings in four months but InsectChange does not allow to know their chronology because it does not report the sampling date, which was however available in the original dataset. Finally, the estimation for the local cropland cover provided in InsectChange is inconsistent with satellite images, with an overestimation or an underestimation depending on plots.

**Study 1351 (Grimm et al. 2007) - LTER Arizona freshwater arthropods**

This interrupted time series natural experiment from the Central Arizona-Phoenix Long-Term Ecological Research examines post-flood recovery in a desert stream ecosystem affected by ten flood events during the observation period, but this factor is not mentioned in InsectChange and the data of the source study on the number of days past flood at each sampling date is not included in InsectChange. The table *SampleData.csv* inadequately refers to "freshwater fauna", while InsectChange only includes insect and arachnid taxa of the original dataset. There are several samplings per month but InsectChange does not allow to know their chronology because InsectChange does not report the sampling date, which was however available in the original dataset.

**Study 1353 (Ernest 2018) - LTER Portal ants at baits**

This dataset is from a controlled experiment (LTER Portal), which deals with ant abundance (at given baits) in the Chihuahuan desert ecosystem near Portal, Arizona in 22 experimental plots and two control plots. In InsectChange, only the two control plots are considered, thus we did not classify it as a controlled experiment (but the two control plots are only poor representative of natural conditions for ant colonies since a set of experimental treatments (such as eliminating ants using poison) were applied to the 22 nearby plots and a surrounding fence protected the 24 plots from livestock). Studies 1353 (abundance of ants) and 1445 (number of nests) are both part of the Portal LTER and both include exactly the same two sites, which are two unmanipulated control plots from the Portal LTER. Therefore, it would have been appropriate to select only one of the two studies.

**Study 1357 (Schowalter 2011) - LTER Luquillo canopy arthropods**

This controlled experiment examines the responses of canopy arthropods to a canopy trimming experiment designed to separate the effects of canopy opening and debris pulse (resulting from hurricane disturbance) in a tropical rainforest ecosystem. Plots were subject to four different types of treatments: (1) canopy trimmed with debris removed, weighed, then redistributed throughout the plot to simulate conditions created by natural hurricanes; (2) canopy trimmed, with trimmed material removed from the plot to simulate canopy opening without debris deposition; (3) canopy undisturbed, with trimmed material from treatment 2 weighed, then distributed throughout the plot to simulate debris deposition without canopy opening; (4) canopy undisturbed and no debris alterations occurred at the forest floor. These experimental conditions are not reported in InsectChange in a comprehensive way, but only in the form of codes. In addition, the plots where arthropods were sampled did not correspond to geographic locations but to individual trees, up to seven per site on six different sites. The same geographic coordinates were assigned to these six sites, while they were distant up to 700 metres, and their individual geographic coordinates were available in the data paper. InsectChange includes insects, arachnids, as well as springtails (Collembola, Entognatha), which are neither insects nor arachnids (and not "all arthropods" as incorrectly mentioned in the table *SampleData.csv*). In this study, the trends of these arthropods show important variations depending on sites and no clear trend emerges. Moreover, there was an error in the counts of these arthropods for four sites (1008, 1013, 1028, 1036 corresponding to A_1_MICO, B_1_DACR, B_2_CECR, C_2_CECR, respectively) for which a zero count was not accounted for in June 2009, June 2004, July 2005, July 2007 respectively. Finally, among the original sites, two sites (B_1_CECR, C_4_PRES) showing a decrease in counts (with a zero count at the last recording date) were not selected while C_4_PRES had a 10-year duration for records and B_1_CECR had a 9-year duration (but other sites with a 9-year duration were selected in this study [e.g. C_4_MANI / Plot 1039]).

**Study 1361 (Pennings 2016) - LTER Georgia coastal grasshoppers**

This observational study examines the dynamics of grasshopper abundance in the Georgia Coastal Ecosystems (GCE) LTER located along three adjacent sounds on the Atlantic coast and including both intertidal marshes and estuaries. It is from a natural experiment whose specific goal was to characterise the responses of these insects in three dominant habitats to variations in salinity and inundation (see: https://lternet.edu/site/georgia-coastal-ecosystems-lter/). As the data on salinity and inundation are not available in the source dataset, we coded it as observational. Trapping effort was standardised by using only the first eight transects per plot (and not the first nine transects per plot as indicated in Appendix S2 of InsectChange). On the study period (2001-2016), Plot 632 (site 2 of the original study) has 0 abundance counts each year, except for a 1 abundance count in 2003. It would have been preferable to exclude this site, in line with the exclusion of sites with almost no insect counts in other studies (such as in study 1499).

**1364 (Tilman et al. 2006) - LTER Cedar Creek herb-layer arthropods**

This controlled experiment examines the effects of plant biodiversity on the dynamics of insects and arachnids sampled by sweepnet from 1996 to 2006. The table *SampleData.csv* inadequately refers to “all arthropods” when sampled communities were herb-layer insects, arachnids and springtails (Collembola, Entognatha), which are neither insects nor arachnids. This dataset includes 172 experimental grassland plots, each measuring 9 m x 9 m, in the reduced spatial scale of a 260 × 270 square metre field, each considered as a site in InsectChange. Six different controlled experimental treatments were different numbers of plant species planted in the plot (0, 1, 2, 4, 8,

16). These treatments carried out in each of the 172 plots introduced heterogeneity in terms of biodiversity, which artificially affected insect counts. Finally, the local cropland cover is overestimated as the experimental plots are coded as croplands instead of grasslands.

**Study 1365 (Gandhi et al. 2011) - Minnesota ground beetles**

This natural experiment compares the evolution of carabid communities in three different remnant habitats (cottonwood stands, oak stands and an old field) in response to the development of a highly urbanised matrix. Neither the surrounding urbanisation nor the land cover is mentioned in InsectChange. A unique pair of geographic coordinates was assigned to the three plots, which actually correspond to different stands at different locations in the Twin Cities Army Ammunition Plant site, in Arden Hills, Minnesota, an area composed of approximately 249 ha of grasslands, 191 ha of marshlands and 80 ha of upland woods (from the source publication of Gandhi et al. 2011, and from Epstein and Kulman 1990). For this reason, the estimation of the local cropland cover in InsectChange is incorrect.

**Study 1367 (Pizzolotto et al. 2014) - Italy ground beetles**

This natural experiment compares changes in ground beetle assemblages at different altitudes below and above the treeline of the Dolomites (Italian Alps) as an effect of climate change and shows that changes vary depending on altitude. The same geographic coordinates were inadequately assigned to the six plots of the dataset, while they are up to approximately 5 km apart from Figure 1 in the source study of Pizzolotto et al. (2014).

**Study 1376 (Shieh and Yang 2000) - Taiwan freshwater insects**

This natural experiment compares the freshwater insect abundance between 1985-1986 and 1995-1996 and for four study sites on a stream, with two sites located upstream agricultural areas and two sites in agricultural areas, one of them also receiving sewage from hotels due to tourism. The same geographic coordinates were inadequately assigned to all four plots, although the most distant plots are more than 3 km apart. It shows that study sites in agricultural areas had poorer stream water and habitat quality. The tables *DataSource.csv* and *SampleData.csv* of InsectChange inadequately refer to "Freshwater invertebrates" and "Freshwater fauna" respectively, when the study deals with freshwater insects of the Ephemeroptera, Plecoptera, Trichoptera, Diptera and Coleoptera orders. The series has only two time periods with data.

**Study 1377 (Roubik 2001) - Panama bees**

The geographic coordinates reported in InsectChange for the unique plot of this study are inappropriate, as Google maps was used to assess the location of the plot, but without pointing to the precise sampling location indicated in the original publication, which is approximately 10 km distant from the InsectChange assigned location.

**Study 1378 (Grøtan et al. 2014) - Costa Rica butterflies**

The geographic coordinates reported in InsectChange for the unique plot of this study are inappropriate, as sampling locations were up to 1.5 km apart.

**Study 1379 (Grøtan et al. 2012) - Equador butterflies**

This observational study examines the dynamics of a tropical butterfly community in Amazonian Ecuador. The geographic coordinates reported in InsectChange for the unique plot of this study are inappropriate, as the latitude of the plot, given in the publication in the degrees/minutes/seconds coordinate format, was inadequately converted as 0.497306 instead of -0.497306 from the original study. The actual location of the plot (latitude: -0.497306, longitude: -76.374694) is approximately 110 km away from that indicated in InsectChange. In addition, the geographic coordinates reported in InsectChange for the unique plot of this study are inappropriate, as sampling locations were up to 1.4 km apart. The local cropland cover provided in InsectChange is overestimated compared to information from the source study.

**Study 1381 (Souza da Silva et al. 2015) - Brazil chironomids**

This interrupted time series natural experiment examines the dynamics of chironomids in a Brazilian lake in relation to different changes over time in trophic conditions and water characteristics (clear and turbid waters) that were associated with substantial changes in the main primary producers (macrophytes and phytoplankton). These conditions likely to affect the chironomid dynamics are not mentioned in InsectChange. While this study pertains to chironomids, the "InvertebrateGroup" cell of the table *DataSource.csv* refers to "Freshwater invertebrates", inadequately suggesting that the whole invertebrate assemblage was considered. From Figure 1 in Trindade et al. (2009), Lake Biguás on the campus of the Universidade Federal do Rio Grande in Brazil is located 500 m northwest of the location provided in InsectChange based on the approximate geographic coordinates provided in the source study.

**Study 1382 (Meserve et al. 2016) - Chile arthropods**

This interrupted time-series natural experiment examines the dynamics of arthropod abundance and biomass in a semiarid thorn scrub community in north-central Chile in relation to high rainfall events. Arthropods considered include non-insects and non-arachnids (Chilipoda, Diplopoda and

Malacostraca [Amphipoda and Isopoda]), from Table 1 in the source publication by Meserve et al. 2016).

**Study 1384 (White 1991) - New Zealand moths**

This observational study examines the dynamics of moth communities at two grassland sites differing for herb diversity. InsectChange only includes two time periods, 1961-1963 and 1987-1989 (reported as 1962 and 1988). InsectChange reports the pooled abundances of the two moth sites during these two time periods from Table 1 of White's source publication, while the change in moth abundance by site was available in Appendix 1 of this publication. The geographic coordinates reported in InsectChange are not adequate for a study at the local scale because the two sites, pooled in InsectChange, were 4 km apart.

**Study 1385 (Quintero and Roslin 2005) - Brazil dungbeetles**

This natural experiment compares dung and carrion beetle assemblages in non-fragmented forests, logging-induced clearcuts and fragmented forests in central Amazonia in two years, one just after the disturbance and the other about ten years after the disturbance has stopped. It aims at studying how assemblage structure evolves over time after forest fragmentation. In this study, on plot 137, the abundance is 239 in the first year of the study (1986). An inaccurate abundance of 634 was reported in InsectChange in the last year of the study (2000). It should be 347 to account for a change in sampling effort reported in the source publication (Quintero, 2015, Table 2: "*The sampling effort implemented in 2000 was three times higher than that of 1986, and absolute numbers are therefore not directly comparable across years*"): that is, the indicated abundance of 1041 for the clearcut habitat in 2000 should have been divided by three, as was done by the authors of InsectChange in the three other sites of the study. This error biases the insect change by overstating the insect increase in this study. Results show that insect abundance decreased between the two years of the study in the control non-fragmented forest habitat, while it increased in the formerly disturbed habitats (fragmented forests and clearcut), where insect recovery occurred as expected. Finally, four types of habitats (1-ha forest fragment, 10-ha forest fragment, matrix habitat between the fragments, continuous forest tract) were coded as four plots sharing the same pair of geographic coordinates in InsectChange, while each of them was sampled in three different sites (Cidade Powell, Colosso and Dimona), up to 18 km apart. The local cropland cover provided in InsectChange is overestimated from the description of sampling sites in the original publication.

**Study 1387 (Langlands et al. 2006) - Australia spiders**

This controlled experiment analyses the recovery of spider assemblages in the Great Victoria Desert after different regimes of experimental fire, comparing plots subject to fires occurring more or less early in the time of record, as well as their interaction with the precipitation regime. The experimental fires were not mentioned in InsectChange. InsectChange includes data on the mean abundance in the four sites, all burned once in the beginning of the study (the record begins in 1990; fires occurred in 1990 in one site and 1991 in the three other sites). These time series, with experimental fires in the beginning of the observations, favour insect recovery and increase. The inclusion in InsectChange of this dataset with experimental sites is questionable. Finally, from Figure 2 of the source publication by Langlands et al. (2006), the sites pooled together in a single plot in InsectChange were up to 1.5 km apart, therefore the geographic coordinates assigned to this plot in InsectChange are inadequate.

**Study 1388 (Bêche and Resh 2007, Resh 2018) - California freshwater invertebrates**

This natural experiment incorporates data from three regions in northern California to capture the recovery of freshwater invertebrates from rare events, such as prolonged and extreme droughts. The dataset only includes data from one of these three regions, because in the other two regions the data covers less than ten years. Four sites differing in their flow regimes were considered and observed from 1984 to 2003. The geographic coordinates of the four plots in Knoxville and Hunting creeks are given in the source study by Bêche and Resh (2007). InsectChange assigned incorrect geographic coordinates to the four plots, distant from approximately 350 m to nearly 2 km from the actual geographic coordinates depending on plots. The extreme droughts occurred in the beginning of the observation period and lasted from 1987 to 1992 but they are not mentioned in InsectChange. The table *SampleData.csv* inadequately refers to "Freshwater invertebrates" while only insects and arachnids have been selected from the original invertebrate communities.

**1391 (Rybalov and Kamayev 2012) - Russia soil fauna**

This natural experiment compares the dynamics of soil fauna on six different sites along an elevation and ecological gradient ranging from forest to tundra ecosystems, but the characteristics of the different sites are not reported in InsectChange. The same geographic coordinates were inadequately assigned to these six sites, which locations are not specified in the source study. The "GroupInData" cell of the table *SampleData.csv* inadequately refers to "macroinvert group", while only ground insects and arachnids were included in InsectChange. There is an error in the insect stratum since the insects (including coleopterans, dipterans, lepidopterans, hymenopterans) and Araneae were sampled using pitfall traps and therefore were from the soil surface and not from the underground.

**Study 1392 (Kocíková et al. 2014) - Slovakia butterflies**

This observational study examines the change in butterfly community following the abandonment of a former pasture, no longer maintained and in the process of overgrowing with shrub vegetation. But this land use change is not indicated in InsectChange. The abundance and diversity of butterflies were shown to increase then decrease and the source article concluded that moderate grazing would provide heterogeneous disturbance of herbal level, as well as soil cover, i.e. a varied supply of microhabitats that many butterflies need for their living. From satellites images, the local cropland cover provided in InsectChange is overestimated.

**Study 1393 (Shafigullina 2009) - Russia island insects**

This natural experiment aims at analysing the impact of floods on insect abundance on the island banks of an artificial reservoir, by comparing three habitats differing in the degree of spring-time flooding: willow scrubs (flooded above 53 m almost every year); middle-level meadows (flooded above 53.3 m some years); high-level meadows (never flooded). The article by Shafigullina (2009) shows a positive trend of insect communities observed in the study sites, first due to a recovery of insects after a catastrophic flood in the second year of the observation period (1979). This initial high flood disturbance and the subsequent flood cycle are not specified in InsectChange. The positive trends are also interpreted as an adaptation of insect communities to flood conditions (migration to higher areas prior to overwintering) and to higher temperatures. InsectChange includes three plots, with some overlap between them: plots 129 and 130 (defined from Figure 2) are respectively willow scrubs and (middle-level + high-level) meadows; while plot 131 (defined from Figure 3) includes regularly flooded habitats, that is, willow scrubs and middle-level meadows. Therefore, willow scrubs are included in both plots 129 and 131, while middle-level meadows are included in both plots 130 and 131. Moreover, there are errors in insects counts in all plots. Indeed, data for plot 129 are extracted from Figure 2, which focuses on herb-layer insects and arachnids captured by a biocenometer, by summing data on all these ‘chortobiont’ invertebrates with detailed data on some of them (Heteropterans, Coleopterans, Lepidopterans and parasitoid wasps), which are therefore counted twice. The same error applies to plot 130, which data are also extracted from Figure 2. Data for plot 131 are extracted from Figure 3, which focuses on soil surface and herb-layer Coleopterans respectively captured by pitfall traps and sweep nets, by summing data for all species with data for species with overwintering preimaginal stages, which are thus counted twice. There are also errors in identities of insect groups (cell “GroupInData” in the sample data of InsectChange): it indicates that the insect groups from Figures 3a, 3b and 3c are “insects” while contrary to those of figure 2 which include several insect taxa, each of these three groups refers to a particular and distinct taxon, coleopterans of the Carabidae, Chrysomelidae and Curculionidae families respectively. In addition, the SampleDate file only provides information at the study level and does not allow to infer which insect groups are included in which plots. This information is not given in any of InsectChange files. Also, the geographic coordinates reported in InsectChange for this study, without any source being provided, are inappropriate, as the original publication describes that the material was collected in several islands of a 6,450 km² wide reservoir. Finally, the local cropland cover provided in InsectChange is inconsistent with information from the source study and is overestimated.

**Study 1394 (Valtonen et al. 2013) - Uganda butterflies**

This observational study examines the seasonal and inter-annual variation in tropical fruit-feeding butterflies in the medium altitude montane rainforest of Kibale National Park, Uganda. The geographic coordinates of the Makerere University Field Station given in the article are approximate and 3.8 km distant from the location indicated on Google Earth. The article describes that 22 butterfly traps were placed in the forest understory at locations 100 m apart in closed canopy forest, implying that the local cropland cover is 0%, and not 100% as assessed in InsectChange.

**Study 1395 (Crosa et al. 2001) - Africa freshwater invertebrates**

This interrupted time series natural experiment examines the effect of a larvicide treatment used to control the blackfly *Simulium damnosum*, vector of the parasitic worm, *Onchocerca volvulus*, in three Guinean rivers. The observation spans from 1984 to 1998 and the rivers were treated with larvicides from 1987 to 1998 but this treatment is not mentioned in InsectChange. The dataset contains the average over the three rivers of the abundance data of all invertebrates, including non-insects, while data was available on four insect taxa (Tricorythidae, Leptoceridae, Philopotamidae and Hydrospychidae: see Figure 3 in Crosa et al. 2001). The source study describes that samples were collected in three Guinean rivers, therefore the geographic coordinates of the average plot included in InsectChange are not adequate to describe drivers at the local or landscape scale.

**Study 1396 (Daghighi et al. 2017) - Germany springtails**

This controlled experiment pertains to springtails (Collembola, Entognatha), which are neither insects nor arachnids. It examines their succession dynamics in relation with the vegetal cover and thereby soil temperature in two experimental plots on a no longer exploited rubble and debris dump in Bremen, Germany. One plot was left undisturbed for natural secondary succession and the other recultivated with rotary-tilling and sowing of grass. Both experimental plots show a decreasing trend of springtail abundance. Finally, the positive local cropland cover provided in InsectChange is overestimated compared to information from the source study.

**Study 1397 (Gallé 2017) - Hungary ants**

This experimental study examines the long-term dynamics of ant colonies over 37 years on a sandy grassland in central Hungary, a region that has been highly affected by climate change and has gradually turned into a semi-desert. It examines ant community assemblage under experimental slates placed on two types of micro-habitats, small dunes with dry sandy soil and sparse vegetation, and between-dune slacks with more humid soil and denser vegetation. InsectChange includes the mean evolution of the number of ant nests on both types of micro-habitats, which shows an increasing trend. However, this increasing trend results from the experimental conditions, that are not mentioned in InsectChange, and cannot be interpreted regardless of these experimental conditions. Indeed, the slate plates were placed as artificial nesting sites in the first year of study and the source study indicates that the low number of nests in the first year presumably results from the disturbance caused by laying the slates, that ants colonised the soil under the slates, and that the ant colonies presumably survived for several years and new ones were also established during the study period. The source study does not address the evolution of the number of nests, but the evolution of the proportions of the different species in the assemblages. Neither the gradual drought nor the microtopography in the experimental design affecting ground temperature and humidity are mentioned in InsectChange. The source study concludes that "[c]ommunity level transformations induced by climate change and especially ground water depletion are leading to gradual rearrangement of communities, homogenisation of species composition across habitat types and a considerable decline of diversity, threatening the fauna of the whole region". Comparison over time and between microhabitats of temperature and water table level allowed the authors of the source publication to suggest that drying of the water table was the primary factor affecting such a change in community structure. The "GroupInData" cell of the table *SampleData.csv* indicates "Ants", while it would be appropriate to refer to "Ant (nests) under experimental slates". Finally, the local cropland cover provided in InsectChange is overestimated given information from the source study and the satellite images.

**Study 1398 (Hodecek et al. 2015) - Czech beetles**

This natural experiment compares the dynamic of beetle communities in post-industrial sites (ancient spoil heaps) depending on the number of years that had passed since spoil depositing had ceased but these conditions are not specified in InsectChange. Appendix S2 of InsectChange specifies that “Because of unequal sample sizes, all species counts in 1975 and 1976 were multiplied by 5/7. In these years there were seven traps installed, whereas there were five in other years”. Actually, from Hodecek et al (2015), p 632, there were eight traps in 1975 and 1976 (Bezruk site, i.e. plot 117, and Zarubek site, i.e. plot 119). Therefore, the abundance levels for these years should have been multiplied by 5/8. The abundance levels are therefore overestimated in 1975 and 1976, which biases the dataset towards overstating insect decrease.

**Study 1400 (Ananin and Ananina 2011) - Russia beetles**

The geographic coordinates reported in InsectChange are inappropriate, as there are several sampling points in each of the four areas of the study, considered as four plots, and these sampling points are more than 900 m apart (the information in the original study not enabling to calculate the distance between sampling points in each area considered as a plot in InsectChange).

**Study 1401 (Tsurikov 2016) - Russia beetles 2**

This observational study examines the dynamics of beetles in different habitats in the Russian Galichya Gora natural reserve. InsectChange includes four plots inadequately sharing the same pair of geographic coordinates, while the source publication (Tsurikov, 2016) describes that insects were collected in several locations, in grass stands of steppe, oakery, and meadow. For this reason, the 100% local cropland cover provided in InsectChange is incorrect and overestimated.

**Study 1402 (Babenko 2013) - Russia springtails**

This observational study investigates the dynamics and composition of the assemblages of springtails (Collembola, Entognatha), which are neither insects nor arachnids, for sites with different plant communities in the Taimyr Peninsula in Russia. As specified by Babenko (2013), in this study including only two years, the sampling method was improved between the first year of the series, 1969, and the last year of the series, 2010. As a result, absolute abundance values cannot be compared between 1969 and 2010. Therefore, the inclusion of this study in the dataset is not appropriate. In the eight plots included in InsectChange, absolute abundance values increase from 1.6-fold to 5.5-fold. The inclusion of this series therefore biases the dataset by overstating the increasing insect trends. It should also be noted that fluctuations of springtail populations can be high in tundra conditions, and therefore a comparison of only two years of data cannot be straightforwardly interpreted. A unique pair of geographic coordinates was inadequately assigned to the eight plots, while the source publication indicates different locations of sampling (two variants of tundra (spotted or frostboil and hummocky), a polygonal mire, riverside meadows, and Dryas - fellfield associations).

**Study 1403 (Fedyunin 2008) - Russia parasitoids**

This observation study examines the population dynamics of Ichneumon parasitic wasps in the Visim Reserve in Russia from 1994 to 2003. InsectChange reports 64 abundance counts in specific days in 1994, 1997, 2002 and 2003 from Figure 1 of this publication. This is in contradiction with the methodology of InsectChange, describing that daily samples were summed or averaged to form a week or month (“Some data sets reported multiple data points within each year (days, weeks, months or seasons). To account for this, we maintained 'week' as the finest temporal grain, hence, data in data sets with a finer temporal grain than week were aggregated per week or larger temporal unit (e.g. daily samples were summed or averaged to form a week or month)”). For example, in July 2002, InsectChange reports separate counts on June 18, June 22, June 24 and June 28, these dates being mentioned in the “period” cell of the table *InsectAbundanceBiomassData.csv*, while the first two dates were in the 25th week of 2002 and the last two dates in the 26th week of 2002. Finally, the geographic coordinates reported in InsectChange are inappropriate as the source study does not specify the sampling locations within the Russian Visim nature reserve, which covers more than 300 $km^2$.

**Study 1404 (Aarhus University 2018) - Greenland arthropods**

This observational study monitors the evolution of arthropod communities in different sites in Greenland. This dataset is not included in the table *InsectAbundanceBiomassData.csv*, because its access license precluded publication of derived numbers, but its meta-data are included. The reference to “all arthropods” in the “GroupInData” cell of the table *SampleData.csv* is not consistent with the statement in Appendix S2 of InsectChange that taxa not belonging to the classes Insecta, Arachnida or Entognatha were excluded. In addition, this meta-data includes Entognatha, which are neither insects nor arachnids. Furthermore, the same geographic coordinates were inadequately assigned to six plots, up to approximately 1 km away from each other.

**Study 1405 (Martikainen and Kaila 2004) - Finland saproxylic beetles**

No identified problem.

**Study 1406 (Nemkov and Sapiga 2010) - Russia steppe arthropods**

This study examines the effect of catastrophic fires on ground-dwelling arthropods in a protected steppe ecosystem. InsectChange includes data from 1990 to 2004 in one site of the original study, the Burtinskaya Steppe in Russia. The geographic coordinates of the unique plot in InsectChange are inadequate as arthropod sampling was performed in different locations of the Burtinskaya Steppe, which has an area of 45 $km^2$ according to Dusaeva et al. (2019). The local cropland cover is overestimated given information from the source study. According to the authors of the original study, in 1998 and 2003, "catastrophic fires of very high intensity destroyed almost the entire protected area". The 1998 fire occurred in the summer, when arthropods were in full activity, and greatly reduced arthropods abundance, in contrast to the 2003 fire, which occurred in the fall when arthropods were diapausing, and thus had a less negative impact on arthropods. Thus, this study essentially shows arthropod recovery after a major fire, but this extreme disturbance factor was not mentioned in InsectChange. Moreover, the terrestrial arthropods trapped in this study include not only specimens of the classes Insecta and Arachnida but also specimens of the classes Myriapoda and Crustacea (woodlice). In addition, these ground-dwelling arthropods were inadequately referenced as merely arthropods in the table *SampleData.csv*.

**Study 1407 (Korobov 2015 - Russia beetles 3)**

This natural experiment compares the dynamics of ground beetle communities in an intact spruce forest and in a spruce forest impacted by windfall. It shows that the windfall created a decrease of the population diversity, followed by a recovery, but the windfall is not mentioned in InsectChange. The geographic coordinates reported in InsectChange are inappropriate as the source study specifies that sampling occurred in the Central Forest Nature Reserve in Russia but does not specify the sampling locations within this reserve, which covers more than 210 km².

**Study 1408 (Huttunen et al. 2017) - Finland freshwater insects**

This natural experiment examines the role of habitat connectivity and in-stream vegetation on the dynamics of benthic insect communities in 23 sites, but the connectivity of habitats and their vegetation cover were not taken into account in the dataset. The 23 plots in InsectChange inadequately share the same geographic coordinates, while they are 23 different streams in the Finnish part of the Koutajoki drainage basin in northeastern Finland, just south of the Arctic Circle. Many of them are located within a Nature conservation Reserve, Oulanka National Park, which represents the westernmost remnants of pristine taiga forests. All the streams drain mixed forests and bogs with minimal anthropogenic impact, less than ten percent of their catchments being modified by any land use activities (mainly forestry). An extreme drought in 2006 impacted all insect assemblages but was not mentioned in InsectChange. The tables *DataSource.csv* and *SampleData.csv* of InsectChange inadequately mentioned "Freshwater invertebrates" and "Freshwater fauna" respectively, whereas InsectChange included freshwater insects, except for chironomids. Indeed, chironomids were excluded from the dataset of the original study because they were not counted every year. This makes this dataset unrepresentative and very different from the other freshwater datasets since chironomids usually make up more than two thirds of freshwater insect assemblages.

**Study 1409 (Hallmann et al. 2017) - Germany flying insects**

This natural experiment examines the dynamics of the total biomass of flying insects caught with Malaise traps over 27 years in 63 nature protection areas in Germany, in relation with changes in weather conditions, land use and habitat at surroundings of the traps. These factors were found unlikely drivers of the major decline in aerial insect biomass observed in the investigated protected areas. The "GroupInData" cell of the table *SampleData.csv* of InsectChange indicates "all", while it would be appropriate to refer to "flying insects". Although it covered a 10-year period and thus met the inclusion criteria for the study, plot PLI1 from the original study of Hallmann et al. 2017 was not included in InsectChange. In this plot the annual average of daily insect biomass decreased by 70% between 2005 and 2014. Also, daily biomass insect counts were reported in InsectChange with a systematic error. For each plot, the source data provides biomass counts for trapping periods in days: for example, for Plot BIR1 of the original dataset, in 2000, the biomass 76.1 g was for the seven-day period extending from May 6 to May 12. Instead of calculating the corresponding daily biomass as 76.1 g divided by 7 days, that is, 10.87 g/day, it was inadequately reported as 12.68 g/day in InsectChange, corresponding to 76.1 g divided by 6 days, a time period inadequately shortened by one day. As a result, all insect counts are incorrect in InsectChange for this study and as the number of days between two collections of samples in traps is highly variable (ranging from 3 to 43 days), the error is not of the same magnitude among records and this affects the trend of insect abundance. Furthermore, there are several samplings in most months but InsectChange does not allow to know their chronology because it does not report the sampling date, which was however available in the original dataset. Also, given that abundance records were obtained by collecting insects in Malaise traps operating continuously but beginning or ending at different dates each year, it would have been more appropriate to sum data per year for each plot, after correcting for changes in sampling duration, as was done in other InsectChange studies relying on continuous trapping (studies 249, 1267 and 1345). Finally, the estimation of the local cropland cover provided in InsectChange is inconsistent with that from satellite images, with an overestimation or an underestimation depending on plots.

**Study 1410 (Karg et al. 2015) - Germany grassland above-ground insects**

This controlled experiment compares insect communities in reference sites and in grasslands restored by stopping fertilisation and reducing mowing in 1995-1996 over an observation period from 1992 to 2005. The date of cessation of agricultural practices is not specified in InsectChange. The nature of the experiment is incorrectly referred as "Oligotrophication" in the table *PlotData.csv* of InsectChange. Moreover, the table *DataSource.csv* refers to "Arthropods" and the table *SampleData.csv* to "All Arthropods" while the dataset deals with "Above ground insects" (mostly flying insects sampled with suction traps). Finally, the source publication indicated that the study was conducted at the Ecological Field Station of the Bavarian Academy for Nature Conservation and Land Care in Laufen, Germany, and that the actual sampling was carried out in the Schinderbach brook valley (which is located 2 km west from this institution according to Google Earth), without specifying the precise location of the sampling sites. The source publication provided inaccurate geographic coordinates for the Ecological Field Station in Laufen, Germany, pointing to a different town, Passau, 83 km away. InsectChange used these inaccurate geographic coordinates provided in the source publication. For this reason, the estimation of the local cropland cover in InsectChange is incorrect.

**Study 1411 (Steinwandter et al. 2017) - Austria soil arthropods**

This natural experiment examines soil community patterns caused by agriculture abandonment. It compares an intensively-managed meadow with an abandoned afforested meadow, and a managed pasture with an abandoned pasture. Insect recovery is thus expected after agriculture cessation.

Insect numbers are not accurately reported from Table 2 and the supplementary materials of Steinwandter et al. (2017). In Plot 10 (managed meadows), Plot 11 (abandoned meadows), Plot 12 (managed pastures) and Plot 13 (abandoned pastures), respectively, abundance numbers in 1998 and 2012 are 921.14 and 314.26 (and not 874 and 314.26 as indicated in InsectChange), 1978.33 and 711.39 (and not 1908.1 and 711.37), 458.37 and 427.45 (and not 444.23 and 428.43), 1813.67 and 318.3 (and not 1745.77 and 324.36); biomass numbers in 1998 and 2012 are 2114.05 and 1488.48 (and not 2115.87 and 1488.46), 3637.74 and 1472.6 (and not 3703.67 and 1472.39), 1075.01 and 2003.39 (and not 1069.71 and 2003.24), 3771.62 and 1420.55 (and not 3843.46 and 1398.47). Abundance numbers are for Aranae and insects, while biomass numbers are for insects only; the groups included are incorrectly referred as "All_soil_fauna" in the table *SampleData.csv* of InsectChange. Finally, the same pair of geographic coordinates was inadequately assigned to the four plots, while they were up to approximately 1.3 km apart.

**Study 1412 (van dam 2009) – Netherlands freshwater arthropods**

This observational study examines the dynamics of freshwater invertebrates over groups of years (i.e., abundance means for 1984-1987, 1988-1992, 1993-1997, 1998-2002, 2003-2007) in the north of the Netherlands. The data from this study covers 415 sites spread over an area of approximately 80 km by 45 km (see Figure 3.1 in the source study by van Dam, 2009). Therefore, the geographic coordinates for the unique plot in this study are inadequate for studies at the local or landscape scale. The observation periods were inappropriately designated as 1987, 1992, 1997, 2002, and 2007 instead of taking the average dates between the two extreme years as in other similar studies. The table *SampleData.csv* inappropriately refers to "Freshwater fauna" instead of "Freshwater insects and arachnids".

**Study 1413 (Kwon et al. 2016) - Korea soil arthropods**

This natural experiment compares the dynamics of soil arthropod communities in South Korean temperate deciduous and coniferous forests close and far from industrialised sites. The study includes non-insects and non-arachnids (Chilipoda, Symphila, Diplopoda, Crustacea, Entognatha) and on average, for all years and all sites, insects and arachnids represented 65% of the assemblage (see Kwon et al. 2016, supplementary material). The tables *DataSource.csv* and *SampleData.csv* of InsectChange refer to "Soil fauna" and "Arthropods" respectively, whereas it would be more appropriate to refer to "Soil arthropods" instead. The article does not provide data on the annual composition of the assemblage and therefore does not allow separation of taxa. The assemblage trend is decreasing in seven of the eight sites, mainly due to air pollution, acid rain and increasing heavy rainfall due to climate change. Finally, the estimation of the local cropland cover provided in InsectChange is overestimated given information from the source study and satellite images.

**Study 1414 (Gardarsson et al. 2004) - Iceland midges**

This observational study examines the dynamics of midge abundances in Lake Myvatn and the River Laxá (2 sites per location) in Iceland from 1977 to 1996. InsectChange includes data from the two sites at Lake Myvatn. The InvertebrateGroup cell of the Table *DataSource.csv* refers to "Mosquitoes" while the source study pertains to chironomids and simuliids.

**Study 1415 (Brunk et al. 2014) - Wisconsin mayflies**

This observational study is from a natural experiment, which investigates the dynamics of mayfly abundance inside and outside a Superfund zone contaminated with oils, tars, wood debris and wood treatment pollutants in the Chequamegon Bay (Lake Superior) to test for an effect of pollution. Outside the Superfund, the less polluted zone, 33 different plots varying for their substrates (clay, sand, ...) were also compared. InsectChange includes only a plot with the mean data from the 33 sites located outside the Superfund zone because only one year of data is available in the Superfund site but the DetailsPlots cell of the table *PlotData.csv* inadequately mentions “all sites”. The dataset only includes two years of data. A unique plot representing the mean of 33 plots was reported in InsectChange. The slight difference among years in mayfly abundance outside the Superfund site led the authors of the original study to suggest that contaminants from the Superfund site did not impact burrowing mayfly populations outside the Superfund boundary to any great extent. By contrast, the biomass of mayflies and its trend were shown to vary according to the dominant substrates of the different plots. Detailed data on biomass and abundance depending on the dominant substrate were available in Table 1 of the source study by Brunk et al. (2014) but were not included in InsectChange. The geographic coordinates of the unique plot considered in InsectChange are inadequate, as there were 33 sampling sites, up to 17 km apart (see Figure 1 in Brunk et al. 2014).

**Study 1416 (Holmes 2018) - LTER Hubbard Brook caterpillars**

This observational study investigates the 1986-1997 dynamics of caterpillar abundance on tree foliage of four tree species on the main bird plot in the Hubbard Brook Experimental Forest and on three additional plots within the White Mountain National Forest (US). InsectChange includes sampling on beech and sugar maple, the only tree species with at least 10 years of caterpillar sampling, summing the caterpillars collected on both tree species per plot and year. While the sampling effort varied from four to six sampling periods per year, this change in sampling effort was only partially corrected, as stated in Appendix S2 of InsectChange: “There was no consistent sampling in sampling period 6, and therefore we excluded this period”. However, there were only four sampling periods in 1987, while there were five sampling periods in all other years. This change in sampling effort was not corrected, leading to underestimate caterpillar abundance in the first year of observation, biasing caterpillar trends towards a higher increase or a lower decrease than the actual trend. Finally, the geographic location of the Hubbard Brook sampling site (Plot 639) is inadequately reported as the barycentre of the 44 $km^2$ area of the Hubbard Brook Experimental Forest bird inventory plots reported in the original dataset. The 2021 revision of the original dataset gives the bounding coordinates of the inventory plots, which are up to approximately 8.5 km apart (Zammarelli et al. 2022).

**Study 1417 (Stout and Rondinelli 1995) - Michigan freshwater insects**

This natural experiment investigates the impact on Michigan freshwater insect biomass of extremely low frequency electromagnetic fields (ELF) by comparing two sites on the Ford river, a site where an ELF antenna crosses the river and a reference site located 7 km downstream, relying on a BACI (Before and After, Control and Impact) procedure and a non-parametric randomised intervention analysis. These two plots inadequately share the same geographic coordinates in InsectChange. The original study finds that natural physical factors appear to be more important than the anthropogenic ELF fields in accounting for changes in the community. The tables *DataSource.csv* and *SampleData.csv* inadequately refer to as "Freshwater invertebrates" while the study deals with "Freshwater insects".

**Study 1418 (Crowley and Johnson 1992) -Tennessee dragonflies**

The geographic coordinates indicated in InsectChange are located at the top of a mountain bordering the lake, 900 metres from the lake. From the source study, sampling was performed in

9-12 stations within the five principal habitats in Bays Mountain Lake, therefore the unique pair of geographic coordinates used in InsectChange is not adequate.

**Study 1419 (Bisevac and Majer 1999) - Australia ants 2**

This natural experiment compares ant communities in rehabilitated and native heathland reference sites of a sand mining operation, in two years. InsectChange only includes the three rehabilitated sites that were sampled in the two years of records, with systematic errors in reported insect counts. Insect counts are correct in the first year but systematically higher than the actual data in the second year. For plot 6 (site 77AS of the original article), abundance is correctly reported as 1435 in 1980, but incorrectly reported as 277 instead of 166 in 1997. For plot 7 (site 77CS of the original article), abundance is correctly reported as 109 in 1980, but incorrectly reported as 128 instead of 101 in 1997. For plot 8 (site 78ES of the original article), abundance is correctly reported as 144 in 1980, but incorrectly reported as 164 instead of 139 in 1997. Besides, data for three control (reference) sites, with decreasing abundance, are not taken into account in InsectChange: from 1980 to 1997, abundance numbers decrease from 685 to 114 in site C1, from 331 to 276 in site C2, and from 389 to 222 in site C3. All these errors and omissions bias the insect trend of the dataset towards minimising its decrease. Finally, the same geographic coordinates were inadequately assigned to the three plots of the study, but the source publication specifies that the mining area where the study was conducted covered approximately 210 $km^2$; Table 1 shows that the three included sites (78 ES, 77 CS and 77 AS) have different environmental characteristics. For this reason, the estimation of the local cropland cover in InsectChange is incorrect.

**Study 1421 (Johnson and Harp 2005) - Arkansas freshwater invertebrates**

This natural experiment compares the dynamics of freshwater invertebrate assemblages of a downstream site in the tailwater of a dam and a control site located further downstream. The dataset contains only two years of observation, 7 and 35 years after the completion of the dam. The cold tailwater initially disrupted the existing macroinvertebrate assemblage (as specified in the abstract of the study), therefore the whole assemblage is expected to increase or stabilise, but not to decrease, but this is not specified in InsectChange. InsectChange reports the total abundance of invertebrates, including Isopoda, Turbellaria, Oligochaeta, Gastropoda and Amphipoda, while it was possible to dissociate Diptera and EPT from tables 1 and 2 of Johnson and Harp (2005). The same geographic coordinates were inadequately assigned to the two plots, located 4 km apart.

**Study 1422 (Smith et al. 2011) - Tennessee freshwater EPT**

This natural experiment studies the recovery of the macroinvertebrate community after pollution abatement in an industrially polluted stream. In this publication, three polluted sites (two very close to the pollution source, one more distant) that were subjected to three successive pollution reduction measures were compared with two control sites for their EPT (Ephemeroptera, Plecoptera, and Trichoptera) densities and assemblage structures. Van Klink and co-authors considered the time series of EPT community recovery from the three polluted sites and one "control site" which is actually not a site. This makes insect counts inadequate: its data was extracted from Figure 2c in Smith et al. (2011), using the middle of the grey shading in this Figure, which represented 95% confidence intervals for sample units from both reference sites from 1988 to 2003, but from only one reference site (Brushy Fork) in 1986 and 1987. The geographic coordinates assigned to this plot (Plot 111) in InsectChange on the base of Google maps data are inadequate: they correspond to the location of one reference site (BFK7), while the other reference site (HCK20) is located 20 km away.

**Study 1423 (Clements et al. 2010) - Colorado freshwater invertebrates**

This natural experiment examines the freshwater invertebrate recovery in a metal-polluted river that was decontaminated during the observation period. Two control sites are located upstream of the main source of contamination. Two other sites are located downstream of the main source of contamination where a water treatment facility was carried out during the observation period; tailings removal and revegetation of riparian areas were undertaken later in the observation period. From Figure 1 in the original publication, the correct locations of Plots 22 and 23 are 4 km and 14 km south of the location given in the publication, respectively. InsectChange reports the abundance of the whole macroinvertebrate assemblage, including non-insects (oligochaetes are mentioned in the article). It is not possible to distinguish insects in the data.

**Study 1425 (Grubaugh and Wallace 1995) - Georgia freshwater invertebrates**

This observational study reports the improvement in the condition of a stream, after a land-use change, i.e. abandonment of intensive agriculture having occurred before the observation period but this important factor susceptible to favour insect recovery was not specified in InsectChange. InsectChange includes the whole invertebrate assemblage. The study only includes two years of record. It was only possible to dissociate insects for the second year of the study. We calculated that in that year, insects accounted for 62% of total abundance and 61% of total biomass. The abundance data in Figure 4 of Grubaugh and Wallace (1995) were reversed between the two years in InsectChange: the 1956 data (hatched in the figure) were reported for 1991, and conversely, the 1991 data (shaded in the figure) were reported for 1956. InsectChange indicates "*Google maps.*

*Description in Nelson & Scott 1962" as the source of the geographic coordinates assigned to the unique plot of this study, but these geographic coordinates actually point to a location 8.5 km northwest of the one indicated in the article by Nelson and Scott (1962), specifying that "[t]he study area [...] was located on the Middle Oconee River, one-half mile upstream from U. S. Highway 29, near Athens, Clarke County, Georgia*".

**Study 1426 (McCreadie et al. 1994) - Pennsylvania blackflies**

This observational study examines the emergence of black flies (simuliids, inadequately referenced as "Freshwater invertebrates" in the Table *DataSource.csv*) in three sites of two first-order streams located in Lake Erie drainage basin differing in temperature regime (a stream with one site, and the other stream with two sites within close proximity). It finds that emergence patterns were synchronous in the two nearby sites of the same stream, but asynchronous between streams, with dissimilarities possibly reflecting divergent temperature regimes. The location of these sites in InsectChange does not correspond to their description in Figure 1 of Adler et al. (1982). Plot 115 only covers a 7-year period (1980-1986) and therefore its inclusion is in breach of the criteria set out in InsectChange and the statement in Appendix S2 of InsectChange that "[a]ll plots with fewer than 9 years of data were removed from the analysis".

**Study 1427 (Rugenski and Minshall 2014) - Idaho freshwater invertebrates**

This natural experiment compares the dynamics of freshwater invertebrate abundances and biomasses in four streams burned by a wildfire with two unburned control streams, with eight years of data before the wildfire and five years of data after the wildfire. Results suggest that the effects of climate warming mitigated the effects of fire, since the decrease of snow-melt runoff caused by climate warming buffered the adverse effect of the fire-mediated runoff from sparsely vegetated uplands. This resulted in increase in basal food resources and invertebrate density and biomass following fire. Abundance and biomass data reported in InsectChange refer to the whole invertebrate assemblage and it was not possible to distinguish insects from non-insects in the source article. However, the dataset by Baxter et al. (2016) on the same streams mentions the presence of Mollusca, Crustaceans, Oligochaeta, Turbellaria, Nematoda, Collembola and Hydrozoans (https://doi.org/10.2737/RDS-2016-0027). A single pair of geographic coordinates, corresponding to the Google maps location of the Frank Church River of No Return Wilderness, was inadequately assigned to the six plots representing six different streams. Furthermore, according to Table 1 of the original study by Rugenski and Minshall (2014), the six sample sites were at elevations of 1095-1220 m, whereas the geographic coordinates used in InsectChange were at an elevation of 2555 m according to Google Earth. Studies 1427 and 1437 share the same pair of geographic coordinates in the table *PlotData.csv* of InsectChange, corresponding to the Google maps location of the Frank Church River of No Return Wilderness, although they cover each a different set of streams. These studies share the same methodology of comparing burned and unburned sites, and present results with identical metrics of density (number/m²) and biomass (mg/m²) of the macroinvertebrate assemblage, in different periods (1979-1989 for study 1437; 1993-2005 for study 1427), but while for study 1427 both metrics were selected in the table *InsectAbundanceBiomassData.csv* of InsectChange, for study 1437 only the biomass data was selected.

**Study 1428 (Bradt et al. 1999) - Pennsylvania freshwater macroinvertebrates**

This interrupted time-series natural experiment examines the stability and resilience in benthic macroinvertebrate assemblages in a US stream surrounded by agricultural and residential zones and disturbed by a series of floods and severe winters. These studied disturbances occasionally impacting the invertebrate abundances are though not mentioned in InsectChange. The source article shows that despite severe winter conditions and floods, the assemblage recovered and exhibited both stability and resilience. InsectChange includes the abundance and biomass of the entire benthic macroinvertebrate assemblage. It was actually possible to separate abundances counts of Ephemeroptera, Trichoptera and Diptera insects, which represent a highly variable share of the total assemblage. The geographic coordinates are not appropriate for a study at the local scale given that from Figure 1 of the original publication, the study area extended over 2.5 km.

**Study 1429 (Rudstam 2018) - New York freshwater insects**

This observational study examines the dynamics of benthic invertebrate densities in Oneida Lake, New York. InsectChange includes only insects extracted from the original dataset. Therefore, the table *SampleData.csv* inadequately refers to “freshwater fauna” instead of “freshwater insects”. There are neither Arachnida nor Entognatha contrary to what is suggested in Appendix S2 of InsectChange. The single pair of geographic coordinates used for this study is inadequate as it corresponds to the centre of Lake Oneida, while there were 5 sampling locations in deep waters (plot 135), up to 12 km apart, and 11 sampling locations in shallow waters (plot 136), up to 16 km apart, from the table *Benthos_locations.csv* in the source dataset.

**Study 1430 (Vinson 2001) - Utah freshwater arthropods**

This natural experiment examines the impact of a dam creation and of later partial restoration of thermal conditions, at different sites located upstream and downstream of the dam. InsectChange includes three plots located downstream of the dam, two upstream and one downstream of an intermittent tributary that causes several changes in the physical and biotic nature of the river. But neither the restoration of thermal conditions nor the location of the studied plots in relation to this tributary were mentioned although they were expected to affect the insect dynamics. In Appendix

S2 of InsectChange, van Klink and coauthors stated: "[w]e selected only those sites that had repeated samplings at the exact same location". In contradiction with this statement, plot 1046 includes data from two different sampling locations, 0.8 km and 1 km downstream of the dam, respectively, as reported in the PlotName cell in the table *PlotData.csv*, "0.8-1KDD", and as we checked by comparing the data from the original study and those considered in InsectChange. It would have been possible to exclude the data from the site 1 km downstream of the dam, which covers only the years 1958 and 1959, and to include only the data from the site 0.8 km downstream of the dam, which covers the period 1963-1999. Similarly, plot 1048 includes data from two different sampling locations, 26 km and 27 km downstream of the dam, respectively, as reported in the PlotName cell in the table *PlotData.csv*, "26-27KDD", and as we also checked by comparing the data from the original study and those considered in InsectChange. It would have been possible to exclude the data from the site 27 km downstream of the dam, which covers only the period 1963-1966, and to include only the data from the site 26 km downstream of the dam, which covers the period 1977-1999. In addition, some insect and arachnid taxa were referred as present but not counted in October 1995 in sites 12KDD (Plot 1047) and 26 KDD (Plot 1048), and the counts are thus erroneous at this date. The table *SampleData.csv* refers to "freshwater invertebrates" while only "freshwater insects, arachnids and Entognatha" have been selected. The dataset includes a small proportion of springtails (Collembola, Entognatha), which are neither insects nor arachnids. Finally, there are several samplings in some months but InsectChange does not allow to know their chronology because it does not report the sampling date, which was however available in the original dataset.

**Study 1431 (Slavik et al. 2004) - Alaska freshwater insects**

This controlled experiment includes several sampling sites on a stream, upstream or downstream a site with phosphorus addition, and therefore with phosphorus fertilisation for downstream sites and absence of fertilisation for upstream sites. This long-term stream fertilisation experiment is performed to evaluate the potential eutrophication of an arctic stream ecosystem. The fertilisation status is reported in InsectChange. InsectChange includes two plots, one for the downstream fertilised portion and one for the upstream control portion of the stream, inadequately sharing the same geographic coordinates while there were over 4 km between the most distant sampling sites in the downstream and upstream parts of the stream. The initial study showed that the effects of the fertilisation treatment depended greatly on insect types. The dataset takes the sum of the average densities of *Baetis*, black flies, *Brachycentrus*, chironomids, *Ephemerella*, and *Orthocladius* from Figure 7 of Slavik et al. (2004) but *Orthocladius* insects are actually chironomids and are therefore counted twice. In the fertilised stream, insects increase and their trend is driven by chironomids that spread after moss colonisation following fertilisation.

**Study 1432 (Wallace et al. 2015) - North Carolina freshwater invertebrates**

This study is from a controlled experiment that examines the impacts of litter exclusion, wood removal and addition of artificial wood on benthic invertebrate communities in streams. It is not classified here as a controlled experiment, given that InsectChange only includes data from reference streams where no experiments were carried out. The data includes the biomass of all invertebrates, including Mollusca, Copepoda, Nematoda and Oligochaeta, which are neither insects nor arachnids (see Appendix A in Wallace et al. 2015). It is not possible to separate insects from non-insects in the data and the article does not provide information on the share of insects in the invertebrate assemblage. The two plots actually correspond to two types of substrates (mixed substrate and rockface). The identical geographic coordinates assigned to them in InsectChange are inadequate as the source study specifies that the study samples were collected in three reference streams, 53, 54 and 55, up to 5.6 km apart.

**Study 1433 (Latli et al. 2017) - Belgium freshwater arthropods**

This study is from a natural experiment examining the roles of environmental disturbances and biological alterations on changing abundance and structure of freshwater invertebrate assemblage in different sites of the river Meuse. We coded it as observational since InsectChange only includes one site. The table *SampleData.csv* refers to "Freshwater invertebrates" while only "Freshwater insects and arachnids" have been selected. The geographic coordinates of the unique plot are inadequate: while the table *PlotData.csv* indicates that the source for its location was the publication, the publication only indicated the name of the town (Sassey-sur-Meuse) where the sampling was carried out, but not the precise location on the Meuse River.

**Study 1434 (Blandenier et al. 2014) - Switzerland spiders**

This natural experiment, carried out in a fragmented agricultural landscape of Switzerland, tests the effect of meteorological changes and habitat loss on the abundance of ballooning spider species, depending on whether they are ground-living or upper-strata species. The authors found that ground-living species tended to decrease contrary to upper-strata species and attributed the decrease of ground-living species to their higher sensitivity to weather changes and to a reduction of habitat due to agriculture and urbanisation. In InsectChange, the ballooning spiders are all pooled and assigned to the air stratum, where they were sampled during their dispersal. However,

Appendix 1, Table 1 in Blandenier et al. (2014) allows to assign the species to their respective strata, ground-living, herb-layer, and trees, which would be more meaningful.

**Study 1435 (Johnson et al. 1994) - Kentucky freshwater invertebrates**

This natural experiment compares the dynamics of the macroinvertebrate assemblage of a spring-fed stream in a relatively undisturbed riparian forest in the US among different habitat types, with only two years of data. The study includes three distant study stations (with four plots per station corresponding to the different habitat types), but InsectChange uses the same geographic coordinates for all plots, even though they may be up to 5 km apart. Abundance numbers reported in InsectChange include Turbellaria, Oligochaeta, Isopoda, Amphipoda, Bivalvia and Gastropoda, while Appendix 2 from Johnson et al. (1994) allows the separation of Epheromeptera, Trichoptera and Diptera. Insects generally represent a minor part of the assemblage at the various sites and their share varies over time.

**Study 1437 (Minshall et al. 2001) - Idaho freshwater macroinvertebrates 2**

This natural experiment examines the recovery of the macroinvertebrate assemblage following a wildfire in mixed-conifer forest streams, by comparing five control unburned sites with five burned sites. While the source study contains data on the abundance of some insect taxa, they are not reported in InsectChange. InsectChange only reports biomass data, which cover the whole macroinvertebrate assemblage. The data in the article do not allow the extraction of insect biomass but the article by Richards et al. (1992) mentions the presence of Oligochaeta and Turbellaria in the same streams. InsectChange used an identical pair of geographic coordinates for 10 sites up to 10 km apart (see Figure 1 in the source study of Minshall et al. 2001). This unique pair of geographic coordinates corresponds to the Google maps location of the Frank Church River of No

Return Wilderness while Table 1 of the source publication gave the geographic coordinates of each of the 10 sites. In addition, the pair of geographic coordinates is identical to that of study 1427 covering other streams in the same area. Studies 1427 and 1437 share the same methodology of comparing burned and unburned sites, and present results with identical metrics of density (number/m²) and biomass (mg/m²) of the macroinvertebrate assemblage, in different periods (1979-1989 for study 1437; 1993-2005 for study 1427), but while both metrics were selected in InsectChange for study 1427, only the biomass data was selected for study 1437 (the abundance data was in Figure 5 of the source publication).

**Study 1439 (Cai et al. 2015) - China freshwater chironomids**

This natural experiment studies changes in macrozoobenthic assemblage in a lake subject to severe eutrophication due to high nutrient inputs, in four sections of the lake characterised by different environmental conditions. It includes only two years of data. Deteriorating environmental conditions were accompanied by an increase in chironomid abundance (the only considered insect taxon). The Table *DataSource.csv* inadequately refers to "Mosquitoes (larvae)" in the InvertebrateGroup cell while the original study pertains to total macrozoobenthos and InsectChange only includes data on chironomids. Finally, InsectChange assigned identical geographic coordinates to four zones of the Lake Taihu which sampling sites were up to 60 km apart (see Figure 1 in the source study of Cai et al. 2015). These geographic coordinates are therefore inadequate for a study at the local or landscape level.

**Study 1440 (Hann et al. 2017) - Winnipeg freshwater insects**

This natural experiment evaluates changes in the zoobenthic community abundance and composition in different portions of Lake Winnipeg in parallel with accelerating eutrophication in response to changes in the environmental conditions. In particular, it tests whether the response of the benthic community to eutrophication coincides with the regime change postulated to have occurred in the South basin since the 1990s but not yet in the North basin. However, the environmental conditions in the lake were not considered in InsectChange. InsectChange includes Ephemeroptera, Trichoptera and Chironomidae only, with a large dominance of Chironomidae, which constitute more than 80% of the insects. An increasing trend of insects is observed over time in the three sites, and is highest in the most polluted and eutrophicated site, and this increasing trend is led by chironomids that are known to proliferate in eutrophicated waters. The table *SampleData.csv* inadequately indicates "larval dragonflies" instead of "Ephemeroptera" (mayflies), and "Chronomidae" instead of "Chironomidae", for Figures 5d and 5f of the original study, respectively. Finally, InsectChange includes three sites with different geographic coordinates, but stations sampled in the sites corresponding to each of these plots could be up to approximately 180 km apart (within one site). Therefore, the geographic coordinates are inadequate for a study at the local or landscape scale.

**Study 1441 (Haynes et al. 1999) - Lake Ontario freshwater insects**

This natural experiment examines changes in benthic macroinvertebrate communities in different sites of lake Ontario before and after invasion by *Dreissena* mussels but the factor studied (the invasive species) is not mentioned in InsectChange. The SampleData inadequately refers to "freshwater invertebrates" while only "families Leptoceridae and Polycentropidae of the order Trichoptera and family Chironomidae of the order Diptera" were selected.

**Study 1444 (Groker 2018) - New Zealand freshwater arthropods**

This observational study from the national river quality network in New Zealand investigates the dynamics of freshwater macro-invertebrates in 66 sites throughout New Zealand. The table *SampleData.csv* refers to “all invertebrates” while InsectChange considers only freshwater insects and arachnids (and does not contain freshwater invertebrates belonging to the class Entognatha, contrary to what is suggested in Appendix S2 of InsectChange). Finally, there are two samplings in some months (three occurrences) but InsectChange does not allow to know their chronology because it does not report the sampling date, which was however available in the original dataset.

**Study 1445 (Ernest 2018) - LTER Portal ant nests**

This dataset is from a controlled experiment (LTER Portal) that deals with the dynamics of ant nest abundance in the Chihuahuan desert ecosystem near Portal, Arizona in 22 experimental plots and two control plots. In InsectChange, only the two control plots are considered, thus we did not classify it as a controlled experiment but the two control plots are only poor representative of natural conditions for ant colonies since a set of experimental treatments (such as eliminating ants using poison) were applied to the 22 nearby plots and a surrounding fence protected the 24 plots from livestock. In addition, it is not specified in the table *SampleData.csv* of InsectChange that ant numbers refer to numbers of ant nests. According to its Appendix S1, InsectChange summed up the nest abundances across species per plot per year, after removing some data indicated by data flags: “Because of differences in counting protocol, we removed data with flags 5, 6, 9, and 10. Values in database: Abundance (number of ant nests) per plot per date”. However the authors actually also removed all data for which no data flag was indicated, and therefore only kept date indicated with the other flags present in the source dataset in these plots: flag 1 (missing data stake), flag 2 (suspect data, count missing assumed to be 1), flag 3 (no stake recorded, nearest stake recorded in notes, used for stake value), flag 4 (colony near or on edge of census boundary, distance recorded in notes) and flag 7 (only presence (=1) recorded, not abundance). This results in a major error in insect counts: for three quarters of the colony numbers per plot per year, the authors took into account less than 25% of the actual number of colonies reported in the source dataset and not indicated with data flags 5, 6, 9 and 10. In addition, the methodology described in Appendix S1 of InsectChange is not appropriate to correct for changes in sampling effort. First the authors did not consider flag 2 mentioning that colony counts were missing and assumed to be 1 in 147 censuses. Second, flag 5 stipulates that *Solenopsis xyloni* was not counted in 1978 and 1979, therefore removing the 229 censuses of 1978 and 1979 flagged as 5 does not lead to a consistency of sampling effort, which could have been reached by removing the years 1978 and 1979 or removing the species *Solenopsis xyloni* in all years. The same problem applies to rare species, which were not counted in 1981 for 67 censuses of Plot 11 (flag 6): considering the 1981 sample of Plot 11 is not relevant, even after removing censuses flagged as 6. Fourth, InsectChange did not consider flag 7, which stipulates that only presence, not abundance, was recorded, and which concerns 616 censuses of *Solenopsis* species in the years/periods 1977, 1980-1986, 1988-1994,1998-2009 for which abundance numbers are thus not adequate. It would have been appropriate to remove these two species from the sample. Finally, studies 1353 (abundance of ants) and 1445 (number of nests) are both part of the Portal LTER and both include exactly the same two sites, which are two unmanipulated control plots from the Portal LTER. Therefore, it would have been appropriate to select only one of the two studies.

**Study 1446 (Grechanichenko 2014) - Russia beetles 4**

This observational study investigates the long-term dynamics of ground beetles in different areas of the forest-steppe zone varying in land use and land cover. In InsectChange, one pair of geographic coordinates was assigned to three different plots, described as “oak forest”, “unmowned steppe” and “annually mown steppe” in the PlotName cell of the table *PlotData.csv*. From the source publication by Grechaninenko (2014), sampling was conducted in different biotopes of the area, and given the names of these plots, they cannot share the same pair of geographic coordinates. For this reason, the estimation of the local cropland cover in InsectChange is incorrect.

**Study 1448 (Baranovskaya 1976, Fefilova et al. 2014) - Russia freshwater invertebrates**

This observational study describes long-term changes of aquatic communities in the Russian tundra Bolshoi Kharbei Lake and connecting smaller lakes, subject to indirect anthropogenic stress from regional industrial activities, notably a coal mining complex located at 100 km from the lake. The data included in InsectChange pertain to the same lake as study 1455, the Bolshoi Kharbei Lake (although the table *PlotData.csv* of InsectChange indicates slightly different pairs of geographic coordinates for the two studies, 900 metres apart in the same lake). The abundance series contains only two periods of data, while the biomass series contains three periods. The sampling periods were inadequately reported as 1960, 1990 and 2000, while it was detailed in the original study that sampling dates were 1968 and 1969 in the 1960s, 1998 and 1999 in the 1990s and 2009, 2010 and 2012 in the 2000s. These sampling years were the same as in study 1455 except for 2012, as was the sampling period (not specified here, end of July-early August). Therefore, data in studies 1448 and 1455 overlap and it would have been appropriate to exclude one of these datasets. The abundance and biomass data reported in InsectChange include insects (chironomids and others not detailed), and non-insects (leeches and hydrozoans), while it would have been possible to consider only chironomids. The table *SampleData.csv* inadequately refers to “Macrofauna” instead of “insects, leeches and hydrozoans”. Finally, the geographic coordinates of the unique plot included in the study are inadequate, as sampling sites over the Russian Lake Kharbey were up to 8 km apart (see Figure 1 in the source study by Fefilova, 2014).

**Study 1449 (Baturina et al. 2017) - Russia freshwater invertebrates 2**

This observational study reports the evolution of the biological components of three small reservoirs in a taiga region of Russia. Two reservoirs have data covering a period of at least ten years and are included in InsectChange. For one of them, the data starts three years after the dam was repaired and the bed cleaned. For the other, the last year of observation is after the dam was rebuilt and the level of water dropped and then rose, and is characterised by an outbreak of chironomids. But these two factors (reservoir creation and reservoir reparation) are not mentioned in InsectChange although they highly influence assemblage trends towards an increase. The abundance and biomass data include chironomids and a group of other invertebrates, comprising insects and non-insects (Hydrozoans, Nematoda, Annelida, Tardigrada, crustaceans, Collembola). These groups are inadequately referred to as “Zoobenthos_taxa” in the “GroupInData” cell of the table *SampleData.csv*. The share of chironomids is highly variable and it is not possible to distinguish insects from non-insects in the group of other invertebrates. The biomass numbers reported from the article are incorrect (they represent the sum of the total biomass, the biomass of chironomids and the composite category “others”). Finally, the pair of geographic coordinates used for the reservoirs is not adequate, given that the reservoirs are respectively 2 and 3 km long, with varying surrounding urbanisation, and given that the sampling locations are not indicated in the source study.

**Study 1451 (Aleksevnina and Presnova. 2017) - Russia freshwater invertebrates 4**

This observational study examines changes in benthic populations after the creation of a reservoir, which is a factor largely promoting insect colonisation but this major factor is not mentioned in InsectChange. The table *SampleData.csv* of InsectChange inadequately refers to "Zoobenthos" while InsectChange includes the biomass of chironomids and "others" from the table on p 329 in Aleksevnina and Presnova (2017) which made up less than 10% of the whole invertebrate assemblage. Moreover, the "others" category includes at least crustaceans, which are not insects nor arachnids, and *Corophium curvispinum* and *Dikerogammarus haemobaphes* are reported as Ponto-Caspian invasive crustaceans in the original study. The article states that the role of insect larvae in the benthic fauna of the reservoir is very limited, with only chironomid larvae providing diversity and constituting one of the most numerous groups of bottom insects. Therefore, keeping only chironomid data would have been representative of the insect assemblage. This study covering the period 1964-2014 is about the Votkinsk reservoir in Okhansk. This reservoir is one of the two reservoirs studied in Istomina (2017), which includes sampling in Okhansk (study 1452 of InsectChange covering the period 2000-2016 with overlapping data). It would have been appropriate to select only one of these two studies for the Votkinsk reservoir.

**Study 1452 (Istomina 2017) - Russia freshwater invertebrates 5**

This observational study examines the change in invertebrate assemblage after the creation of two reservoirs, but these factors largely promoting insect colonisation are not mentioned in InsectChange. InsectChange includes eight plots. Plot 455 corresponds to Kama reservoir, while Plots 456, 457 and 458 correspond to the upper, central and dam parts of Kama reservoir, respectively. Plot 459 corresponds to Votkinsk reservoir, while Plots 460, 461 and 462 correspond to the upper, central and dam parts of Votkinsk reservoir; respectively. The assemblage considered is the whole invertebrate assemblage and is dominated by non-insects in plots 455 and 459, while it only includes insects and crustaceans at Plots 456, 457 and 458, and insects and leeches at Plots 460, 461 and 462. The GroupInData cell of the table *SampleData.csv* in InsectChange refers to "groups Zoobentos" at the study level, and users have no information that the assemblages differ among plots. The table *SampleData.csv* does not mention in the "DataCarrier" cell that Table 1 from the original study by Istomina (2017) is used. This is where the data for Plots 455 and 459 were extracted from. These two plots are not reported in the table *PlotData.csv* in InsectChange. They include mostly noninsects, and their total biomass has increased in the reservoirs due to the massive development of large molluscs, which can be calculated as constituting 95% and 96% of the assemblages respectively at the end of the observation period. In addition, each of these two plots is actually an average of three other sites included in InsectChange. These problems were actually introduced in the erratum by van Klink et al. (2020b), which added Plots 455 and 459 in the database that was to become InsectChange, while these plots were absent from the former data supporting van Klink et al. (2020a). Moreover, the six plots included in the table *PlotData.csv* were assigned inadequate geographic coordinates, as each plot aggregates data of several sampling sites, up to 40 km apart within one plot. Besides, data on the dynamics of invertebrate biomass in the Votkinsk reservoir since its construction is also the subject of study 1451 covering the period 1964-2014 and focusing on sampling in Okhansk. This overlaps with this study covering the period 1960-2013 for Plot 459 and the period 2000-2016 (with three periods, 2000-2003, 2007-2009 and 2010-2016, respectively reported as years 2002, 2008 and 2013 in InsectChange) for Plot 461 (which includes sampling in Okhansk). It would have been appropriate to select only one of these two studies for the Votkinsk reservoir.

**Study 1453 (Pavlovsky 2014) - Russia freshwater chironomids**

This observational study investigates the dynamics of the biomass and abundance of macrozoobenthos of Lake Syamozero from 1954 to 1993 in the autumn period. The lake is subject to eutrophication due to an increase in the intensity of the anthropogenic impact but this factor of disturbance is not mentioned in InsectChange. InsectChange deals with chironomidae only while the table *SampleData.csv* inadequately refers to "zoobenthos". Chironomids are known to increase with water eutrophication. Furthermore, the geographic coordinates of the unique plot are inadequate, as the source publication (Pavlovsky, 2014) indicates that samples were collected at several stations in the Russian lake Syamozero, which is approximately 20 km long.

**Study 1454 (Petukhov et al. 2017) - Russia freshwater invertebrates 7**

This observational study analyses the dynamics of the zoobenthic biomass of subarctic Lake Krivoe (western coast of the White Sea) in relation with climatic variables. As described in the article, insects (chironomids) constitute a minor part of the total assemblage, nematodes making the vast majority of invertebrates. The Figure in the original study by Petukhov et al. (2017) separately reports the evolution of zoobenthos in the coastal and deepwater areas of the lake during time periods 2003-2015 and 2004-2015, respectively. In contrast to other datasets (for example study 1429 distinguishing deep and shallow waters in two different plots), InsectChange includes a single site aggregating this data. The biomass counts reported in InsectChange are not adequate because they are calculated as sums (instead of averages) of zoobenthic biomass in the two areas of the lake. As a result, the difference in sampling periods between the two areas is not taken into account: for the first year of observation in 2003, when data are not available for deepwater areas, the biomass figure only includes the coastal areas. The lower, uncorrected sampling effort in the first year of the series underestimates the intercept and biases the slope of the zoobenthic biomass over time for this study. The geographic coordinates indicated for the unique plot in this study, (62.906209, 34.365442), are inadequate for a study at the local or landscape scale as they are approximately 350 km away from the study site, Lake Krivoye near the white sea, with correct geographic coordinates (66.156040, 33.156739).

**Study 1455 (Baranovskaya 1976, Baturina et al. 2012) - Russia freshwater invertebrates 8**

This observational study describes the dynamics of the zoobenthos in the Russian tundra in Lake Kharbei subject to indirect anthropogenic stress from regional industrial activities, notably a coal mining complex located at 100 km from the lake, with a decreasing activity. Data include abundance and biomass of all macroinvertebrates, including Hydrozoa, Turbellaria, Hirudinea, Nematoda, Oligochaeta, Tardigrada, Mollusca, Cladocera, Harpacticoida, Cyclopoida, Ostracoda, Amphipoda and Collembola (Entognatha), whereas it would have been possible to consider the abundance and biomass of insects only, and insects represent a highly variable portion of the total assemblage. As detailed in the comment on study 1448, the studies 1448 and 1455 are on the same lake, with overlapping data, and it would have been appropriate to exclude one of these datasets. Furthermore, this 1455 study includes only one plot. Its geographic coordinates are inadequate as sampling sites over the Russian Lake Kharbey were up to 8 km apart (see Figure 1 in the source study by Baturina, 2012).

**Study 1456 (Nechvalenko 1973, Kurina et al. 2016) - Russia freshwater invertebrates 9**

This observational study examines changes in benthic populations after the creation of a reservoir, a factor largely promoting insect colonisation but not mentioned in InsectChange. Plots 466, 467 and 468 represent the upper, middle and lower parts of the reservoir, respectively. But from Google Earth, in accordance with the geographic coordinates provided in InsectChange, all three are inadequately located in the lower part of the Saratov reservoir, which extends from the city of Syzran' to the Balakovo hydroelectric power plant. Plot 469 (Saratov Reservoir) is inadequately included in the dataset since it represents the average of plots 466, 467 and 468. While the dataset of InsectChange covers all invertebrates, including oligochaetes, polychaetes, crustaceans and molluscs for plots 466, 467, 468, it only includes a subset (representing less than 20% of the whole) with insects, mites, leeches and nematodes for plot 469. The GroupInData cell of the table *SampleData.csv* in InsectChange refers to "Zoobentos" at the study level, and users have no information that the assemblages differ between plots. The abundance and biomass of the whole assemblage are considered for plots 466, 467 and 468, while only biomass is considered for plot 469. Their trend is determined by invasive species such as an invasive polychaete, which broke through in 1988, accompanied by crustaceans, and an invasive snail, which broke through in 2011.

**Study 1457 (Golovatyuk and Abrosimova 2015) - Russia freshwater invertebrates 10**

This natural experiment examines the dynamics of macrozoobenthos composition and abundance from 1991 to 2007 in different sections of a Russian river depending on the hydrological and hydrochemical characteristics of each river section, including a wastewater discharge area. InsectChange includes three sections of the river differing in their amount of pollution (upstream, midstream and downstream of the wastewater discharge area) without mentioning the pollution. It considers a portion of the freshwater invertebrate assemblages composed of insects, arachnids as well as hydrozoans and nematodes, both of which are neither insects, nor arachnids. The "GroupInData" cell of the table *SampleData.csv* inadequately indicates "zoobenthos" instead of these freshwater taxa. The three plots were assigned inadequate geographic coordinates, as each plot aggregates data of several sampling sites, up to 40 km apart within one plot.

**Study 1458 (Kuznetsova 2005) - Russia springtails 2**

This natural experiment compares the 1981-1995 dynamics of springtails (Collembola, Entognatha), which are neither insects nor arachnids, among three sites of the Russian Darwin Nature Reserve differing in pine forests presenting a moisture gradient, i.e. xerophytic (with lichen), mesophytic (with blueberries) and hygrophytic (with sphagnum) pine forests. The moisture factor is not mentioned in InsectChange. The dynamics of springtail populations are shown to depend on biotopes. Springtails are more abundant in the blueberry pine-forest site where they strongly increase, they are intermediate but decrease in the lichen pine-forest site and are less populous but remain stable in the sphagnum pine-forest site. Furthermore, the same geographic coordinates were inadequately assigned to the three plots in InsectChange, while they correspond to three types of forests in the Darwin Nature Reserve, which covers an area of over 1000 $km^2$, and therefore cannot share a unique location.

**Study 1459 (Gryuntal 2008) - Russia beetles 5**

This observational study examines the dynamics of ground beetles in different biotopes in Russia. InsectChange includes two plots inadequately sharing the same geographic coordinates while from the names of the plots, “oak-spruce forest” and “birch forest”, they cannot be in the same location. The most recent data from this study is from 1990, while ESA-CCI data on local cropland cover was available only for plots where sampling ended in 1992 or later, therefore InsectChange provides no local-scale estimate of the cropland cover and urbanisation. It still would be necessary to check whether the unique pair of geographic coordinates provided for these plots is adequate for a study at the landscape scale, for which estimates of cropland cover and urbanisation are provided.

**Study 1460 (Guseva 2017) - Russia rovebeetles**

This interrupted time series controlled experiment examines the effect of different crops on the abundance and composition of rove beetle populations (Staphylinidae), with a particular interest for those of the genus *Aleochara* that are biological control agents for the cabbage root fly. Contrary to what is stated in the table *PlotData.csv*, the geographic coordinates of the study are not given in the source publication by Guseva (2017). The geographic coordinates used in InsectChange are inadequate as they refer to a location in a forest area, while the publication describes that the study was carried out in the agrolandscape of a Russian experimental station. For this reason, the local cropland cover provided in InsectChange is underestimated. The study includes beetle records on sunflower in 1983, rapeseed in 1984 and potato in 2003 and 2005, all planted alternately in the same field, which constitute the time series included in InsectChange, as well as beetle records on forest edges in 2003 and 2004 and a vineyard in 2004, which are not included in InsectChange. The study highlights that the abundance of *Aleochara* beetles is especially high on rapeseed (and this is the reason of the higher abundance of rove beetles in the corresponding year of the time series). However, the different crops were the sampling occurred in the different years of the time series are not specified in InsectChange.

**Study 1461 (Mutin 2015) - Russia hoverflies**

The geographic coordinates reported in InsectChange for the unique plot of this study are inappropriate, as InsectChange uses geographic coordinates obtained from Google maps, which inadequately point to the town centre of Komsomolsk-on-Amur in Russia, 3 km apart from the closest location of the Silinsky park where the sampling occurred.

**Study 1462 (Sasova 2008) - Russia butterflies**

The geographic coordinates reported in InsectChange for the unique plot of this study are inappropriate. Indeed, InsectChange uses geographic coordinates obtained from Google maps, which point to a location in the Ussurisky Nature Reserve in Russia where the sampling occurred. However, the precise locations of sampling within this 400 $km^2$ natural reserve are not specified in the original publication.

**Study 1464 (Shlyakhtenok 2007a) - Chernobyl Sphecidae**

This study examines the dynamics of insect communities in the exclusion zone of the Chernobyl power plant after the Chernobyl disaster in 1986. It states that insect state and dynamics are determined primarily by the level of radioactive contamination and succession. It is hard to see how this time series could be representative of insect trends outside of this very specific area. The drivers of the Chernobyl power plant explosion and of the absence of further anthropic disturbance are not mentioned in the analysis. In addition, the geographic coordinates reported in InsectChange for the unique plot of this study are inappropriate, as sampling was performed in three different villages up to 10 km apart.

**Study 1465 (Chen et al. 2011) - Malaysia moths**

This natural experiment aims at analysing elevation shifts of different insect species in response to climate change, by examining their dynamics in ten sites having different elevations. There are only two years of data. While most plots of study 1465 in InsectChange have approximately the same elevation as in the source study (Figure 1 in Chen et al. 2011), plot 181 has an elevation of 1550 m according to Google Earth, compared to 1450 m in the source study, and its location in InsectChange with respect to the next plots 180 and 182 does not correspond accurately to the map in Figure 1. Therefore, the geographic coordinates of this plot are inadequate. In addition, the local cropland cover is overestimated for two plots given ESA-CCI information and satellite images.

**Study 1466 (Krupa et al. 2013) - Kazachstan freshwater invertebrates**

This observational study analyses the dynamics of aquatic communities in the Balkhash Lake located in an arid zone of Kazakhstan, as a function of water level, water salinity and mean annual air temperature. The data include all freshwater invertebrates, and it is not possible to distinguish insects. According to the source study (Krupa et al. 2013), "*Since 1996, there was a tendency towards a sharp increase in the bottom community biomass ... due to the mollusc* Hypanis colorata *...The share of* Hypanis colorata *in the total macrozoobenthic biomass has increased over the recent decades to 75.3-95.6 %*" (own translation from Russian). *Hypanis* [later revised as *Monodacna*] *colorata* is an invasive clam species of Caspian origin, introduced secondarily into Lake Balkhash (Vinarski and Kantor 2016). But the disturbance due to this invasive species is not mentioned in InsectChange. The geographic coordinates of the unique plot of this study are inadequate for a study at the local or landscape level, as the source study by Krupa (2013) indicates that samples were collected in 60 stations covering the entire area of Lake Balkhash (Russia), which is over 400 km long.

**Study 1467 (Shlyakhtenok 2007b) - Belarus Aculeate Hymenoptera**

This observational study examines the dynamics of Hymenoptera Aculeata in raised bogs in Belarus over the period 1986-2003. It shows that abundance of wasps from various families varied noticeably from year and year and was correlated with climatic parameters (notably air temperature and sum of precipitation in spring). Abundance numbers were erroneously retrieved from Figure 2 of the source study by Shlyakhtenok (2007b): the highest abundance count is in July 1992 in the source study, as opposed to July 1993 in InsectChange. In addition, the geographic coordinates reported in InsectChange for the unique plot of this study are likely inappropriate as the precise sampling area is not specified in the source study and there is no source for the geographic coordinates used in InsectChange. Finally, the local cropland cover provided in InsectChange is overestimated given information from the source study.

**Study 1468 (Ploquin et al. 2013) - Spain bumblebees**

This study is from a natural experiment, which samples bumblebees in several sites in the Asturian region of Spain in order to analyse changes in the abundance and distribution of bumblebee species along an elevational gradient. The abundances showed different trends according to species and altitude. InsectChange only includes two record periods (1988-1989 and 2007-2009), reported as 1988 and 2008 in InsectChange, and one plot, aggregating 32 sites from different altitudes that were sampled in both periods. The geographic coordinates of this plot are inadequate for a study at the local or landscape scale level, as sampling sites were up to almost 200 km apart from the source publication by Ploquin (2013). For this reason, the local cropland cover provided in InsectChange is incorrect. Since abundance data were available only for this average plot in the original study, we did not classify it as a natural experiment.

**Study 1470 (Shlyakhtenok 2007c) - Belarus Chrysididae**

This observational study examines the dynamics of Chrysididae in various ecosystems in Belarus and their relation with different climatic factors with a correlative approach. InsectChange includes a single plot averaging the different sites. The source study by Shlyakhtenok (2007c) indicates that the study covered several natural areas in Belarus, including the Berezinsky Biosphere Reserve and the Polesie State Radioecological Reserve that are 370 km distant. The unique pair of geographic coordinates given in InsectChange is not provided in the original publication and is outside of the global sampling area.

**Study 1471 (Nitochko 2012) - Ukraine beetles**

This observational study examines the dynamics of ground beetles and tenebrionid beetles in two sites of the Black Sea Biosphere Reserve of Ukraine, which area is approximately 900 $km^2$. InsectChange includes data from one site, located in the Ivano-Rybalchansky district of the reserve. No source is indicated for the geographic coordinates provided for this plot in InsectChange and the InsectChange-assigned location is 27 km distant from the geographic coordinates for this site provided in Pilato et al. (2011).

**Study 1472 (Szabó et al. 2007) - Hungary moths 2**

This observational study explores the dynamics of night-active Macrolepidoptera fauna in the Aggtelek karst region in Hungary from 1990 to 2004. The geographic coordinates provided in the publication and incorporated in InsectChange are approximate and located 48 km from the light trap. The local cropland cover provided in InsectChange is overestimated given information from the source study and satellite images at the correct location.

**Study 1473 (Mebane et al. 2015b, 2015a) - Idaho freshwater invertebrates 3**

This natural experiment investigates the recovery of freshwater invertebrate communities in mining-damaged streams following efforts to restore water quality, comparing reference streams and restored mining-damaged streams. The observation period is 1993-2013, restoration efforts began in 1995, and a wildfire affected all streams in 2000. As specified by Mebane et al. (2015a), datasets were collected with different sampling and analysis methods prior to 2002. This prevents comparisons of absolute values of abundance and biomass before and after 2002. In Mebane et al. (2015a), instead, samples from mining-influenced sites were normalised to values collected from concurrent reference sites enabling to extend record across the older datasets and to compare the relative values. By including time series on absolute values of invertebrate abundances from 1993 to 2013, InsectChange does not take this change in sampling method into account. Moreover, InsectChange includes abundance and biomass numbers for the whole invertebrate assemblage, whereas data on the abundance of insects could have been extracted from Appendix 2 of Mebane et al. (2015a). For the biomass data, from numbers in the table *InsectAbundanceBiomassData.csv*, plot 285 where the invertebrate trend is increasing is a pooling of plots 279, 281 and 282. This plot is not listed in the table *PlotData.csv*.

**Study 1474 (Schuch 2011, Schuch et al. 2012a) – Germany Orthoptera, Hemiptera: Heteroptera & Auchenorrhyncha**

This natural experiment compares the abundance of Auchenorrhyncha, Heteroptera and Orthoptera in Lower Saxony (Germany) between 1951 and 2009 for nine sites (mainly pastures) differing in moisture and agricultural practice (some mowed or grazed, some not in the last year) according to the source study by Schuch (2011), but InsectChange does not mention these factors susceptible to influence insect trends. Appendix S1 of InsectChange states that the taxonomic focus is "Orthoptera, Hemiptera: Heteroptera, Hemiptera: Auchenorrhyncha", inadequately referred as "plant and leafhoppers" in the table *SampleData.csv*. There is a major inconsistency in the sampling effort between the periods reported in InsectChange. For example, for Plot 489 (site 1 in the original study), the table *InsectAbundanceBiomassData.csv* reports three abundance counts in August 1951 (4, 258 and 366). We checked in the original dataset that Orthoptera, Heteroptera and Auchenorrhyncha were all sampled on August 6 and 21, 1951, with respective total abundance counts of 258 and 366, while actually only Orthoptera were sampled on August 30, 1951, with an abundance count of 4. This problem of change in sampling effort only occurs in the first year of record, 1951 and not in the unique other year of record, 2009, thereby biasing the insect trend of the dataset by reducing its rate of decrease. Moreover, there were several samplings in most months (e.g. three samples in August 1951 for Plot 489 in the example given above) but InsectChange does not allow to know their chronology because it does not report the sampling date. Finally, the local cropland cover provided in InsectChange is inconsistent with information from the source study and satellite images, with an overestimation of cropland cover for all plots.

**Study 1475 (Schuch 2011, Schuch et al. 2012b) – Eastern Germany Hemiptera: Auchenorrhyncha**

This observational study examines the dynamics of Auchenorrhyncha assemblages in dry grasslands of eastern Germany in 27 sites equally distributed in Brandenburg, Thuringia and Saxony and reveals a general decrease mostly explained by modern land use practices and habitat loss. There are two samples in July 2009 for plot 502 but InsectChange does not allow to know their chronology because it does not report the sampling date. Finally, the local cropland cover provided in InsectChange is inconsistent with information from the source study and satellite images for several plots, with an overestimation.

**Study 1476 (Swengel and Swengel 2015a) - US butterflies Grasslands**

Studies 1476 (US butterflies Grasslands), 1477 (US butterflies Barrens) and 1478 (US butterflies Bogs), all three on butterflies in Wisconsin (US), were carried out with similar methods by the same investigators and are presented together in Appendix S1 of InsectChange. For consistency with other studies in the dataset, it would have been more relevant to group them together. In these three studies, based on raw data from the authors, InsectChange includes 138 different sites, some of them with identical geographic coordinates, and some others only 13-metre distant, while publications in relation to this dataset considered only 58 sites (Swengel, A. B., and S. R. Swengel. 2015a and Swengel, S. R., and A. B. Swengel. 2015b). Inflating the number of sites in these studies leads to overweighting them compared to other studies. Moreover, there are sometimes several samplings per month (e.g. three samples in July 2017 for plot 339) but InsectChange does not allow to know their chronology because it does not report the sampling date. In addition, the local cropland cover provided in InsectChange is inconsistent with information from the source study and satellite images.

**Study 1477 (Swengel and Swengel 2015b, 2015a) - US butterflies Barrens**

See remarks on study 1476. In addition, 5 plots (350, 351, 359, 360, 368) include six abundance data in some months (respectively in 2, 1, 2, 2 and 2 months), and five or less in all other months. This is not consistent with the methodology detailed in InsectChange (p. 17 of the detailed presentation), describing that the temporal resolution was between the week and the year, except in six datasets where plots were sampled between six and eight times in any month. Finally, there are several samplings per month for about half of the months but InsectChange does not allow to know their chronology because it does not report the sampling date. In addition, the local cropland cover provided in InsectChange is inconsistent with information from the source study and satellite images, with an overestimation or an underestimation depending on plots.

**Study 1478 (Swengel and Swengel 2015b) - US butterflies Bogs**

See remarks on study 1476. In addition, there are several samplings per month for about half of the months (e.g. three samples in June 2013 for plot 393) but InsectChange does not allow to know their chronology because it does not report the sampling date.

**Study 1479 (Driessen and Kirkpatrick 2017) - Tasmania arthropods**

This natural experiment compares the dynamics of invertebrates from different strata (soil surface and herb layer) on sites that were subject to more or less recent fires and different burning regimes. The history of each site with respect to burning is not reported in InsectChange (except for mentioning which sites were or not subject to recent burning). Appendix S1 of InsectChange states that the taxonomic focus is "all arthropods. Myriapods and crustaceans are excluded for this data product". From Table 5 in the source publication by Driessen and Kirkpatrick (2017), this assemblage includes springtails (Collembola, Entognatha), which are neither insects nor arachnids, and which were the main constituent of the ground-active invertebrates and a significant constituent of the foliage-active invertebrates in 2004. In addition, the table *SampleData.csv* refers to "terrestrial invertebrates", while the adequate taxonomic focus is insects, arachnids and springtails.

**Study 1480 (Doran et al. 2003) - Tasmania invertebrates**

This natural experiment examines the relationships between the altitudinal gradient and the associated transition in biodiversity. The dataset spans over four years but actually covers four months, December, January, February and March, in 2001-2002 then in 2011-2012, therefore including eight time-records in only two years. From Table 1 in the source publication by Doran et al. (2003), there were non-insects and non-arachnids among the major taxa of the invertebrate assemblage: Mollusca, Annelida, Enthognata (Diplura), Crustacea, Chilipoda, Diplopoda. From the table *SampleData.csv*, the assemblage included in InsectChange is composed of “terrestrial invertebrates” (GroupInData cell in the table *PlotData.csv*). Appendix S1 of InsectChange inconsistently reports that the taxonomic focus was ground-dwelling arthropods, excluding myriapods and crustaceans. In both cases, whether the assemblage is the one described in the table *SampleData.csv* or in Appendix S1 of InsectChange, it includes non-insects and non-arachnids.

**Study 1481 (Pe’er and Comay 2019) - Israel butterflies**

This observational study explores the dynamics of lepidopteran abundances in several localities of Israël from 2010 to 2018. There are several samplings in most months but InsectChange does not allow to know their chronology because it does not report the sampling dates. The geographic coordinates are inappropriate for Plot 1691 because sampling occurred on a transect where sampling points were up to 1.5 km distant. In addition, the local cropland cover provided in InsectChange is inconsistent with satellite images, with an overestimation for several plots.

**Study 1484 (Lister and Garcia 2018) - Mexico arthropods**

This observational study compares the mean dry weight biomass of arthropods caught per day with sticky ground traps in the Chamela forest in Jalisco, Mexico, in July 1987 and 1988 and again in August 2014. InsectChange includes these data in plot 929 but insect biomass numbers were inaccurately reported from Fig S2 of the supplementary materials by Lister and Garcia (2018) as respectively 2452, 1874 and 271, in 1987, 1988 and 2014, while the correct figures in Fig S2 were 491, 375 and 51. This error biases the trend by slightly minimising the insect decrease. In addition, for ID 275 in the table *SampleData.csv*, there is an error in the stratum assignment: the correct stratum is soil surface, and not herb layer, for these arthropods caught in ground traps. The assemblage is inadequately referenced as “all arthropods” in the table *SampleData.csv*, while it consists of ground-dwelling arthropods. Finally, the geographic coordinates provided in InsectChange are inappropriate because they point to a location in the biological station in Mexico where the sampling occurred but not necessarily to the study site in this 16 $km^2$-wide biological station.

**Study 1485 (Lister and Garcia 2018) - Luquillo arthropods**

This observational study examines the dynamics of the average dry-weight biomass of arthropods caught per 12-h day in ground and canopy traps within the same sampling area in the Puerto Rico's Luquillo rainforest over two periods and four dates, July 1976, January 1977, July 2012 and January 2013. The ground and canopy arthropod data were differentiated into two plots, plot 930 and plot 931 in InsectChange, even though they were from the same study area according to the Lister and Garcia publication. They could simply have been assigned to the same plot but differentiated by stratum, as was the case in other datasets in InsectChange. Moreover, plot 931 (canopy), with a strongly decreasing trend in insects, was incorrectly assigned the soil stratum (instead of the tree stratum). Its inclusion in the tree stratum is clear from the material and methods of Lister and Garcia (2018) and from Table 4b of the article. The exclusion of this plot from the tree stratum biases this stratum towards an arthropod increase, especially since this tree stratum has a small sample size. In the table *SampleData.csv*, the arthropods were inadequately referenced as "all arthropods", while it would have been appropriate to refer to herb-layer arthropods for ID 277 and ground and canopy-layer arthropods for ID 276, and to split this 276 SampleID row into two rows, one for each stratum. The invertebrate group included in InsectChange is not consistent across plots and this cannot be easily inferred since InsectChange does not indicate the invertebrate group at the plot level.

**Study 1487 (Schowalter 2017) - LTER Luquillo canopy arthropods**

This study is a natural experiment comparing canopy arthropod dynamics on different types of trees from 1991 to 2016 in two types of areas: gaps created by Hurricane Hugo in 1989 (openings in the canopy, in which all or most of the trees fell or had their main stems broken during Hugo) and non-gaps (plots in which few or no canopy trees fell or were severely damaged by the hurricane). In addition, the plots where arthropods were sampled did not correspond to geographic locations but to individual trees. The arthropod trend after hurricane disturbance is shown to depend on the insect guild in the original studies. InsectChange includes insects, arachnids, as well as springtails (Collembola, Entognatha) which are neither insects nor arachnids (and not "all arthropods" as incorrectly mentioned in the table *SampleData.csv*).

**Study 1488 (SLU 2018) - Sweden freshwater arthropods**

The "GroupInData" cell of the table *SampleData.csv* inadequately refers to "Freshwater invertebrates", suggesting that the entire freshwater assemblage has been selected, whereas only insects and arachnids have been selected in InsectChange. In addition, there are several samplings in some months but InsectChange does not allow to know their chronology because it does not report the sampling date, which was however available in the source dataset. Also, some data available in the original dataset are missing in InsectChange, as is the case for 2012 and 2013 abundance counts in Plot 1723, which are reported as "NA" in InsectChange table InsectAbundanceBiomassData.csv whereas they are respectively 27.6 and 275 individuals per sample according to the source data. Finally, some geographic coordinates assigned in InsectChange are inappropriate, as they do not correspond to the sampling sites. For example, InsectChange plots 1723 and 1578 are assigned the same geographic coordinates pointing to the middle of the southern part of Lake Bolmen in Sweden (Bolmen, södra in the original dataset), whereas the source data distinguishes two sampling sites, one near the south-eastern shore of the lake and the other further south in the middle of the lake, each three kilometres distant from the InsectChange location and 2 kilometres from each other.

**Study 1491 (Sistema de Informação sobre a Biodiversidade Brasileira - SiBBr 2018, Aguila et al. 2018) - Brazil Freshwater arthropods 2**

This observational study investigates the dynamics of freshwater macroinvertebrate community sampled biannually from 1999 to 2010 in lakes from the middle Doce river lake system, and rivers segments in the Piracicaba River basin (sub-basin of Doce river), Minas Gerais State, Brazil. The SampleData refers to "all invertebrates" while from its Appendix S2 InsectChange includes only taxa belonging to the classes Insecta, Arachnida or Entognatha (Entognatha, which are neither insects nor arachnids, in a small proportion in the dataset). Most counts were inadequately reported from the source data to InsectChange. For example, for site 1059 (lake Águas Claras), the mean abundance count for the taxa considered was 181 in Winter 1999 (1 sample) and 23 in Summer 2009 (total of 138 divided by 6 samples), while InsectChange reported abundance counts of 107 and 36.5, respectively, for these two periods. Finally, from the source study, there were two or more sampling stations in the coastal region of each lake to ensure that spatial heterogeneity was considered. The unique pair of geographic coordinates provided in the source study for each lake does not indicate the precise sampling locations.

**Study 1493 (Hu et al. 2016) - UK migrating insects**

This study records biomass and abundance of migrating insects at high elevation in Southern England (> 150 m above ground) during 10 years. The study region is the volume of atmosphere between 150–1200 m above ground level, above an area delimited by a circle of 300 km diameter (Hu et al. 2016, Supplementary materials). This is a much higher elevation than in other studies pertaining to flying insect: suction traps at 12.2 m above ground level in studies 1495 and 1496; Malaise traps at the ground level in study 1409. It is not adequate to consider the impact of the local land cover at the ground level, in a 900 m-side square around a point defined by is unique pair of geographic coordinates inside this 300 km-diameter circle, on these migrating insects. For this reason, the estimation of the local cropland cover in InsectChange is incorrect. The data sometimes greatly vary over time, depending on the dominant wind during the year. But while variation in the number of insects observed at such high flight altitude in the study region may reflect, for a large part, variation in wind direction, rather than variation in population size, this wind factor was not taken into account in InsectChange.

**Study 1494 (Aebischer 1990) - UK sawflies**

This study is from an interrupted time series natural experiment examining the impact of the introduction of the dimethoate aphicide on the dynamics of sawfly abundances in five farms cultivating cereals. We classified it as an observational study since InsectChange only includes the period from 1970 to 1988 before the application of dimethoate, and not year 1989 of dimethoate application. A single pair of geographic coordinates was inappropriately used for the five different farms (each corresponding to a plot in InsectChange), although the original study specified that the data stemmed from a monitoring study of cereal invertebrates on a 62-km2 area of farmland in West Sussex.

**Study 1495 (Shortall et al. 2009) - UK flying insects**

This observational study examines changes over 30 years in the total biomass of aerial insects collected in four areas of Southern Britain in 12.2 m high suction traps (Rothamsted Insect Survey). The table *SampleData.csv* inexactly refers to "all flying invertebrates" instead of "flying insects" as specified in the table *DataSource.csv*. See comments for study 502: plots 895, 896, 900 and 902 (respectively, Hereford, Rothamsted Experimental Station, Wye College and Starcross) of study 502 on aphids are actually the same as plots 1617, 1618, 1620 and 1619, respectively, in study 1495 on flying insects (including aphids) in the period 1973-2001. Therefore, some insect data from this study overlap with that of study 502, covering the period 1969-1990. It would have been appropriate to select only one of these datasets.

**Study 1496 (Benton et al. 2002) - UK Stirling arthropods**

This observational study examines the dynamics of arthropods from 1972 to 1998 using a high suction trap operating at the farmland edge of the University of Stirling campus (UK) and investigates the effect of agricultural practices and climate change on their dynamics via a correlative approach. The table *DataSource.csv* of InsectChange inadequately mentioned "Flying insects" instead of insects, arachnids and springtails, springtails (Collembola, Entognatha) being neither insects nor arachnids. The table *SampleData.csv* indicates Diptera and other orders, when it would be better to specify the groups involved, i.e., Diptera, Hymenoptera, Coleoptera, Thysanoptera, Collembola, and Spiders. While the authors summed the abundance numbers of Diptera in Fig. 1e and the abundance numbers of the other orders shown in Fig. 3a,b,c,d,e, they improperly took as biomass data the volume of insects shown in Fig. 3f but did not specify it in the sample data file. The insects there refer to the total number of insects trapped and not just the cumulative numbers of the six groups shown in Fig. 3, so there is an inconsistency between the groups used for the abundance and biomass data. Finally, the estimation of the local cropland cover provided in InsectChange is underestimated given information provided in the source study.

**Study 1497 (Brown and Roy 2018) - UK ladybeetles**

This observational study examines the dynamics of native and invasive ladybeetles in different sites varying for land cover and anthropogenic impact. In the tree stratum, changes were observed in two lime tree sites and one pine tree site. In the lime tree sites, trees were pollarded (in 2013 in plot 1627; in 2008 and 2013 in plot 1628), while in the pine tree site (plot 1629) trees were not pollarded. Each pollarding event was followed by an increase in abundance of the invasive ladybeetle as well as the total ladybeetles, while the ladybeetle population decreased at the non-pollarded pine tree site. These pollarding events strongly affected ladybeetle abundances and more specifically those of the invasive species but these highly disturbing factors are not reported in InsectChange. In this study, insect trends show important variations depending on sites and no clear insect trend emerges. Finally, the local cropland cover provided in InsectChange is overestimated for two plots given information from the source study and satellite images.

**Study 1498 (Woodward et al. 2015) - Ireland freshwater invertebrates**

This observational study examines the recovery of an invertebrate assemblage from a catastrophic flood that occurred in 1986, in the beginning of the observation period (1985-1998) but this catastrophic flood is not mentioned in InsectChange. InsectChange includes the abundance of all freshwater invertebrates, including non-insects such as molluscs, crustaceans or Amphipoda, whereas it would have been possible to include only the abundance of insect taxa (Chironomidae, Ephemeroptera, Plecoptera and Trichoptera). In addition, the abundance numbers for the invertebrate assemblage reported in InsectChange are incorrect due to an inadequate transformation of the numbers reported on a log10 scale in the article. The geographic coordinates assigned to the unique Plot 1631 in InsectChange are inadequate because they are approximately 24 km from the actual sampling site. They were extracted from the source study by Woodward et al. (2015), which was too imprecise, and did not correspond to the location of the sampling site provided in Figure 1 of this publication. The adequate geographic coordinates of the plot are provided in Murphy and Giller (2000).

**Study 1499 (Soulsby et al. 1995) - Scotland mayflies**

This dataset is from a natural experiment, which examines the recovery of mayflies (particularly sensitive to acidity) in different streams affected by post-industrial acidification of water in North-East Scotland, after acid deposition has been reduced. It finds that the abundance of mayflies has increased in acid sensitive streams, but not in chronically acidified streams. In the article, the dynamics of mayfly abundance is shown in an acid sensitive site and in a chronically acidified site, taken as examples. InsectChange only includes the acid sensitive stream characterised by an increase in mayfly abundance, but not the chronically acidified stream where abundance is close to zero in all years. The initial disturbance, i.e. the acidity of the stream is not reported in InsectChange.

**Study 1500 (Durance and Ormerod 2007) - Wales freshwater invertebrates**

This observational study is from a natural experiment which analyses changes in stream macroinvertebrates in the United Kingdom in relation to winter climate, climate change and acidification. InsectChange reports abundance numbers of the less acidic group of streams, the two circumneutral moorland streams being reported as a single plot with mean abundance values over streams, but these abundance data are only available for the whole invertebrate assemblage, including non-insects. The PlotName cell inadequately refers to "River Towy" for the unique plot of this study, while from the source study (Durance and Ormerod, 2007), this plot includes data from two tributary streams of the Tiwy River in the UK, LI6 and LI7. The geographic coordinates for this plot are inadequate, as first, these two streams are 1 km apart, and second, the geographic coordinates assigned to this plot in InsectChange are 1 km apart from the nearest of these two streams. The sampling period is inadequately reported as "19" in the table *InsectAbundanceBiomassData.csv*, while samplings occurred in April according to the source study.

**Study 1501 (National Biodiversity Data Centre 2018) - Ireland butterflies**

This observational study is from the Irish butterfly monitoring scheme, a running citizen science scheme from the Irish Biodiversity Data Centre, which monitors butterfly populations on a weekly basis between 1st April and 30th September of each year based on a network of transects established and walked by volunteer recorders. InsectChange includes only two sites, WX01 - Raven Nature Reserve (Plot 1672) and WX05 - Wildfowl Reserve Drive (Plot 1671). Plots 1671 and 1672 include six abundance data in one month for each of them, and five or less in all other months. This is not consistent with the methodology detailed in InsectChange (p. 17 of the detailed presentation), describing that the temporal resolution was between the week and the year, except in six datasets where plots were sampled between six and eight times in any month. Moreover, there are several samplings per month for most months but InsectChange does not allow to know their chronology because it does not report the sampling date, which was however available in the original dataset. In addition, the geographic coordinates assigned to Plot 1671 in InsectChange are inadequate as they are approximately 900 m distant from those in the source study. Finally, the local cropland cover provided in InsectChange for this plot is underestimated given satellite images.

**Study 1502 (Zhang et al. 2018) - China ground beetles**

This natural experiment compares the recovery of carabid assemblages after desalination and fertilisation processes between sites where desalination occurred at different dates, and began before the beginning of insect sampling. The context of the different sites with respect to this desalination is not reported in InsectChange. Finally, a unique pair of geographic coordinates was inadequately assigned to three plots located in three different villages (Zhang et al. 2018). For this reason, the estimation of the local cropland cover in InsectChange is incorrect.

**Study 1503 (Paul et al. 2018) - Australia freshwater invertebrates**

This observational study investigates the long-term dynamics of the macroinvertebrate assemblage in six different sections of an Australian river in relation to salinity mitigation strategies, floods and droughts. The desalination processes in the beginning of the observation period, which promote invertebrate recovery, are not mentioned in InsectChange. There were also high disturbances such as major droughts and flood events, which are not mentioned either. Abundance numbers reported in InsectChange include all macroinvertebrates, and it is not possible to separate insects from other invertebrates such as crustaceans, mussels, snails, and worms.

**Study 1504 (Herbst et al. 2018) - California mine freshwater invertebrates**

This natural experiment analyses the recovery of freshwater invertebrates from acid mine drainage, comparing unaffected reference streams with streams exposed to metal contamination, where water was actively treated one year after the beginning of observation, and where regular depollution events took place during the observation period. The abundance data reported in the data document involve the entire invertebrate assemblage, including oligochaetes, turbellarians and ostracods. It was not possible to distinguish insects in the data.

**Study 1505 (Blanchet et al. 2018b, 2018a) - Finland oak herbivores**

This study is from a natural experiment examining whether feeding guild, voltinism, similarity in parasitoid community and/or phylogenetic relatedness explain similarities in temporal dynamics among herbivorous community members from Finland and Japan. It is classified here as an observational study, given that InsectChange only includes the site in Finland (11 years), characterised by a strong increase in insect populations. The site in Japan (9 years), with a decreasing trend in insect populations, is not included in InsectChange. This is not consistent with the inclusion of plots with nine years of data if other plots within the same data set had at least 10 years (Appendix S2 of InsectChange). The exclusion of the Japan site biases the insect trend towards an increase in this study. In addition, the geographic coordinates provided in InsectChange are inappropriate as they were retrieved from Google maps and point to a terrestrial location in the 5 $km^2$ island where the sampling occurred, but not necessarily the actual sampling site. Satellite images show that the local cropland cover varies on the island, therefore the local cropland cover at InsectChange geographic coordinates is not necessarily representative of the one at the actual sampling site.

**Study 1506 (Gutiérrez-Fonseca et al. 2018) - Costa Rica freshwater invertebrates**

This controlled experiment includes two streams observed from 1997 to 2011 that flow into the Rio Sarapiquo (Costa Rica), one with an experimental phosphorus enrichment over eight years (from 1998 to 2006), and the other without phosphorus enrichment, but these experimental conditions are not mentioned in InsectChange. The study also examines the effect of climate change and increased flood events on the abundance and biomass of the entire macroinvertebrate assemblage. It was not possible to distinguish insects from non-insects but the article specifies that contributions from non-insects to the total abundance were modest, with approximately respectively 3.1% and 3.6% of the total abundance, and 7.2% and 9.6% of the total biomass, in each of the two sites. The same geographic coordinates were inadequately assigned to both plots, corresponding to the two streams. The source study by Gutiérrez-Fonseca et al. (2018) does not provide information on their locations and sampling sites.

**Study 1507 (Shulepina 2010) - Russia freshwater invertebrates 11**

This pollution-disturbed study is from a natural experiment, which examines the dynamics of the structure of macrozoobenthos and tests its value as an indicator of water quality in various types of water bodies in the Yenisei River basin under different anthropogenic pollutions, in particular pharmaceutical and aluminium production. InsectChange includes the dynamics of the entire macrozoobenthic biomass and abundance of the Krasnoyarsk reservoir, one of these water bodies exposed to a complex impact of industrial and residential factors, but the pollution is not mentioned. As only this site is included in InsectChange, because the source study did not report assemblage changes over time for the other sites, we did not classify this dataset as a natural experiment. It is not possible to separate insects from non-insects in the assemblage. However, the details reported in the article show that insects were mainly chironomid larvae and that they represented only 37% of the total biomass in 1981, and 11% in 2002; a peak in total biomass was observed at the end of observation time (2005) and was due to the massive development of invasive amphipods, thus biasing the trend of the study towards an increase. The geographic coordinates of the unique plot are inadequate for studies at the local or landscape scales, given that surface area of the Krasnoyarsk Reservoir is 2,000 km². The sampling areas are not specified in the source study by Shulepina (2010).

**Study 1508 (Chernenkova et al. 1995, Tanasevitch et al. 2009) - Russia soil fauna 2**

This natural experiment covers all mesofauna (including earthworms, myriapods, molluscs, crustaceans, chilipods) and it is not possible to separate insects from non-insects. These invertebrates are referred to in the table *DataSource.csv* as freshwater invertebrates whereas they are underground terrestrial invertebrates. The study compares sites at a greater or lesser distance from a polluting plant emitting sulphur, nitrogen, fluorine and oxides of heavy metals into the atmosphere, and having decreased its emissions at the beginning of the study and then four times more during the observation period. These natural experimental conditions are not reported in InsectChange. In the four sites included in the study, between the two years included in the study, abundance numbers increase respectively by 0%, 7%, 26% and 71%, while biomass numbers decrease by 35% and 10% on two sites and increase by 1% and 7% on two other sites. From Google Earth, the locations given for the four plots in InsectChange are each about 3 km further away from the Severonikel factory than what was described in the article and specified in the PlotName cell of the table *PlotData.csv* in InsectChange, and are therefore inadequate. There are only two years of data.

**Study 1509 (Kashulin et al. 2012) - Russia freshwater invertebrates 13**

This observational study examines changes in the structure and functioning of hydrobiont populations under conditions of high pollution in two Russian lakes subject to eutrophication, due to water pollution by heavy metals in association with climate warming, but the highly disturbing pollution context is not mentioned. The original study reports the dynamics of the abundance and biomass of the whole invertebrate community and separate data for insects are only available for one lake in one year and show that insects represent a minor part of the abundance and biomass of-the invertebrate assemblage, which are dominated by Oligochaetes. In lake Imandra, the freshwater invertebrates are dominated by Oligochaetes and chironomids but the amphipods increase 18-fold from 1996 to the end of the observation, biasing the trend of the dataset towards an increase. The geographic coordinates are inadequate for a study at local scale as Plot 1712 represents lake Ozero Malyy Vud"Yavr in Russia, which is 1.6 km long Plot 1713 represents lake Imandra in Russia, which covers an area of over 800 km².

**Study 1510 (Novoselov et al. 2017) - Russia freshwater arthropods 14**

This observational study examines the dynamics of abundance and biomass of freshwater macro-invertebrates in lake Lacha in Russia from 2003 to 2015. The “GroupInData” cell of the table *SampleData.csv* inadequately refers to “freshwater invertebrates”, while only freshwater insects and arachnids were selected in InsectChange. The surface area of Lake Lacha is 356 km² and the source study by Novoselov (2017) specified that sampling was performed in 12 stations across the lake, approximately up to 15 km apart, as represented in Figure 1 of this publication. Therefore, the geographic coordinates of the unique plot considered on this lake are inadequate for a study at local scale.

**Study 1511 (Bezmaternykh et al. 2008) - Russia freshwater invertebrates 15**

This study is from a natural experiment that analyses the zoobenthos in different parts of the Siberian lake Chany in relation to substrates, water level, salinity and depth. Data consider biomass of the whole invertebrate assemblage including gastropods, bivalves, crustaceans and worms (Oligochaeta). For the last year of observation (2004), an upper estimation of the insect share

(64.3%) could be processed from the means of Dipterans + 'others' biomass and the overall biomass available from different locations of the lake in figure 1 in Bezmaternykh et al. 2008. The share of insects in the assemblage is highly variable in the different locations of the lake. InsectChange includes a single site for this study while there were 53 sampling points with different geographic coordinates across this lake of 2,000 km², up to 64 km apart, therefore its geographic coordinates are inadequate for a study at the local or landscape scale. InsectChange reports the dynamics of the average biomass of the whole invertebrate community in the lake from 1925 to 2004, including non-insects and non-arachnids. As only average biomass data was available in the original study, we did not classify it as a natural experiment.

**Study 1512 (Homburg et al. 2019) - Germany ground beetles**

The geographic coordinates reported in InsectChange for the unique plot of this study are very likely inappropriate, as the original publication does not mention the precise location of the study site within the 231.5 km² Lüneburg Heath Nature Reserve in Germany, and there is no indication in InsectChange on how the geographic coordinates pointing to a location in this nature reserve were chosen from Google maps.

**Study 1513 (Lencioni 2018) - Italy freshwater invertebrates**

This natural experiment studies the influence of glacier recession on macroinvertebrates in streams, including glacier and non-glacier sites, but these conditions are not mentioned in InsectChange. The abundance data reported in InsectChange includes the entire invertebrate assemblage. It would have been actually possible to distinguish the abundance of insects at two of the six sites. Each site has data for only two years.

**Study 1515 (Cuesta and Lobo 2019) - Spain dung beetles**

Only two years with data

**Study 1516 (Gran and Götmark 2019) - Sweden saproxylic beetles**

This controlled experiment examines the dynamics of saproxylic and herbivorous beetles during three years of record, 2001, 2004 and 2013, on seven sites subject to conservation-oriented thinning versus seven sites without thinning. InsectChange only includes data pertaining to saproxylic beetles. The controlled experiment shows a lower decreasing trend of saproxylic beetle abundance with the experimental thinning. The experimental condition therefore biases the insect trend in this dataset towards underestimating their decrease over 12 years. InsectChange includes the sum of saproxylic beetles in experimental (thinning) and control (minimal intervention) sites, and not only the control site with minimal intervention as was inadequately stated in the PlotData paper, while their trends differ and while beetle abundances were separately available for the two types of sites in Table 1 of Gran and Götmark (2019). Moreover, the authors pooled all sites together in a unique plot 1736 but this is not adequate to assess local or landscape drivers, as the seven sites of sampling were up to 250 km apart from Figure 1 in the source publication. For this reason, the estimation of the local cropland cover in InsectChange is incorrect.

**Study 1517 (Guo et al. 2018) - China mosquitos**

This natural experiment monitored changes in mosquito populations during the period of construction of the Three Gorges dam (China), in particular those before and after episodes of water-impoundment and mosquito control policies in two types of habitats, human dwellings and cattle sheds, in three different villages. Four active policies (including pesticide use) were indeed led over the observation in the same time as water impoundment events to control mosquitoes in order to prevent the spread of diseases among humans during the creation of the dam. The dam creation, a highly disturbing factor, is not mentioned in InsectChange. The other studied factors of variation (water impoundment favouring mosquito increases and mosquito control policies favouring their decreases) are not mentioned either. The types of habitats are mentioned in the PlotName cell. Moreover, for Plot 1736 (Zigui, sheds), the abundance number in 1997 (annual total mosquito density) is incorrectly reported from Figure 3 of Guo et al. (2018) as 56.436 in InsectChange, while it is actually much smaller than the density in 1998, reported as 17.822. Finally, the geographic coordinates provided in InsectChange are inappropriate as the original study specifies that data were collected in ten fixed human dwellings and ten cattle sheds in each of the villages of the study, while in each village a unique pair of geographic coordinates was retrieved from Google maps and attributed to the two types of habitat, human dwellings and cattle sheds.

**Study 1518 (Wepprich et al. 2019) - Ohio butterflies**

This observational study monitors the dynamics of butterfly populations in 60 sites in Ohio (US). The common geographic coordinates reported in InsectChange for Plots 1786 and 1798 are very likely inappropriate, as these plots have different locations in the Kitty Todd Nature Preserve in Ohio (U.S.), as indicated by their names ("Kitty Todd Nature Preserve" and "Kitty Todd Nature Preserve-Moseley Barrens"). InsectChange provides no explanation on why it used Google maps rather than the geographic coordinates provided in the original study for the location of these plots. For 15 plots, the local cropland cover provided in InsectChange is overestimated given satellite images.

**Study 1519 (Rochlin et al. 2016) - US mosquitos**

This natural experiment examines the dynamics of mosquito assemblages depending on DDT use, urbanisation and temperature in New York and New Jersey sites of North America. The mosquito densities are particularly correlated to the DDT concentration, recovering after the stop of DDT use, which occurred quite early in the time series, and also with urbanisation, increasing with increasing urbanisation. However, beginning and stop of DDT use, major factors of disturbance of insect communities, are not mentioned in InsectChange. From the source study by Rochlin et al. (2016), Plot 1737 represents 12 trap sites in Suffolk county, New York, and Plot 1738 represents seven trap sites in Ocean County, New Jersey. Therefore, the geographic coordinates provided for these two plots are inadequate for a study at the local or landscape scale.

**Study 1520 (Giraldo-Calderón et al. 2015, Iowa Mosquito Surveillance 2019, Field et al. 2019) - Iowa mosquitos**

The geographic coordinates assigned in InsectChange are very likely inappropriate as for several plots, they are approximative locations found using Google maps. For example, for Plot 1812, the column "SourceGeogrData" of InsectChange table *PlotData.csv* indicates "Google maps, wild guess" as the source for the geographic coordinates indicated in InsectChange, while the name of the plot (Lyme Dump, Linn County, Iowa, USA) is not indicative of a particular place.

**Study 1521 (Petersen et al. 2004) - Denmark springtails**

This study pertains to springtails (Collembola, Entognatha), which are neither insects nor arachnids. This study is a natural experiment exploring long-term changes in these microarthropod communities after the introduction of livestock grazing in abandoned fields with herb–grass vegetation, and comparing grazed versus ungrazed abandoned arable fields. Springtail populations tend to slightly increase in both grazed and ungrazed sites, the unexpected increase in grazed sites was attributed to the population increase of some particular species with adaptation capacities to the new conditions of temperature and humidity of grazed sites. Moreover, there are several errors concerning strata, dates of records, and springtail counts. Only the underground stratum was mentioned in InsectChange, but one third of springtails were collected in the vegetation/litter layer. While there were no springtail records in 1990, InsectChange reported the 1991 springtail records in 1990 and conversely incorrectly reported an absence of springtail record in 1991. On ungrazed sites (Plot 1729), the springtail records of June 1988 (density of 4916.201 per square metre) and November 1988 (density of 23128.492 per square metre) were reversed in InsectChange.

**Study 1524 (Hallmann et al. 2019) - Netherlands light-attracted Lepidoptera and Coleoptera**

This observational study examines the long-term dynamics of insect abundance and biomass from two locations in the Netherlands, nature development area De Kaaistoep and nature reserves near Wijster. In the first location, macro-moths (Lepidoptera), beetles (Coleoptera), caddisflies (Trichoptera), lacewings (Neuroptera), true bugs (Hemiptera-Heteroptera and Hemiptera-Auchenorrhyncha) and mayflies (Ephemeroptera) were monitored using a light trap from 1997 to 2017 for the first two orders and from 2006 to 2017 for the other ones, while in the second location, ground beetles (Carabidae) were monitored using pitfall traps at several sites in the Wijster locality over a longer period extending from 1986 to 2016. The two groups of insects show a declining trend over years but with a sharper slope for ground beetles. InsectChange chose to include light-attracted moths and beetles from the first location (plot 1740) and not ground beetles with a sharper decrease from the second location, while their mean abundance and biomass over all sites were available in Figures 4 and 5b of the source publication by Hallmann et al. (2019), respectively. Moreover, the table *SampleData.csv* inadequately mentioned “insects” in the GroupInData cell, while from Fig. 2 showing 6 light-trapped insect orders from which abundance data were mentioned to be extracted and from our calculations, InsectChange considers the sum of the data for Coleoptera (Figure 2e) and Lepidoptera (macro-moths, Figure 2f) for abundance, the only two groups with a continuous recording over the full period (other groups having available data for the years 2006 and 2009-2017 only). InsectChange also considers the biomass data from Figure 5a, but for Lepidoptera (macro-moth) only. However, this was not indicated in the table *SampleData.csv*, thereby obscuring this inconsistency of selected taxa over metrics. As the biomass data were not available for the Coleoptera at the first location, it would have been more consistent to consider at this location both abundance and biomass for lepidopterans (macro-moths) only. Or the most appropriate selection may have been to focus solely on the abundance of beetles but on both locations and both strata. Finally, the source publication refers to Felix and van Wielink (2008) for a description of the De Kaaistoep site. From Box 1 of this publication, the actual location of the light trap is actually 500 m west from the geographic coordinates assigned to the unique plot of this study in InsectChange. It is difficult to assess the local land cover from satellite images.

**Study 1525 (Giraldo-Calderón et al. 2015) - Indiana mosquitos**

No identified problem.

**Study 1526 (Giraldo-Calderón et al. 2015) - Florida mosquitos**

No identified problem.

**Study 1527 (Rochlin et al. 2016) - US mosquitos**

This natural experiment examines the dynamics of mosquito assemblages depending on DDT use, urbanisation and temperature in a California site of North America. The mosquito densities were shown to increase with the cessation of DDT use and the increasing urbanisation of the site. The beginning and stop of DDT use are not mentioned in InsectChange. From the source study by Rochlin et al. (2016), Plot 1737 represents four trap sites in Sutter-Yuba county, California. Therefore, its geographic coordinates are inadequate for a study at the local or landscape scale.
