## Appendix S2 for "InsectChange: Comment"

**Supporting Information** for "*InsectChange: Comment*". Laurence Gaume, Marion Desquilbet.

### Appendix S2.
### Freshwater studies of the InsectChange database including noninsect taxa

The "insects" considered in the InsectChange database (van Klink et al. 2021) are actually intended to be insects, arachnids and entognaths. Therefore, for the sake of brevity, insects, arachnids and entognaths are hereafter referred to as insects.

**Table S1.** This table specifies the freshwater studies of InsectChange included in Figure 3 (i.e. studies involving non-insects) and shows their distribution according to their degree of information on insect share. Out of the 62 freshwater studies of InsectChange, 28 considered noninsects, among which 24 considered the whole invertebrate assemblage. The study identifiers are those used by Van Klink et al. 2021 and the associated references can also be found in *Problems.xlsx*.

**Table S2**. This table synthesises the qualitative information available on the noninsects, i.e. the different taxa included in the freshwater studies that considered either the whole invertebrate assemblage or a subset of this assemblage. It shows that these studies most frequently include worms, crustaceans and molluscs, especially taxa that are often signs of poor water quality.

**Table S3.** This table credits the photographs used in Figure 3 of the comment to their authors. It specifies the type of noninsect shown in the photograph, the name of the species shown, the name of the author to whom the photograph should be credited, the way the source was used, the copyright license, the date of the photo, additional details and the link where all this can be found.

**Figure S1.** This figure shows the insect share (mean % per plot) and its variation over time (standard deviation per plot) in the time series that have considered the entire invertebrate assemblages instead of only insects and where it was possible to extract the percentage of insects for all or for some records of the time series. Insect shares were often low with high standard deviations over time, meaning that the insect trend cannot be inferred from the trend of the whole invertebrate assemblage

**Table S1** - Distribution of freshwater studies in InsectChange according to their information on the insect share in the invertebrate assemblage

| Invertebrate assemblage considered | Study identifier | # studies |
|---|---|---|
| Whole invertebrate assemblage considered, where insects were not dissociable in any metric or site | 1395, 1423, 1427, 1432, 1437, 1498, 1500, 1503, 1504 | 9 |
| Whole assemblage considered, where insects were reported to be in a minority in the assemblages of all sites | 1454, 1466 | 2 |
| Whole invertebrate assemblage considered, where insects were dissociable in some records in at least one metric or site | 1425, 1452*, 1456*, 1506, 1507, 1509, 1511 | 7 |
| Whole invertebrate assemblage considered, where insects were dissociable over the whole period in at least one metric or site | 1421, 1428, 1435, 1455, 1473, 1513 | 6 |
| Subset of the whole invertebrate assemblage considered, where insects were not dissociable in any site or metric (abundance/biomass) but inferred to be dominant because chironomids represented more than 50% of the quantity of invertebrates. | 1448, 1449, 1451,1457 | 4 |
| Only insects considered | 63, 478, 1347, 1351, 1376, 1381, 1388, 1408, 1412, 1414, 1415, 1417, 1418, 1422, 1426, 1429, 1430, 1431, 1433, 1439, 1440, 1441, 1444, 1453, 1488, 1491, 1499, 1510, 1517, 1519, 1520, 1525, 1526, 1527 | 34 |

Note: Studies with only abundance data are in black, studies with only biomass data are in blue, studies with both abundance and biomass data are in green. *: Studies for which some plots considered the whole assemblage of invertebrates whereas the other plots considered a subset of the assemblage with an unknown insect share.

**Table S2** - Information on non-insect taxa in the freshwater invertebrate assemblages in InsectChange (entirely or partially available in 26 of the 28 studies including non-insects)

| Invertebrates | | | | Molluscs | | Crustaceans | | Worms | | | Tardigrade | Hydrozoans |
|---|---|---|---|---|---|---|---|---|---|---|---|---|
| Study | Metric | Considered assemblage | Information on non-insects | Bivalves | Snails | Amphipods | Others* | Annelids (Oligochaeta /Polychaeta/ Hirudinea) | Flat worms (Turbellaria and others) | Round worms (Nematodes) | | |
| 1421 | A | whole | complete | X | X | XX | XX | X | XX | - | - | X |
| 1423 | A | whole | partial | NA | NA | NA | NA | X | NA | NA | NA | NA |
| 1425 | AB | whole | complete for some records | XX(invasive) | XX | - | X | X | X | X | - | X |
| 1427 | AB | whole | complete | X | X | X | X | X | - | X | - | X |
| 1432 | B | whole | partial | X | NA | NA | X | X | NA | X | NA | NA |
| 1435 | A | whole | complete | X | XX | XX | XX | XX | X | - | - | - |
| 1437 | B | whole | partial | NA | NA | NA | NA | X | NA | X | NA | NA |
| 1448 | AB | subset | complete | NA | NA | NA | NA | X | NA | NA | NA | X |
| 1449 | AB | subset | complete | NA | NA | NA | X | X | NA | X | X | X |
| 1451 | B | subset | partial | NA | NA | NA | X(invasive) | NA | NA | NA | NA | NA |
| 1452 | B | whole 2 plots subset 6 plots | complete for some records | XX(invasive) | X | X | X | X | - | - | - | - |
| 1454 | B | whole | reported as a majority | NA | NA | NA | X | X | NA | XX | NA | NA |
| 1455 | AB | whole | complete | XX | - | X | XX | XX | X | X | X | X |
| 1456 | AB | whole 3 plots subset 1 plot | complete for mean 3 plots | X(invasive) | XX(invasive) | X (invasive) | X | X(invasive) | - | X | - | - |
| 1457 | AB | subset | partial | NA | NA | NA | NA | NA | NA | X | NA | X |
| 1466 | AB | whole | reported as a majority | XX(invasive) | NA | X | X | X | NA | NA | NA | NA |
| 1473 | AB | whole | complete | X | X | X | X | X | X | X | - | - |
| 1498 | A | whole | partial | NA | X | X | X | X | NA | NA | NA | NA |
| 1500 | A | whole | partial | NA | NA | NA | NA | X | X | NA | NA | NA |

Table S2 - continued

| **Invertebrates** | | | | **Molluscs** | | **Crustaceans** | | **Worms** | | | **Tardigrade** | **Hydrozoans** |
|---|---|---|---|---|---|---|---|---|---|---|---|---|
| **Study** | **Metric** | **Considered assemblage** | **Information on non-insects** | **Bivalves** | **Snails** | **Amphipods** | **Others*** | **Annelids (Oligochaeta /Polychaeta/ Hirudinea)** | **Flat worms (Turbellaria and others)** | **Round worms (Nematodes)** | | |
| 1503 | A | whole | partial | NA | X | X | X | X | X | X | NA | X |
| 1504 | A | whole | partial | NA | X | NA | X | X | X | NA | NA | NA |
| 1506 | AB | whole | partial, overall % known | NA | NA | NA | X | X | NA | X | NA | NA |
| 1507 | AB | whole | complete for some records | - | X | XX | - | X | X | X | - | - |
| 1509 | AB | whole | partial | X | X | XX | X | XX | NA | NA | NA | NA |
| 1511 | B | whole | complete for some records | X | X | - | X | X | - | - | - | - |
| 1513 | A | whole | complete | - | - | - | XX | XX | X | XX | - | - |
| 1395 | A | whole | none | NA | NA | NA | NA | NA | NA | NA | NA | NA |
| 1428 | AB | whole | none | NA | NA | NA | NA | NA | NA | NA | NA | NA |
| **Total number of studies** | | | **28** | | | | | | | | | |
| **# studies with information** | | | **26** | **12** | **13** | **12** | **20** | **24** | **10** | **14** | **2** | **8** |
| **% Studies including the subgroup** | | | | **46.2%** | **50%** | **46.2%** | **76.9%** | **92.3%** | **38.5%** | **53.8%** | **7.7%** | **30.8%** |
| **# studies with information** | | | **26** | **16** | | **20** | | **24** | | | **2** | **8** |
| **% Studies including the group** | | | | **61.5%** | | **76.9%** | | **92.3%** | | | **7.7%** | **30.8%** |

Note: The references of the studies are available in *Problems.xlsx*. A: study with abundance data, B: study with biomass data, AB: study with abundance and biomass data, X: invertebrate group reported to be present in the assemblage, XX: invertebrate group observed to be particularly abundant in the assemblage, -: invertebrate group absent or not mentioned, Others*: other crustacean types among shrimp, crayfishes, crabs, isopods, copepods, cyclopods, ostracods, cladocerans… NA: no available information. Information on noninsect taxa involved was extracted from the studies themselves except for study 1427 where information on noninsects was available in Baxter and Minshall (2016). When in a study some plots considered the whole invertebrate assemblage whereas other plots considered a subset of this assemblage involving noninsects, we show the noninsects that were present in the whole assemblage.

**Table S3 -** Credits for the photographs used in Figure 3 and information on their sources

| **Non-insect type (Figure 3)** | **Invertebrate species name** | **Source (Author of the photo)** | **Use of the source** | **Copyright license** | **Date** | **Details / Link** |
|---|---|---|---|---|---|---|
| Mussels | *Dreissena polymorpha* - Moule zebrée (zebra mussels) | F. Lamiot | Modified (focus on some mussels only) | CC BY-SA 1.0 | Oct.2006 | Zebra mussel, an invasive freshwater species, photographed in the canalised deule at Lambersart, near Lille (Northern France, Europe), in 2006.<br>https://commons.wikimedia.org/w/index.php?curid=1316814 |
| Snails | *Planorbis planorbis* (Linnæus, 1758) | H. Zell | Modified (focus on the bottom left specimen) | CC BY-SA 1.0 | Oct.2019 | Diameter 1.0 cm; Originating from Rappenwörth, Karlsruhe, Germany; Shell of own collection, therefore not geocoded.<br>https://commons.wikimedia.org/w/index.php?curid=1316814 |
| Crayfishes | *Austropotamobius pallipes* (white-clawed crayfish) | D. Geke | Not modified | CC BY-SA 3.0 | Dec.2007 | https://fr.wikipedia.org/wiki/%C3%89crevisse_%C3%A0_pattes_blanches#/media/Fichier:Austropotamobius_pallipes.jpg |
| Shrimps | *Nototropis swammerdamei* (Milne-Edwards, 1830) - Amphipoda | H. Hillewaert | Not modified | CC BY-SA 3.0 | Oct.1999 | Belgian Continental Shelf. Camera mounted on a Zeiss Stemi C-2000 binocular microscope. Length: ~4 mm. Geo-location not applicable as the picture was taken in the lab. Note the typically deeply pleated coxal gills.<br>https://commons.wikimedia.org/wiki/File:Nototropis_swammerdamei.j pg |
| Leeches | *Erpobdella octoculata* (Hirudinea) | W. Walas | Not modified | CC BY-SA 3.0 | Jul.2012 | https://commons.wikimedia.org/w/index.php?curid=20294374 |
| Worms | *Tubifex tubifex* (Sludge worm) | J.Reischig | Not modified | CC BY-SA 3.0 | Mar.2014 | Total preparation. Optical microscopy technique: Negative phase contrast. Magnification: 120x (for picture width 26 cm ~ A4 format).<br>https://commons.wikimedia.org/wiki/File:Sludge_worm_(265_31)_Total_preparation.jpg |

**Figure S1 -** Insect share (mean percentage) in time series included in InsectChange relating to whole invertebrate assemblages and where it was possible to calculate the insect share and its variation over time (standard deviation), (A) for all time records for 37 plots from 6 studies or (B) for some available time records for 11 plots from 7 studies.

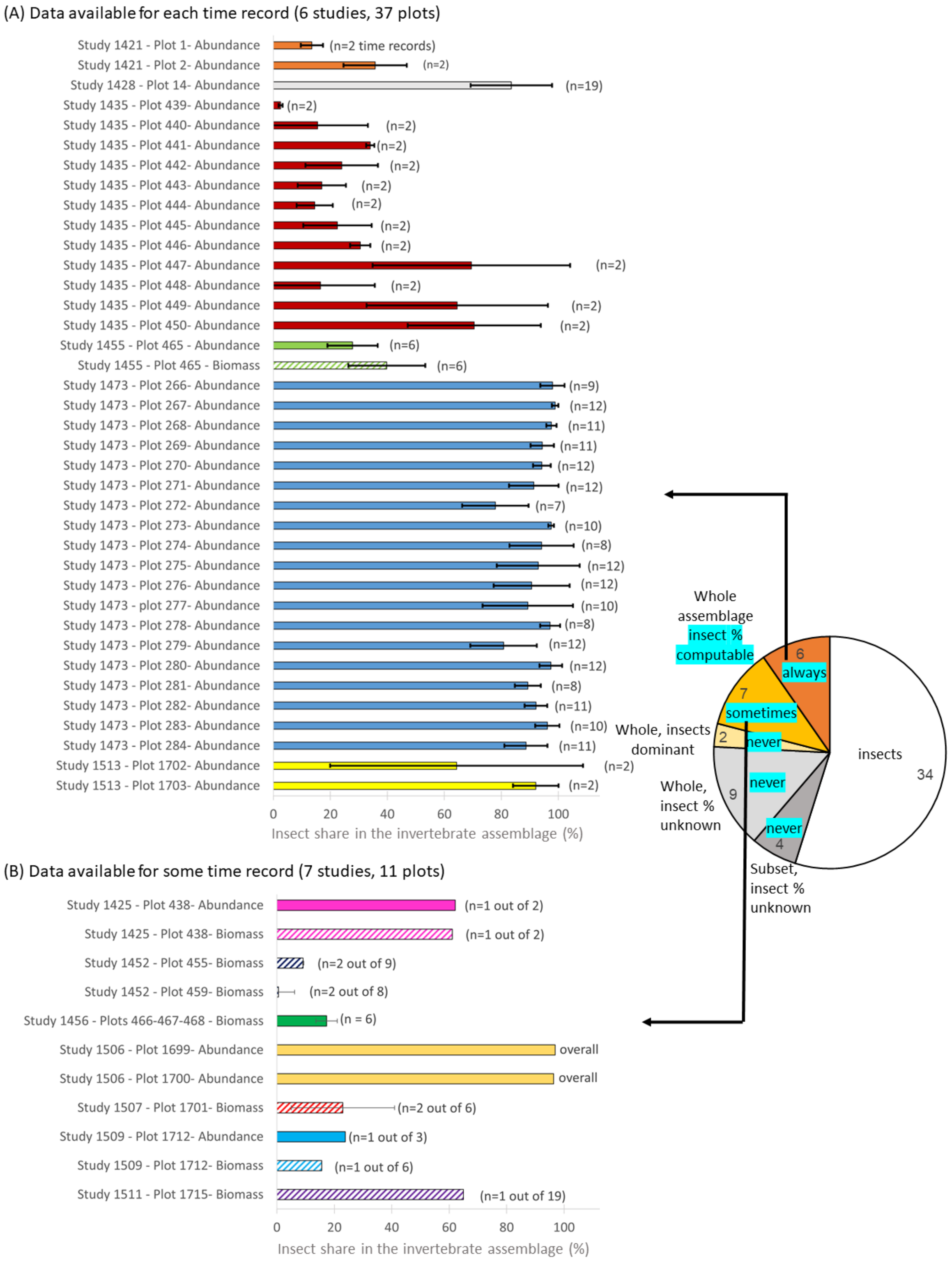


Note: One colour per study. Solid bars: abundance data; hatched bars: biomass data. The pie chart is that shown in Figure 3 of the comment. For study 1506, only the mean insect share over time was available.

**References.**

Baxter, C.V. and G.W. Minshall. 2016. Invertebrate abundance and biomass data from Big Creek tributaries (1988-2012). Fort Collins, CO: Forest Service Research Data Archive. https://doi.org/10.2737/RDS-2016-0027

van Klink, R., D. E. Bowler, O. Comay, M. M. Driessen, S. K. M. Ernest, A. Gentile, F. Gilbert, et al. 2021. InsectChange: a global database of temporal changes in insect and arachnid assemblages. Ecology **102**:e03354.
