## Appendix S3 for "InsectChange: Comment"

**Supporting Information** for "*InsectChange: Comment*". Laurence Gaume, Marion Desquilbet.

### Appendix S3.

### Expected trends in the studies of InsectChange affected by internal drivers

We classified the studies of the InsectChange database (van Klink et al. 2021) affected by internal drivers according to three categories:

- **Controlled experiments**, when the focus of the study was a particular factor/treatment studied through experimental variations of this factor/treatment across space and/or time;
- **Natural experiments**, when the focus of the study was a particular factor/disturbance studied through natural variations of this factor/disturbance across space and/or time, or exclusively across time in interrupted time series where a qualitative factor/disturbance varied during the time record;
- **Studies with a major disturbance** (affecting all or some sites).

We did not include studies where the only factor of interest was linked to climate change, given that climate change is global and thus the factor may not be specific to the study.

We then qualified in each study the habitat change resulting from the internal driver of the study (in terms of habitat creation, natural restoration, remediation/restoration, modification or degradation) and the subsequently expected insect change (colonisation, recovery, proliferation, decline) in the Table below. By expected changes, we mean changes that were expected under the hypotheses tested in the source studies or logical changes anticipated to arise given the particular situations caused by the internal drivers and/or the taxa selection by InsectChange from the source studies (e.g. proliferation of stress-tolerant, sometimes non-insect, taxa, selected in InsectChange in freshwaters described by the source study as undergoing eutrophication). Although in most cases these expected or anticipated changes were indeed observed in the study, it was not systematically the case. For example, Stout and Rondellini (1995) investigated the impacts of low frequency electromagnetic fields on freshwater assemblages from electromagnetic fields and observed no effect (study 1417).

Our analysis shows that out of the 88 studies with an experimental design or a major disturbance, the invertebrates considered were expected to increase in 42 studies, to decrease in 9 studies, to be variable in 35 studies or there was no expectation in 2 studies.


Synthesised table shown next page

**Table S1.** Expected trends in experimental studies or studies with a major disturbance

| Habitat change | Expected invertebrate change | No of datasets | Specific condition (Dataset) |
|---|---|---|---|
| **a) Expected increase** | | **42** | |
| Habitat creation | colonisation | 8 | artificial ponds (63)<br>polder (1324)<br>artificial nesting sites (1397)<br>reservoirs (1449, 1451, 1452, 1456)<br>streams fed by a melting glacier (1513) |
| Habitat natural restoration | recovery | 9 | initial forest fragmentation (1385)<br>initial fire (1387, 1437, 1479)<br>initial drought (1388)<br>initial flood (1393)<br>initial windfall (1407)<br>initial cold thermal regime of a dam (1421)<br>initial hurricane (1487) |
| Habitat remediation or restoration | recovery | 18 | cessation of agriculture (313, 1410, 1411, 1425)<br>cessation of industrial activity (1398, 1419, 1499)<br>cessation of DDT use (1519, 1527)<br>measures of pollution abatement (1422, 1423, 1473, 1508)<br>measure of desalination (1502)<br>drainage of acid mine (1504)<br>revegetalisation of a rubble dump (1396)<br>thermal restoration (1430)<br>conservation-oriented thinning (1516) |
| Habitat modification | proliferation | 1 | proliferation of invasive clams included in the considered assemblage (1466) |
| Habitat degradation | proliferation of stress-tolerant taxa (focus of the study or largely dominant in the assemblage) | 6 | proliferation of chironomids following water eutrophication (1431, 1439, 1440, 1453)<br>proliferation of amphipods/oligochaetes included in the considered assemblage (1507, 1509) |
| **b) Expected decrease** | | **9** | |
| Habitat degradation | decline | 9 | habitat loss/fragmentation<br>due to infrastructure development or agriculture (294, 1434)<br>pesticide use (478, 1395)<br>pollution/industrialisation (1376, 1413, 1457)<br>electromagnetic field (1417)<br>competitive invasive species (1441) |
| **c) Variable expected change / no expectation** | | **37** | |

| | | | |
|---|---|---|---|
| Habitat modification | no expectation relative to the assemblage, only to specific species | 1 | Abandonment of pasture and overgrowing with shrubs (1392) |
| Habitat modification | depend on competition or facilitation | 1 | Controlled exclusion of interacting species (1346) |
| Occasional degradation(s) / improvement(s) of habitat by major natural events | decline then recovery / increase then decline | 8 | oscillations according to flood and post-flood events (1351, 1428)<br>oscillations according to changes in primary producers and water quality (1381)<br>oscillations according to high rainfalls (1382)<br>middle-time catastrophic fires (1319, 1406, 1427)<br>middle-time catastrophic flood (1498) |
| Habitat modification | depends on modalities of factor studied | 15 | agricultural practices (300, 1460, 1474)<br>land uses (1312, 1335, 1340)<br>moisture (1458)<br>managed plant diversity (1364)<br>elevation (1367, 1391, 1465, 1480)<br>habitat types (1435)<br>wind disturbance in atmosphere (1493)<br>null, negative or positive land disturbance (1102) |
| Habitat modification, remediation, restoration, degradation | depends on outcomes of multiple factors studied | 12 | experimental fires and grazing (301)<br>experimental tree treatments simulating death by phytophages/preventive salvage operations (1261)<br>tree species and experimental treatments simulating hurricane (1357)<br>local undisturbed land covers, surrounding urbanisation (1365)<br>habitat connectivity, in-stream vegetation and extreme climatic events (1408)<br>land use and habitat type (1409)<br>radioactive contamination and cessation of human activity (1464)<br>land cover and pollarding (1497)<br>salinity mitigation, floods and droughts (1503)<br>phosphorus enrichment and increase in flood events (all invertebrates) (1506)<br>water impoundment and mosquito-control policies (1517)<br>agriculture abandonment and livestock grazing (1521) |

Note: study numbers in brackets, terrestrial studies in red, freshwater studies in blue.
